# Coordination between *E. coli* Cell Size and Cell Cycle Mediated by DnaA

**DOI:** 10.1101/2020.02.14.949446

**Authors:** Qing Zhang, Zhichao Zhang, Hualin Shi

## Abstract

Sixty years ago, bacterial cell size was found as an exponential function of growth rate. Fifty years ago, a more general relationship was proposed, in which the cell mass was equal to the initiation mass multiplied by the ratio of the total time of the C and D periods to the doubling time. This relationship has recently been experimentally confirmed by perturbing doubling time, C period, D period or the initiation mass. However, the underlying molecular mechanism remains unclear. Here, we developed a mechanistic and kinetic model to describe how the initiator protein DnaA mediates the initiation of DNA replication in *E. coli.* In the model, we introduced an initiation probability function involving competitive binding of DnaA-ATP (active) and DnaA-ADP (inactive) at replication origin to determine the initiation of replication. In addition, we considered RNAP availability, ppGpp inhibition, DnaA autorepression, DnaA titration by chromosomal sites, hydrolysis of DnaA-ATP along with DNA replication, reactivation of DnaA-ADP and established a kinetic description of these DnaA regulatory processes. We simulated DnaA kinetics and obtained a self-consistent cell size and a regular DnaA oscillation coordinated with the cell cycle at steady state. The relationship between the cell size obtained by the simulation and the growth rate, C period, D period or initiation mass reproduces the results of the experiment. This model also predicts how the number of DnaA and the initiation mass vary with the perturbation parameters (including those reflecting the mutation or interference of DnaA regulatory processes), which is comparable to experimental data. The results suggest that the regulatory mechanisms of DnaA level and activity are associated with the invariance of initiation mass and the cell size general relationship for matching frequencies of replication initiation and cell division. This study may provide clues for concerted control of cell size and cell cycle in synthetic biology.

## I. INTRODUCTION

In the bacterial cell cycle, DNA replication, cell division, and cell growth are coordinated with each other [1]. Cell cycle events have featured timings. Bacteria cell size increases over the entire interdivision cycle [2–6], while the chromosome is duplicated in the “C period” (from the initiation to the termination), and the time between replication termination and cell division is called the “D period” [7]. The relationship between cell size and other cell cycle parameters (e.g., doubling time, C period, and D period) has been quantitatively studied by various experiments and phenomenological models [7–12]. Schaechter *et al.* [8] discovered an exponential relationship between cell mass and growth rate of *S. typhimurium* under various nutrient conditions, which is known as the nutrient growth law [11]. Cooper and Helmstetter [7] found that when the doubling time was 20-40 minutes, C and D periods of *E.coli* were approximately constant (40 min and 20 min, respectively). When the doubling time is shorter than *C* + *D*, it is impossible to perform a round of DNA replication in an interdivision cycle. In order to solve this problem, Cooper and Helmstetter [7] further proposed that there is a gap between two adjacent runs of DNA synthesis in slow growth and multiple replication forks exist in fast growth, based on their direct measurement on DNA synthesis rate [13]. The Cooper-Helmstetter model phenomenologically answers how DNA replication is coupled to cell division cycle at different growth rates. Combining the Cooper-Helmstetter model and the nutrient growth law, Donachie [9] proposed that the ratio of cell mass (*M*_*i*_) to origin number (*N*_*i*_) at replication initiation (*M*_*i*_/*N*_*i*_) (termed initiation mass) is invariant at different growth rates and cell mass at division (*M*_*d*_) can be expressed as an exponential function of (*C* + *D*)*/τ* by

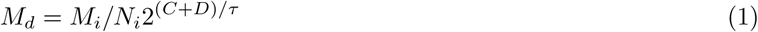

where *τ* denotes the doubling time. Since *M*_*d*_ is proportional to the average cell mass (size) and *C* + *D* is constant according to the Cooper-Helmstetter model, Eq. 1 captures the exponential relationship between cell size and the growth rate, i.e. the nutrient growth law. In addition to the experiment of Schaechter *et al.* [8] for *S. typhimurium*, many other experiments supported the increase in cell size with growth rate and the (approximate) invariance of initiation mass in other bacterial organisms, e.g., *E.coli, B. subtilis*, and *S. meliloti* [11, 14–19]. The concepts of the Cooper-Helmstetter and Donachie models are widely accepted for its simplicity, although they may not hold precisely under some actual experimental conditions [15, 20, 21]. By incorporating or respecting the Cooper-Helmstetter and Donachie models, a number of theoretical models involving the control of DNA replication initiation have been constructed (e.g. [18, 22–29]).

Recent experimental data indicated that the Donachie relation still holds under various perturbations that do not affect nutrient quality, which greatly extends the nutrient growth law [10–12]. In particular, Si *et al.* [11] proposed a general growth law based on experiments specifically perturbing growth rate, C period, D period, initiation mass, and etc., that is, cell size (*S*) is the sum of unit cells (*S*_0_), i.e.

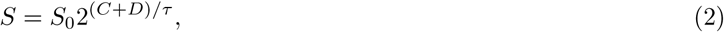

where *S*_0_, as the ratio of the cell size to the origin number, is proportional to the initiation mass (used interchangeably with initiation mass in the following). The general growth law phenomenologically relates cell size to cell cycle timings (*τ, C*, and *D*) and initiation mass (*S*_0_). From Eq. 2, a specific perturbation of *τ, C, D*, or *S*_0_ (while other parameters remain unchanged) gives a particular relationship between cell size and the perturbed parameter, corresponding to a particular growth law. However, the regulatory circuits controlling these individual growth laws are poorly understood.

The initiator protein DnaA can bind to the vicinity of the chromosomal replication origin (*oriC*) and form a DnaA-ATP oligomer (∼20 DnaA-ATP monomers) which is responsible for opening the duplex unwinding element (DUE) of *oriC* [30, 31]. Many previous experiments (reviewed in [31–33]) and models (e.g. [22, 24–26, 34, 35]) have shown that the regulation on DnaA controls the initiation of DNA replication and mediates the coupling of DNA replication to cell division. In this study, we aim to investigate the role of the regulation on DnaA in harmonizing cell size and cell cycle timings.

The availability of active DnaA (DnaA-ATP) for initiating *oriC* is regulated in a number of ways, including autorepression at transcription initiation, titration by non-functional DnaA-boxes, inactivation with replication fork, reactivation with special DNA sites or membrane, and competition between DnaA-ATP and DnaA-ADP (see details below or [26]). Many models of DnaA controlling the initiation of DNA replication have been proposed, e.g. the replicon model [36], autorepressor model [37], initiator titration model [24], and competition model [26, 34]. In these models, different regulation details on DnaA are included, and different variables related to the DnaA level are used to determine if *oriC* is initiated (equivalently). Variables used for setting the initiation condition include DnaA concentration [36, 37], free DnaA-ATP concentration [22, 25], the DnaA/DnaA-box ratio[24], the DnaA-ATP/genome content ratio [35] and the DnaA-ATP/DnaA-ADP ratio [34].

Donachie and Blakely [34] proposed a qualitative model of initiator competition in which the competition between DnaA-ATP and DnaA-ADP in binding to *oriC* determines the initiation of DNA replication. With respect to the competition between DnaA-ATP and DnaA-ADP in binding to *oriC*, we previously proposed an initiation probability function to set the initiation condition and developed a mathematical model to show that the regulation of DnaA oscillation is able to coordinate DNA replication with cell cycle at different growth rates [26]. However, both Cooper-Helmstetter and Donachie models were used in advance in this previous model.

Based on the previous model [26] and up-to-date empirical data [10–12], we have developed a novel competitive initiation model here. This model is used to resolve how the regulation of DnaA dynamics determines the initiation of DNA replication and couple DNA replication to cell division when perturbing growth rate, C period, D period, or initiation mass in *E. coli.* In the model, cell size was an output variable and its variation with the perturbed parameter was obtained by simulation. The results show that the relationship between cell size and the disturbed parameter follows the general growth law under various perturbations of growth rate, C period, D period, or initiation mass. This suggests that changes in cell size help regulate DnaA kinetics, thereby regulating or maintaining the initiation frequency of DNA replication to match the growth rate. Our model also predicts regular oscillations of each form of DnaA that are harmonized with other cell cycle events, and the average number of each form of DnaA as a function of the perturbed parameter. In particular, the results support that ppGpp may be a major global regulator that inhibits the initiation of DNA replication to couple with the reduced growth rate. Current experimental evidence suggests that ppGpp’s inhibition of replication initiation can be achieved through three possible indirect mechanisms (see the description of the model). Our model shows that these three mechanisms lead to different dependences of DnaA number (concentration) and DnaA-ATP/DnaA-ADP ratio on growth rate, indicating that the dominant mechanism could be determined by measuring the growth rate-dependence of DnaA. Our model also predicts the quantitative effects of some mutations/interferences of DnaA regulatory processes (for example, DnaA knockdown and *oriC* block) on DnaA level and activity and initiation mass that are comparable to the experiment, whereas the phenomenological models cannot. We also show that, at least in some cases, our initiation condition cannot be replaced with others. In addition, we discussed the effect of changing each regulatory process of DnaA and pointed out which regulatory processes are crucial for maintaining a regular cell cycle under perturbed conditions.

## II. MATERIALS AND METHODS

### A. A theoretical model of DnaA mediating DNA replication initiation to couple to cell cycle

Here we constructed a kinetic model of DnaA-mediated DNA replication initiation by adapting the previous model [26]. First, we used the same initiation condition as previously [26]. At *E.coli oriC*, it has been identified three widely spaced similarly strong boxes (R1, R2, and R4) and more than five weak boxes (e.g., R3, R5M, I1, I2, I3) located between the strong ones [38–41] (Fig. 1 A). At the initiation, strong boxes bind DnaA-ATP molecules as anchors, and then weak boxes bind DnaA-ATP gradually with the cooperation of adjacent bound DnaA-ATP; finally, more than ten DnaA-ATPs form an ordered polymeric pre-replication complex (pre-RC), which opens DUE of *oriC* [39–41]. To model the initiation process, we coarse-grained wild-type *oriC* of *E.coli* [40] as one DUE neighbored with three identically strong and four identically weak DnaA boxes alternatively interspersed (the same for [26])(Fig. 1 A). One strong box binds DnaA-ATP independently, while one weak box binds DnaA-ATP only if its adjacent strong box has bound a DnaA-ATP. For a single cell, the initiation happens (DUE is opened) when all seven DnaA boxes bind DnaA-ATP molecules. Given concentrations of free DnaA-ATP and free DnaA-ADP (denoted by *c*_1_ and *c*_2_, respectively), we calculated the probability of all seven boxes binding DnaA-ATP in a cell. For a population average cell, we assume that *oriC* is initiated when this probability is equal to or higher than a threshold and thus this probability is called the “initiation probability” hereinafter. Based on the above assumptions, the initiation probability can be derived as (see details in Supporting Text or Ref. [26])

**FIG 1.**
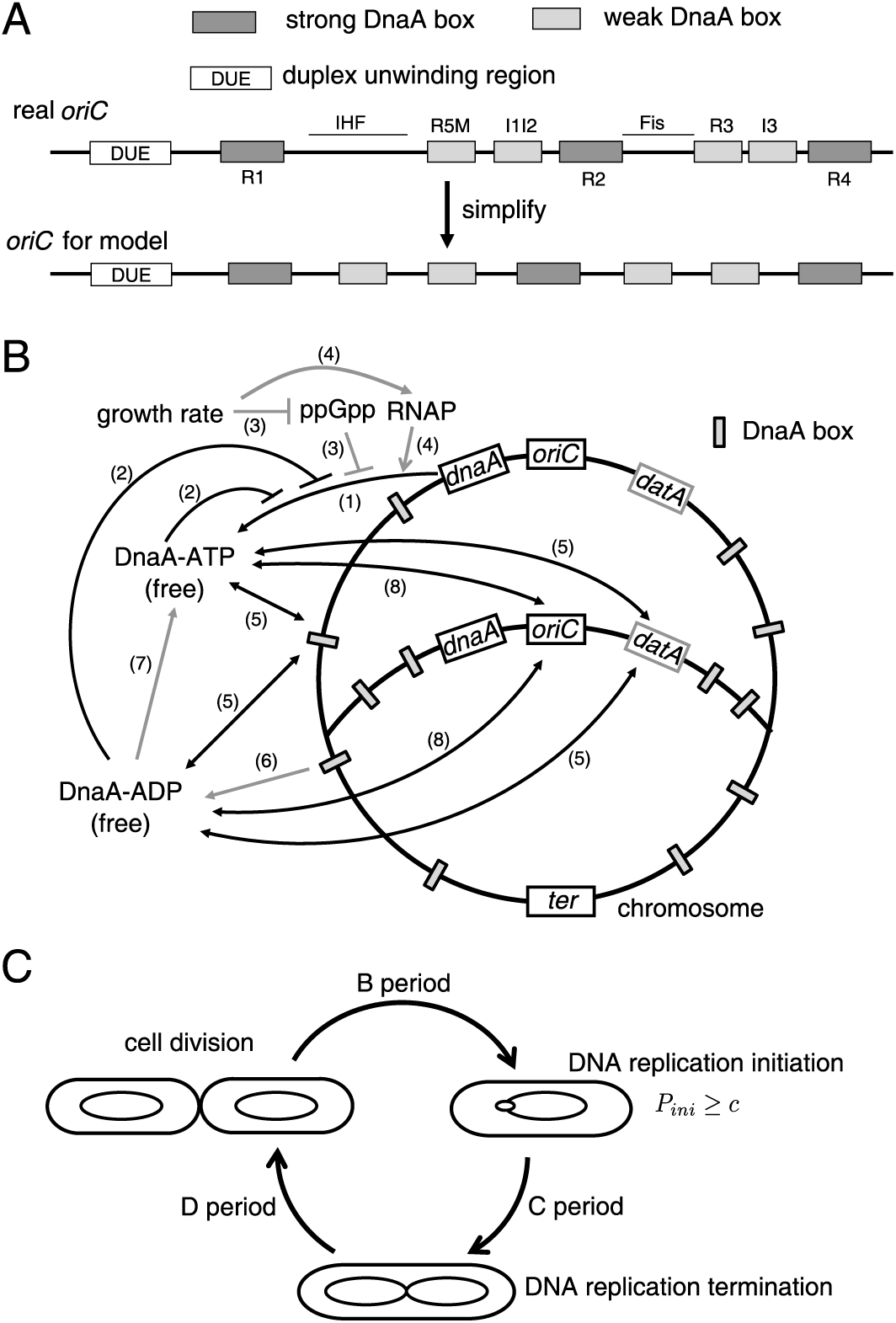
Schematic illustration for the DnaA-mediated replication initiation model. (A) Structure characteristics of *E.coli oriC* (adapted from [26]). In real *oriC* ([40]), DnaA boxes, Fis and IHF sites, and the duplex unwinding region (DUE) are indicated. Fis and IHF regulate DnaA binding to their boxes [40], as the background of the formation of DnaA-ATP oligomer at *oriC.* The simplified *oriC* for the model includes three identical strong DnaA boxes (corresponding to R1, R2, and R4) and four identical weak ones (corresponding to R5M, I1I2, R3, and I3). DnaA-ATP competes with DnaA-ADP for binding to strong boxes and DnaA-ATP bound to strong box enhances largely the binding of DnaA-ATP to the adjacent weak box. (B) Regulatory mechanisms for DnaA (adapted from [22, 25, 26]): (1) Nascent DnaA binds with ATP. (2) *dnaA* transcription is auto-repressed by DnaA-ATP and DnaA-ADP. (3) (p)ppGpp, whose concentration decreases with growth rate, inhibits *dnaA* transcription. (4) The availability of RNA polymerase (RNAP), increasing with growth rate, affects *dnaA* transcriptional level positively. (5) DnaA-ATP and DnaA-ADP are titrated by nonfunctional DnaA boxes on the chromosome. (6) DnaA-ATP on the chromosome is hydrolyzed into free DnaA-ADP along with DNA replication. (7) Free DnaA-ADP is reactivated into free DnaA-ATP. (8) Free DnaA-ATP competes with free DnaA-ADP for binding to *oriC.* The processes shown by grey lines were considered differently from previous models ([26]). (C) DnaA kinetics determines the initiation of DNA replication, which is coupled with other cell cycle events. When the initiation probability *P*_*ini*_ reaches the threshold (*c*), DNA replication is initiated and after a time of *C* (i.e. C period), it is terminated. The cell is divided after a time of *D* from replication termination (i.e. D period). Note that when the growth rate is high enough, replication initiation occurs one or two cell cycles in advance, leading to the disappearance of B period (the time between cell birth and replication initiation) and the existence of multiple (more than two) replication forks. Moreover, cell size increases exponentially with the time during the cell cycle as shown in Eq. 4.

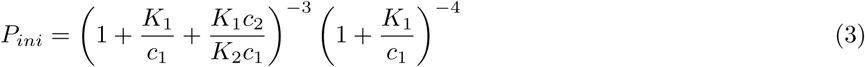

where *K*_1_ and *K*_2_ denote dissociation constants of DnaA-ATP from DnaA boxes and of DnaA-ADP from strong DnaA boxes. When *c*_2_ ≪ *K*_2_, one has 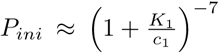, which indicates that free DnaA-ATP concentration determines the initiation, corresponding to the noncompetition initiation. If *c*_1_ ≫ *K*_1_ and *c*_2_ ≫ *K*_2_, it yields 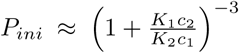, which shows that the initiation is determined by the ratio of DnaA-ATP to DnaA-ADP, belonging to the full competition initiation.

There is a minimal interval between two adjacent initiations, so-called the “eclipse” period [42]. Right after one initiation of chromosome replication, SeqA binds to hemimethylated GATC sites at *oriC* to repress re-initiation of *oriC* until these GATC sites are fully methylated by Dam, which is the major source of the eclipse period [43–45]. In our model, therefore, we set that the initiation is permitted to happen after the eclipse period (*T*_*e*_). Typically, *T*_*e*_ has an order of 10 − 20 min [24, 42, 46]. Decreased replication elongation rate may increase *T*_*e*_ by slowing methylation of *oriC.* It was also reported that *T*_*e*_ is around 60% of the doubling time [47]. We tested three cases: a constant *T*_*e*_ (typically 10 min), *T*_*e*_ = 0.25*C*, and *T*_*e*_ = 0.6*τ*, and there was no difference in results. It indicates that *T*_*e*_ has little effect on replication initiation frequency, in agreement with previous models [24, 26].

Second, we adopted that cell volume (*V*) increases with the time (*t*) as

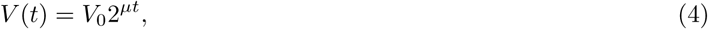

where *μ* denotes the growth (doubling) rate (i.e. 1*/τ*) and *V*_0_, birth cell size. Notice that the definition of growth rate here lacks a scaling factor ln 2, compared with another common definition of growth rate by ln 2*/τ*, and correspondingly, the unit of growth rate in our definition is doublings/hour (abbreviated as dbls/h) rather than 1/hour. The growth pattern of individual bacterial cells has been described by exponential, linear, bilinear and other forms, while currently available empirical data is more likely to support the exponential model [2–6].

Third, we formulated the dynamics of DnaA synthesis, transformation, and translocation based on the Ref. [26] (Fig. 1 B). Let *x, y, x*_*f*_, *y*_*f*_, *x*_*b*_ and *y*_*b*_ denote numbers of all DnaA-ATP, all DnaA-ADP, free DnaA-ATP, free DnaA-ADP, DNA-bound DnaA-ATP, and DNA-bound DnaA-ADP in a cell, respectively. The concentrations of free DnaA-ATP and free DnaA-ADP can be represented by *c*_1_ = *x*_*f*_/*V* and *c*_2_ = *x*_*f*_/*V*.

DnaA promoters (i.e., *dnaAp1* and *dnaAp2*) are autorepressed by DnaA [48] and suppressed in the stringent response [49]. Their activity decreases in a similar manner with the growth rate, and regardless of the growth rate, the activity of *dnaAp2* is significantly higher than that of *dnaAp1* [49, 50]. The growth rate-dependent regulation of DnaA promoters is mainly at the transcription level [49]. To determine the synthesis rate of DnaA (denoted as *ρ*(*t*)), for simplicity, we only consider DnaA expression from *dnaAp2*). Corresponding regulators (mainly working at transcription initiation) include: (1) The availability of RNAP determines the increase in the transcription from a constitutive promoter with the growth rate [51–53]. The previous model has suggested that free RNAP concentration may be a candidate for coupling DNA replication initiation to growth rate by affecting DnaA transcription [26].

According to the difference between its sequence and the consensus sequence and its activity measurement [50, 54] *dnaA* promoter is weak, so it is reasonable to assume that *ρ*(*t*) is proportional to the concentration of free RNA polymerase (RNAP) ([*RNAP*]_*f*_) [26]. (2) Gene dosage is another factor that contributes to the growth rate dependence of gene expression [55]. *dnaA* is close to *oriC* on the chromosome [54], so the copy number of *dnaA* (*n*_*dnaA*_(*t*)) is approximately equal to that of *oriC* (*n*_*ori*_(*t*)), that is, *n*_*dnaA*_(*t*) ∝ *n*_*ori*_(*t*). (3) *dnaA* promoter is autorepressed by DnaA-ATP and DnaA-ADP [48]. A formula consistent with experimental data has been proposed to quantitatively describe the autorepression [26]. (4) The stringent response alarmone (p)ppGpp is a major regulator controlling the growth rate [56–58]. The initiation of DNA replication is inhibited in the stringent response [49, 59–61], by ppGpp repressing the *dnaA* expression [49, 50, 60]. Combining the effects of all these factors, we obtain the DnaA synthesis rate (see derivation details in the Supporting Text)

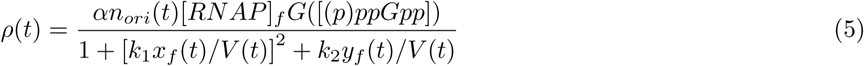

where *k*_1_(= 0.024*μm*^3^) and *k*_2_(= 0.0135*μm*^3^) are binding rate constants of DnaA-ATP and DnaA-ADP to the *dnaA* promoter (fitted with empirical data of Speck *et al.* [48]), the power 2 reflects the self-cooperation of DnaA-ATP in binding to the promoter, and *α* is the synthesis rate constant. Compared with the previous model ([26]), we newly introduced *G*([*ppGpp*]) as a decreasing function of ppGpp concentration to reflect the effect of ppGpp inhibition on DnaA expression. When *G*([*ppGpp*]) = 1, formula 5 degenerates into the formula used in Ref. [26]. Since the details of ppGpp inhibition of DnaA expression are unknown, we use a Hill function, i.e.

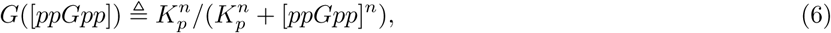

to roughly represent the ppGpp effect, where *K*_*p*_ and *n* denote equilibrium constant and Hill coefficient, respectively, which are fixed by fitting experimental data (see Fig. 2). Note that (1) Equation 6 mainly reflects the inhibitory effect of ppGpp at the transcriptional level [49], and it may also reflect the inhibitory effect of ppGpp at the translational level, which reduces the active ribosome fraction at small growth rates [58]. (2) There are two other possible mechanisms of ppGpp inhibition that do not directly affect DnaA synthesis but mediate the coupling between replication initiation and growth rate. To simulate these two possible mechanisms, we reset *G*([*ppGpp*]) to be a constant (see below). Current experimental data are still insufficient to clarify which mechanism plays a dominant role. For simplicity, we mainly show the results for the mechanism of ppGpp inhibiting DnaA synthesis. The comparison between simulations of these three mechanisms is present in the Discussion. (3) Under a perturbation that affect the growth rate little, we also replace *G*([*ppGpp*]) with a constant. DnaA binds tightly to ATP and ADP, and has a higher affinity for ATP than ADP [62]. In addition, ATP is much more than ADP in cells, therefore, newly synthesized DnaA is more likely to become DnaA-ATP [48]. Therefore, *ρ*(*t*) actually indicates the production rate of DnaA-ATP.

**FIG 2.**
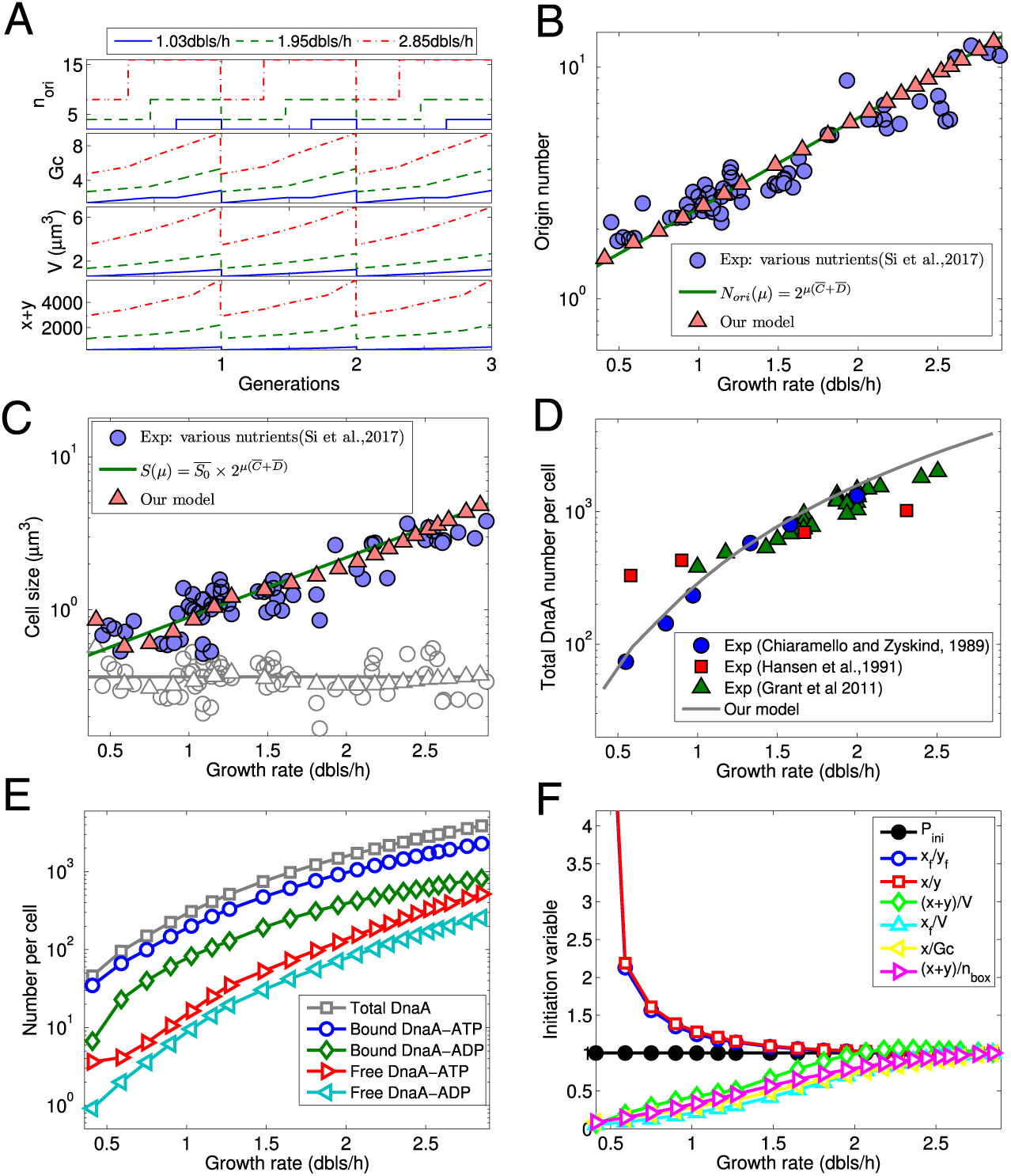
Our model quantifies origin number, genome content, DnaA numbers, cell size, and initiation variables as a function of growth rate based on experimental data of Si *et al.* [11] under various nutrient conditions. In the simulation, the corresponding average experimental values (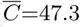 min, 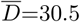 min) were assigned to *C* and *D.* There free parameters were determined by fitting experimental data on cell size as a function of growth rate: *α* = 2.4 × 10^−3^*μm*^3^*/s, K*_*p*_ = 11*μm*^−3^, and *n* = 2.9. Other parameters were predetermined (see the Supporting Text). (A) At steady state with different growth rates, regular oscillations in total DnaA number, origin number, genome content, and cell size are coupled to the cell cycle. (B) The predicted number of origins increases exponentially with the growth rate, exactly following the theoretical formula. (C) By fitting experimental data, cell size basically increases exponentially with growth rate, following the nutrient growth law. Unit cell sizes (initiation masses) from the model (grey open triangles) and experiment (grey open circles) are located around the line 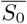 (grey). (D) The predicted total DnaA number per cell increases with growth rate (increase around 50 times from 0.5 dbls/h to 2.5 dbls/h), basically in agreement with experimental data of Chiaramello and Zyskind [83] and Grant *et al.* [35]. The predicted increase is faster than experimental data measured by Hansen *et al.* [84]. (E) The predicted number of each form of DnaA per cell increases with growth rate. (F) *P*_*ini*_ at the initiation of DNA replication is constant since the same initiation threshold was used at different growth rates, whereas other variables at initiation increase ((*x* + *y*)/*V, x*_*f*_ */V, x/Gc*, and (*x* + *y*)*/n*_*box*_) or decrease (*x*_*f*_ */y*_*f*_ and *x/y*) with growth rate. Each initiation variable was divided by the value at a growth rate of 2.85 dbls/h.

As shown in Eq. 5, the transcription rate of DnaA depends on the concentrations of free RNAP and ppGpp. Klumpp and Hwa [52] predicted an increase in free RNAP concentration ([*RNAP*]_*f*_) with growth rate based on an RNAP partition model. If we integrate their mechanistic model directly into our model, our model will become much more complicated. Instead, we coarse-grained their predicted relationship between free RNAP concentration and growth rate by expressing [*RNAP*]_*f*_ as an exponential function of doubling time *τ* (= 1/*μ*), i.e.

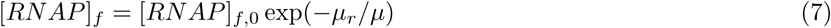

where [*RNAP*]_*f*,0_ and *μ*_*r*_ can be determined by linearly fitting the data of log10[*RNAP*]_*f*_ versus 1/*μ* obtained by Klumpp and Hwa [52]: [*RNAP*]_*f*,0_ = 883*μm*^−3^ and *μ*_*r*_ = 0.68 dbls/h (Fig. S1 A in the Supporting Text). The decrease in ppGpp concentration with growth rate from the experiment [15, 63] can also be roughened as an exponential equation [53], i.e.

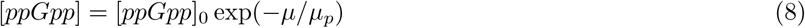

where [*ppGpp*]_0_ = 92 pmol/OD_460_ and *μ*_*p*_ = 1.11 dbls/h, obtained by linearly fitting experimental data of log10[*ppGpp*] versus *μ* [15] (Fig. S1 B). By Equations 5-8, RNAP and ppGpp link DnaA expression and DnaA-mediated DNA replication initiation to the rate of cell mass growth.

In addition to the functional binding sites of DnaA (DnaA-boxes) at *oriC* or DnaA-regulated promoters (e.g. *dnaA* and *nrdAB* promoters), many non-functional DnaA-boxes also spread on the chromosome, especially more than half concentrate at the *datA* locus [64–68]. Titration of these non-functional DnaA-boxes to DnaA molecules is important for controlling DNA replication initiation [24, 25, 66]. As done previously [26], we assume that nonfunctional DnaA boxes that are not at *datA* are homogeneously distributed on the chromosome and duplicate along with the replication. Along with chromosome replication, DnaA-ATP is hydrolyzed to DnaA-ADP, which is commonly referred to as “regulatory inactivation of DnaA (RIDA)” [69–72]. The RIDA depends on the ATPase activity of the Hda-clamp complex which consists of Hda protein and *β* clamp of DNA polymerase III holoenzyme sliding on DNA [69–72]. To express the role of RIDA in the model, we simply believe that DnaA-ATPs bound at one chromosomal site are all inactivated into DnaA-ADPs after replication of this chromosomal site. Furthermore, we assume that DnaA binds to all the nonfunctional DnaA boxes with the same affinity, regardless of the energy form. When DnaA-ATPs bound at one chromosomal site are hydrolyzed, DnaA boxes there are duplicated as well. Thus the hydrolysis rate of DnaA-ATP (*ϵ*(*t*)) can be expressed as *ϵ*(*t*) = (*x*_*b*_(*t*)*/n*_*box*_(*t*))*dn*_*box*_*/dt* (see details in Supporting Text). From the change in the numbers of origins, forks, and DnaA boxes over time, the deactivation rate can be further derived as (see derivation details in the Supporting Text)

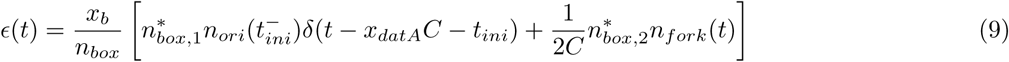

where *n*_*box*_ is the total number of nonfunctional DnaA boxes, 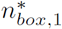 and 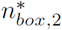 are numbers of DnaA boxes per copy of *datA* and per chromosome excluding the *datA* region, respectively, *x*_*datA*_ = 0.2, indicating the distance of *datA* from *oriC* divided by the chromosomal half-length (*x*_*datA*_ = 0 in the previous model [26]). Here, the replication speed in C period (denoted by *C*) is considered to be a constant. We also checked the effect of different *x*_*datA*_ (see Discussion).

DnaA-ADP can be reactivated to DnaA-ATP in a way dependent on membrane acidic phospholipids or specific genomic sequences DARS [73–76], but the underlying details remain poorly known. For simplicity, we assume that the reactivation rate *σ*(*t*) is linearly dependent on the numbers of free and bound DnaA-ADP (*y*_*f*_ (*t*) and *y*_*b*_(*t*)), *σ*(*t*) = *βy*_*f*_ (*t*) + *β*′*y*_*b*_(*t*), where *β* and *β*′ are reactivation rate constants for free and bound DnaA-ADP, respectively. When *β*′ = *β*, the equation becomes *σ*(*t*) = *βy*(*t*), which was used in the previous model [26]. Theoretically, free DnaA-ADPs should diffuse much faster than DNA-bound DnaA-ADPs to the binding sites on the membrane or/and DARS. Consistent with this conjecture, our fitting of experimental data in [11, 12] for C period prolongation indicates that *β*′ ≪ *β.* For simplicity, therefore, we directly consider

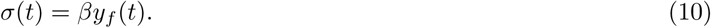

elsewhere in this study. The Fis protein activates the reactivation role of DARS [76], while its concentration increases with the growth rate because of the inhibition of ppGpp [77–79]. So, it is possible that *β* increases with the growth rate via ppGpp regulation. This indicates another mechanistic possibility that ppGpp couples replication initiation to growth rate (see the Discussion and Fig. S25).

Integrating the above processes, the time evolution of total DnaA-ATP (*x*) and total DnaA-ADP (*y*) can be described as [26]

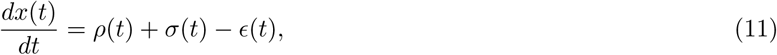

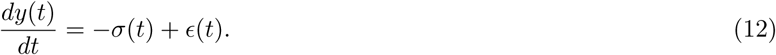

where *ρ*(*t*), *σ*(*t*) and *ϵ*(*t*) indicate DnaA synthesis, reactivation, and inactivation rates, as shown in Eqs. 5, 9, and 10, respectively.

DnaA titration process is much faster than other processes, such as DNA replication, cell growth, DnaA synthesis, reactivation, and inactivation [80]. Therefore, the dynamics of DnaA binding (unbinding) to (from) DnaA boxes can be represented by

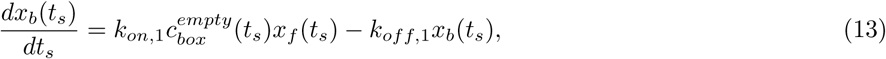

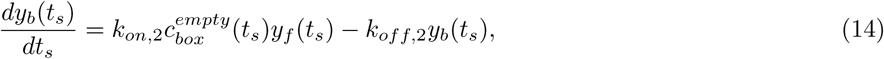

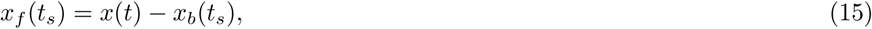

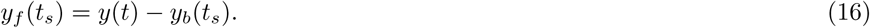

where the concentration of unbound DnaA-boxes *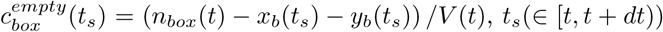* indicates a much smaller time scale, *k*_*on*,1_ and *k*_*on*,2_ indicate binding rates of DnaA-ATP and DnaA-ADP to nonfunctional boxes, and *k*_*off*,1_ and *k*_*off*,2_ indicate the corresponding unbinding rates [26].

As mentioned above, the initiation of *oriC* occurs when the initiation probability reaches the threshold and the eclipse period has passed. For the above two possible mechanisms of ppGpp inhibiting replication initiation, the initiation threshold is considered constant. Recently, Kraemer *et al.* [81] proposed that ppGpp inhibits replication initiation by indirectly affecting supercoiling at *oriC.* We also tested this possible mechanism by adopting a ppGpp(growth rate)-dependent initiation threshold (see the Discussion and Fig. S24).

Following the Helmstetter-Cooper model, we consider the duration of C period (*C*) and the duration of D period (*D*) as two independent parameters, and then replication termination time and cell division time are *t*_*ini*_ + *C* and *t*_*ini*_ + *C* + *D*, respectively (Fig. 1 C). At cell division, all the extensive variables (including cell volume, numbers of DnaA in different forms, the number of origins, the number of DnaA boxes, etc.) in the model are halved.

According to the above model, we simulated the evolution of the levels of DnaA in each form with cell cycles to get a self-consistent cell birth volume. In the simulation, arbitrary initial values were assigned to all variables, but constraints among these variables should be satisfied. We updated the cell birth volume (*V*_0_) for a new cell cycle based on the relative deviation of the simulated inter-division time from the doubling time (*τ*) given in advance by 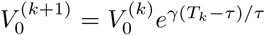, where *T*_*k*_ is the inter-division time of generation *k*, 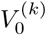 and 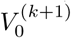 are the birth volumes of generations *k* and *k* + 1, respectively and *γ* = 0.1 (typically). The parameter *γ* in the update formula for the birth volume affects the convergence speed and precision. We can also choose different update formulas, but this will not affect our results. The simulation stops when the relative difference between the inter-division (inter-initiation) time and the preset doubling time is smaller than a threshold or the preset running time (⩾5 ×10^6^s) is over. In the former case, a regular DnaA oscillation coupled with cell cycle will be achieved, so that we can obtain a converged birth volume. Whereas, in the latter case, irregular DnaA oscillation and abnormal cell cycle will be obtained. With proper parameters, a steady-state regular cell cycle can be obtained in the simulation. At steady state, the population average cell size and other desired variables were obtained with the cell age distribution (*g*(*t*) = (2*ln*2*/τ*) ·2^−*t/τ*^) [7, 11]. We used *S* to denote the average cell size (volume) i.e. *S* = ⟨*V*(*t*)⟩, *S*_0_ to denote the average cell size per average origin (unit cell size or initiation mass). With the Cooper-Helmstetter model, one can derive origin number and genome content as a function of *C, D*, and *τ* : *n*_*ori*_ = 2^(*C*+*D*)*/τ*^ [82] and *Gc* = [*τ/*(*C*·*ln*(2))][2^(*C*+*D*)*/τ*^ −2^*D/τ*^] [7]. We used these two theoretical formulas to check if a regular cell cycle is indeed achieved. In the simulation, C, D, and doubling time *τ* are selected by averaging their relevant experimental values. The values of other parameters were derived directly from the literature or by fitting the data in the literature with the least squares method. More details of the simulation and fits can be found in the Supporting Text.

## III. RESULTS

### A. The model shows that as the growth rate increases, the cell size increases approximately exponentially, and the number of DnaA increases significantly

Under various nutrient conditions, Si *et al.* [11] measured cell size, doubling time *τ*, C and D from steady-state populations of MG1655 and NCM3722. Their measurement shows that C and D are constant in a wide range of growth rates, while cell size increases exponentially with growth rate as the nutrient growth law, which can be formulated as

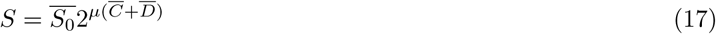

where 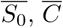, and 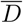 denote experimental averages for *S*_0_, *C*, and *D*, which are independent o<u>f</u> growth rate. From Eq. 17, cell size increases exponentially with growth rate (1/doubling time) with an exponent of 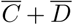, given a constant initiation mass 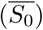. To decipher the nutrient growth law from the level of molecular regulation, we performed a series of simulation runs with different doubling times based on the corresponding experimental data of Si *et al.* [11]. In the simulation, C and D periods were assigned the experimental average values 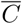 and 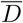, respectively; the parameters *α, K*_*p*_, and *n* are determined by fitting the experimental relationship of cell size and growth rate; other parameters are determined by literature or fitting other data (see the Supporting Text).

When the simulation reaches a steady state, DnaA number, origin number, genome content, and cell volume oscillate regularly with the cell division cycle and all these variables are greater throughout the cell cycle at a higher growth rate (Fig. 2 A). Origin number and genome content predicted by the model completely follow the theoretical formulas 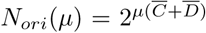 and 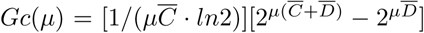 that are consistent with experimental data (Figures 2 B and S2 A) (The theoretical formulas are derived based on the model of Cooper and Helmstetter [7], which has been integrated into our model). This suggests that our simulation did achieve a steady state at each tested growth rate. The coordination between constant C period and shorter inter-division time leads to earlier initiation of DNA replication in the cell cycle (even in the last one) and the formation of multifork (Fig. 2 A; [7, 26]). This also explains the increase in the origin number and genome content (Figs. 2 A-B and S2 A).

By fitting the experimental data of Si *et al.* [11], the obtained cell size increases approximately exponentially with growth rate, which basically obeys the nutrient growth law shown by Eq. 17 (Fig. 2 C). The fluctuation in the simulated relationship may reflect a real phenomenon (since it is surrounded by noisy experimental data) or indicates that our model is incomplete (e.g., missing certain regulator’s effect). In order to maintain a regular cell cycle, the frequency of DnaA-mediated replication initiation should be equal to the frequency of cell division. When the cell grows faster, the frequency of cell division will be higher, and then the frequency of replication initiation will also be higher. As growth rate-dependent factors, ppGpp, RNAP, and cell size may cooperatively adjust the frequency of replication initiation to match the frequency of cell division by affecting the DnaA transcription rate (*ρ*(*t*)) (see Eq. 5). As the growth rate increases, [*RNAP*]_*f*_ goes up, whereas [*ppGpp*] goes down, so *G*([*ppGpp*]) goes up according to Eqs. 6-8. Cell size also increases with the growth rate as shown by Fig. 2 C. According to Eq. 5, the increases in [*RNAP*]_*f*_, *G*([*ppGpp*]), and cell size *V*(*t*) all lead to a higher *ρ*(*t*), which brings a faster oscillation of DnaA level and a higher frequency of DNA replication initiation.

The predicted initiation mass (unit cell size) *S*_0_ is roughly constant over the range of growth rates considered and slightly fluctuates around the experimental mean 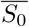 (Fig. 2 C). The difference between the predicted initiation mass and 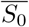 is determined by the difference between cell sizes derived by the fit and the nutrient growth law (17), because the dependence of the simulated number of origins (i.e. *S/S*_0_) on growth rate exactly matches the theoretical formula 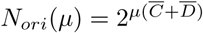 (Fig. 2 B).

Our model also predicts that the total DnaA number per cell increases with the growth rate, which basically agrees with the experimental data of Chiaramello and Zyskind [49] and Grant *et al.* [35] (Fig. 2 D). Faster growth of the cell brings a stronger dilution effect on DnaA and more DnaA boxes which titrate more DnaA molecules. Therefore, the DnaA number is required to increase with the growth rate, in which ppGpp, RNAP, and cell size all make a contribution as discussed above. In addition, we can predict the average numbers and concentrations of DnaA in four specific forms, i.e. DnaA-ATP bound with DNA, DnaA-ADP bound with DNA, free DnaA-ATP and free DnaA-ADP, as a function of growth rate (Figs. 2 E and S2 B). The results show that as the growth rate increases from ∼0.5 to ∼3 doublings/hour, the number of each form of DnaA increases about 100 times (Fig. 2 E). The increase in the DnaA number per cell obtained by Hansen *et al.* [84] is slower than that predicted by our model and measured by Chiaramello and Zyskind [49] and Grant *et al.* [35]; especially, at low growth rates, the number of DnaA from Hansen *et al.* [84] is significantly higher than that from Chiaramello and Zyskind [49] (Fig. 2 D). The fold change in the DnaA protein level from Chiaramello and Zyskind [49] is similar to the fold change in the DnaA mRNA level from their own [83]. The fold change of DnaA protein level from Hansen *et al.* [84] is more consistent with that estimated from mass spectrometry [79]. The two distinct patterns of DnaA expression level depending on growth rate from the experiments may result from the difference in the used strains or growth conditions. Different response mechanisms to the limitation of cell growth may coexist and the dominant mechanism may differ in different strains or growth conditions. We tested another two possible mechanisms of ppGpp inhibiting replication initiation and the obtained dependence of DnaA number on growth rate agrees with the data of Hansen *et al.* [84] (see the Discussion and Figures S24 and S25). Neither of these two mechanisms is achieved through the inhibition of DnaA synthesis, so it is understandable to obtain a slower change in DnaA number with growth rate.

Finally, we compared different initiation conditions by quantifying how the relevant variables at the replication initiation (initiation variables) depend on the growth rate in our model (Fig. 2 F). In our simulation, DNA replication is initiated when the initiation probability (*P*_*ini*_) increases to the threshold. From Eq. 3, *P*_*ini*_ depends on two variables, i.e., the ratio of free DnaA-ATP to free DnaA-ADP (*x*_*f*_ */y*_*f*_) and the concentration of free DnaA-ATP (*x*_*f*_ */V*). Previous models used the other variables to set the threshold initiation conditions (see details in the Introduction). Fig. 2 F presents the dependence of these initiation variables on growth rate. It shows that *P*_*ini*_ at replication initiation is invariant with the growth rate. Under our initiation condition, DNA replication is initiated when *P*_*ini*_ is equal to the threshold, and then *P*_*ini*_ decreases to be below the threshold rapidly. Since we used the same initiation threshold at different growth rates, *P*_*ini*_ at replication initiation should be constant as well (the same for the below cases). Other initiation variables change significantly at low growth rates and slightly at high growth rates (Fig. 2 F). It indicates that our initiation condition is irreplaceable, especially when the cells are growing slowly. Interestingly, as the growth rate increases, *x*_*f*_ */y*_*f*_ and *x/y* at initiation decrease in the same way, while (*x* + *y*)/*V, x*_*f*_ */V, x/Gc*, and (*x* + *y*)*/n*_*box*_ increase in a similar manner. The former two variables reflect the competition between active and inactive DnaA in binding to DnaA boxes, which can be used to set the full competition initiation condition. The latter four roughly reflect DnaA concentration, which can be used to set the non-competition initiation condition. So our initiation condition is a tradeoff between the full competition and non-competition initiation conditions.

### B. The model gives an approximate exponential increase in cell size and a significant increase in DnaA number with the prolongation of C period

Various methods have been used to slow DNA replication (i.e. prolong C period) specifically, for example, titrating ribonucleotide reductase (RNR) gene expression, knocking down DNA helicase Rep, and adding the drug hydroxyurea [11, 12]. In the case of perturbing C period, the growth law can be represented by

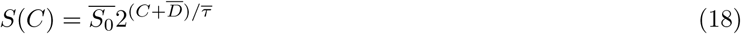

where 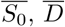 and 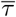 are experimental averages for *S*_0_, *D*, and *τ*, which are independent of *C.* Eq. 18 shows that cell size increases exponentially with *C* with an exponent of 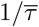, which we call the “replication” growth law.

Based on experimental data for RNR titration [12], Rep knockdown [11], and hydroxyurea addition [11], we simulated the effect of changing the length of C period with our model. The results of using these three data sets are basically similar. This suggests that the effect of C perturbation is essentially independent of the particular perturbed gene or process. Individual different results depend on differences in growth conditions and perturbation intensity. Since the replication duration *C* is a parameter in our model, we varied *C* directly to see if cell size changes exponentially as shown by Eq. 18. In the simulation, the experimental averages (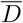 and 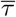) were assigned to the theoretical *D* and *τ.* With a given *τ* (or *μ*), free RNAP concentration, ppGpp concentration, and the intensity of ppGpp inhibition can also be estimated based on Eqs. 6-8. However, these estimates depend on the fitting parameters in Eqs. 6-8, which may result in a significant deviation between the estimated cell size and the empirical cell size at individual growth rates (Fig.2). Therefore, we did not use these estimates but regarded *α*[*RNAP*]_*f*_ *G*([*ppGpp*]) as an independent new parameter *α*′ which can be determined by fitting (the same for the perturbation of D period). Both *α*′ and *β* (reactivation rate constant) were determined by fitting the empirical relationship between cell size and *C* under Rep knockdown [11]. The obtained value of *β* was also used for all other perturbations in this study. In the simulation with the data under hydroxyurea induction [11] and RNR titration [12], *α*′ is the only parameter that fits experimental data.

Simulations have shown that in regular oscillations with the extended C period, DNA replication begins earlier, even at least one round ahead, leading to the coexistence of more origins and more replication forks (Figs. 3 A, S3 A-B, S4 A-C, S5 A-C). At any time during one division cycle of the cell, genome content, DnaA number, and cell volume increase with *C* (Figs. 3 A, S4 A, and S5 A). Oscillating patterns of genomic content and DnaA number also changes in response to earlier initiation (Figs. 3 A, S4 A, and S5 A). The average origin number and the average genome content obtained by the simulation increase with the increase of *C*, exactly following the theoretical formulas 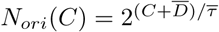 and 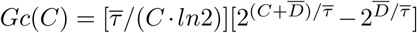 (Fig. S3 A-B, S4 B-C, and S5 B-C). Importantly, the average cell size obtained by fitting experimental data increases with *C* in an approximately exponential way, which roughly captures the replication growth law as shown in Eq. 18 (Fig. 3 B, S4 D, and S5 D). The simulation with the data under RNR titration [12] gives that cell size increases from *C* = 25 min to *C* = 90 min more than 10 times (Fig. 3 B). The increase is a little faster than the experiment given, due to a slight decrease in the experimental growth rate (Fig. 3 B). The simulations with the data under both Rep knockdown and hydroxyurea induction [11] give an around three-fold increase in cell size from *C* = 30 min to *C* = 80 min (Fig. S4 D and S5 D). In both cases, cell size increases more slowly with *C* when *C* > *τ* than when *C < τ.* The transition point *C* = *τ* shifts with the change in *τ* rather than in other parameters. It implies that the change in *C* affects DnaA dynamics less significantly when *C* > *τ* than *C < τ.* Prolongation of the C period alters DnaA dynamics by directly affecting DnaA titration and hydrolysis. The increase in cell size is required for matching the inter-initiation time t<u>o</u> the doubling time in response to the altered DnaA dynamics. The simulated initiation mass stays around the line 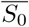 (Fig. 3 B, S4 D, and S5 D), in which the small fluctuation comes from the same reason for that in the simulated cell size.

**FIG 3.**
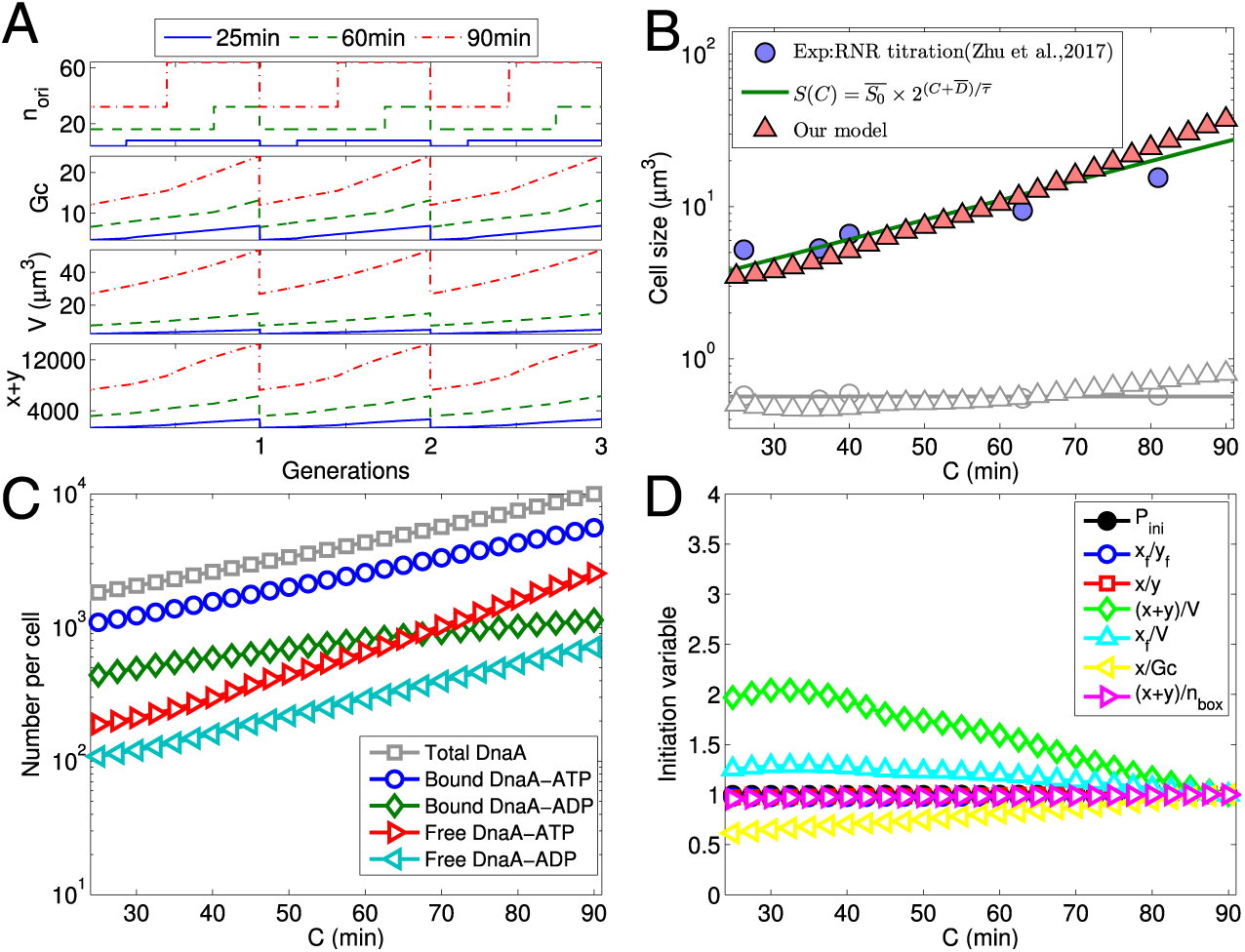
Our model quantifies origin number, genome content, cell size, DnaA numbers, and initiation variables as a function of the duration of C period (*C*) based on experimental data of Zhu *et al.* [12] under ribonucleotide reductase (RNR) titration. The average doubling time and D period (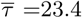 min, 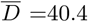 min) were assigned to the theoretical *τ* and *D.* By fitting experimental data on cell size as a function of *C*, we have *α*′ = 0.4*μm*^3^*/s.* Other parameters were determined as shown in the Supporting Text. (A) At steady state with different *C*, regular oscillations of origin number, genome content, DnaA number, and cell size are coupled to the cell cycle. (B) The fitted cell size approximately increases exponentially with *C*, basically following the “replication” growth law. (C) The model predicts that the number of DnaA in each form increases exponentially with *C.* (D) Variables at the initiation of DNA replication increase (*x/Gc*), decrease ((*x* + *y*)/*V, x*_*f*_ */V*) or do not change (*P*_*ini*_, *x*_*f*_ */y*_*f*_, *x/y*, (*x* + *y*)*/n*_*box*_) with *C.* Each initiation variable was divided by the value at *C* = 90 min.

Our model also predicts that the (average) number of each form of DnaA in one cell increases with the duration of C period (Figs. 3 C, S4 E, and S5 E), because of the earlier initiation and more DnaA sites titrating DnaA. The number of each form of DnaA from the simulation increases 3-13 times from *C* = 25 min to *C* = 90 min (with the data under RNR titration [12]) or 2-3 times from *C* = 30 min to *C* = 80 min (with the data under either Rep knockdown or hydroxyurea induction [11]) (Figs. 3 C, S4 E, and S5 E). In addition, the increase in free DnaA-ATP is the greatest, while the increase in bound DnaA-ADP is the least. The concentrations of total DnaA number, bound DnaA-ATP, bound DnaA-ADP, and free DnaA-ADP decrease slightly (bound DnaA-ADP decreases more than others), whereas free DnaA-ATP increases a little (Figs. S3 C, S4 F, and S5 F). This indicates that the change in cell size is slightly larger than the change in the number of each form of DnaA (opposite for free DnaA-ATP). The different changes in bound DnaA-ADP and free DnaA-ATP suggest that prolongation of the C period perturbs DnaA dynamics by mainly affecting DnaA titration and hydrolysis.

Finally, our simulation shows that at the replication initiation, *P*_*ini*_ does not change with *C* as expected (see explanation above), *x/y, x*_*f*_ */y*_*f*_, and (*x* + *y*)*/n*_*box*_ are invariant as well, (*x* + *y*)/*V* and *x*_*f*_ */V* decrease more or less, whereas *x/Gc* increases more or less (Fig. 3 D, S4 G, and S5 G). It suggests that, under the perturbation of C period, our initiation condition can be replaced by that set according to *x/y, x*_*f*_ */y*_*f*_, or (*x* + *y*)*/n*_*box*_, but not replaced by that set by (*x* + *y*)/*V, x*_*f*_ */V*, or *x/Gc.*

### C. The model predicts an exponential increase in cell size and DnaA number with the extension of D period

D period can also be specifically perturbed in a variety of ways, such as by overexpressing SulA, adding the antibiotic cephalexin or titrating the expression of MreB or FtsZ [10, 11]. In the case of only affecting D period, the growth law becomes

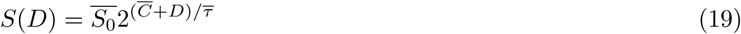

where 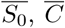, and 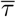 are experimental averages for *S*_0_, *C*, and *τ*, which are independent of *D.* Eq. 19 expresses the cell size an exponential function of D period, with an exponent of 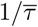, which we call the “division” growth law.

Based on experimental data under MreB/FtsZ titration [10], SulA overexpression [11], and cephalexin addition [11], we used our model to simulate the effect of perturbing D period. Simulations with these data yielded similar results, suggesting that the effect of D perturbation is independent of the particularly perturbed gene or process. In the simulation, we changed the parameter *D* directly. The duration of DNA replication and doubling time were assigned the corresponding experimental averages (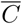 and 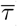), respectively. The only free parameter here is *α*′, which can be fixed by fitting experimental cell size as a function of *D* (see the Supporting Text). Effects of *D* perturbation on oscillations of origin number, genome content, DnaA total number, and cell size are similar to that of *C* perturbation (Figs. 4 A, S7 A, S8 A, and S9 A). The average origin number and average genome content from the simulation increase with *D*, exactly the same as predicted by theoretical formulas 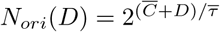 and 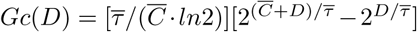 (Figs. S6 A-B, S7 B-C, S8 B-C, and S9 B-C). From fits with experimental data, cell size exponentially increases with D period, perfectly in agreement with Eq. 19; the simulated initiation mass (unit cell size) is located at the experimental value 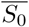, invariant with *D.* Note that the fitted *α*′ here affects the absolute values of cell size and initiation mass, but does not affect the exponential increase in cell size and the invariance of initiation mass. (Figs. 4 B, S7 D, S8 D, and S9 D). As an interesting prediction, the number of each form of DnaA also increases exponentially with *D*, and the exponent is the same as for cell size (Figs. 4 C, S7 E, S8 E, and S9 E). This results in a constant concentration of DnaA (Figs. S6 C, S7 F, S8 F, and S9 F). In addition, the number of each form of DnaA as well as cell size increases by 3-5 times from *D* = 20 min to *D* = 80 min (Figs. 4B-C,S7 D-E, S8 D-E, and S9 D-E). Finally, the initiation variables are invariant with *D* (Figs. 4 D, S7 G, S8 G, and S9 G). In our model, the time of cell division is determined by the time of replication initiation. However, the regulation of DnaA and the replication timing does not depend on the time of cell division. So the change in *D* does not affect DnaA concentration and each initiation variable as a function of the ratios between extensive quantities. Delayed D period leads to the same exponential increase in the number of each form of DnaA, genome content, the number of DnaA boxes, and other extensive quantities as in cell size.

**FIG 4.**
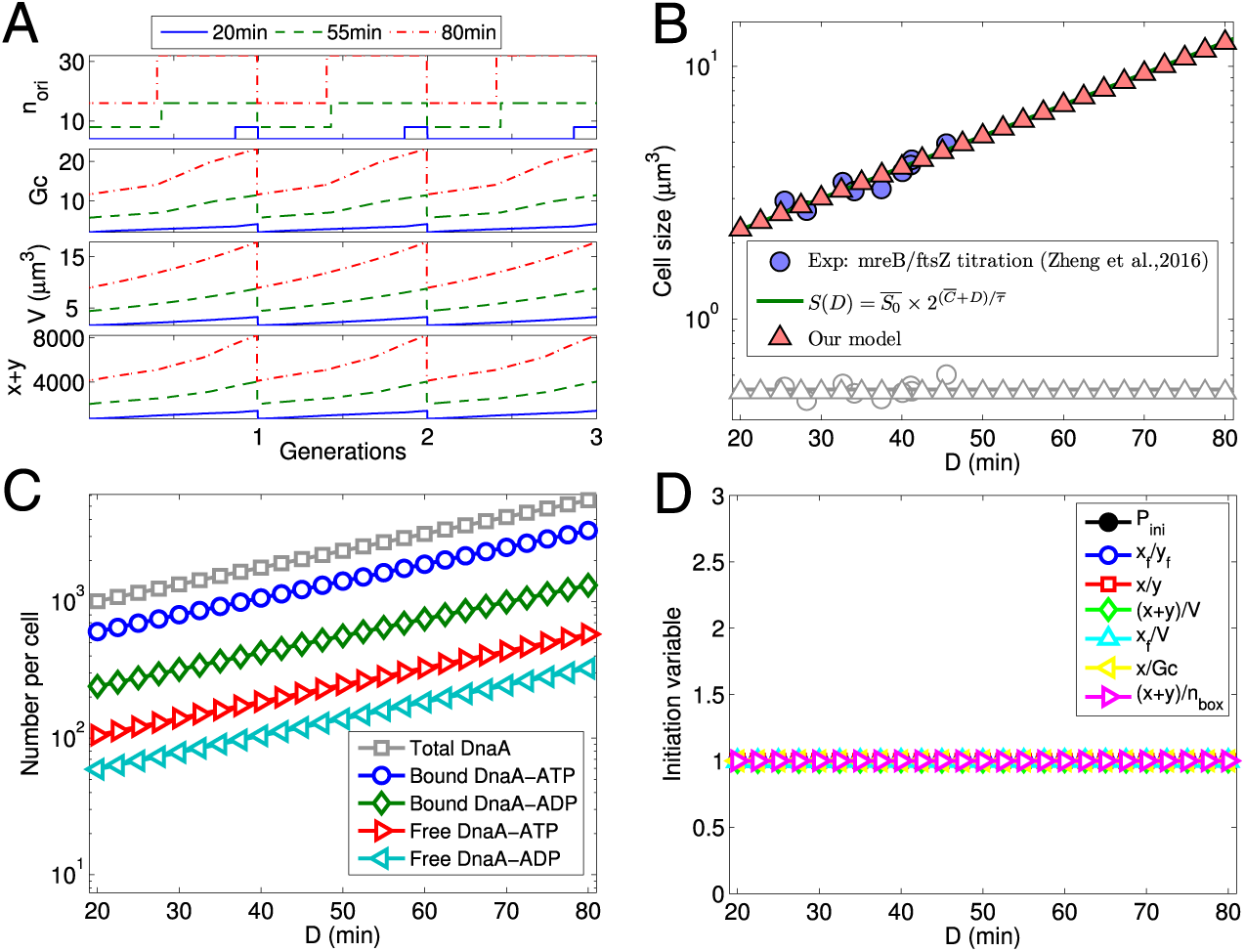
Our model quantifies origin number, genome content, cell size, DnaA numbers and initiation variables as a function of the duration of D period (*D*) based on experimental data of Zheng et al. [10] under mreB/ftsZ titration (growth medium: RDM + glucose). In the simulation, *τ* and *C* were assigned experimental averages (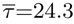 min; 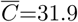 min). By fitting experimental data on cell size as a function of D period, we have *α*′ = 0.28*μm*^3^*/s.* Other parameters are given in the Supporting Text. (A) Regular oscillations of origin number, genome content, total DnaA number, and cell size are coupled with cell cycles at steady state with different *D.* (B) The fitted cell size increases exponentially with *D*, which is completely consistent with the division growth law. (C) As a prediction of the model, the number of DnaA in each form increases exponentially with *D.* (D) Variables at the initiation of DNA replication do not change with *D.* Each initiation variable was divided by the value at *D* = 80 min.

### D. The model predicts a linear increase in cell size and a significant decrease in DnaA concentration with the enlargement of the initiation mass caused by DnaA knockdown or *oriC* block

Under the above three types of perturbations, the initiation mass is constant, so cell size can be predicted as an exponential function of growth rate, *C* or *D.* The initiation mass can be also specifically altered, for example, by *oriC* block or DnaA knowdown with tunable CRISPR interference [11]. Under this type of perturbation, the growth law can be expressed as

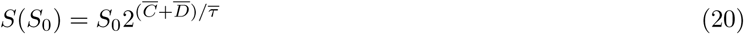

where 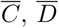, and 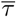 are experimental averages for *C, D*, and *τ*, respectively, which are independent of the initiation mass (*S*_0_). Eq. 20 shows that the cell size increases linearly with *S*_0_, which is called the “initiation mass” growth law.

To simulate the effect of specifically perturbing initiation mass, we considered the effect of RNAP and ppGpp by using Eqs. 7, 8, and 6 in Eq. 5. All parameters were determined from the literature or by fitting other data sets, i.e., no parameter was fitted here. In the simulation, we fixed *C, D*, and *τ* in advance as their experimental averages. Si *et al.* [11] blocked DnaA transcription elongation with tunable CRISPR interference, which corresponds to the decrease in DnaA transcription rate constant (*α*) in our model. Therefore, we simulated the effect of DnaA knowdown by lowering *α.* The simulation shows that both origin number and genome content do not change with *α* (*S*_0_), regardless of their oscillations during in the cell cycle or their averages (Figs. 5 A, S10 A-B, and S11 A-C). A smaller *α* leads to a bigger initiation mass *S*_0_ and the smaller *α*, the larger increase in *S*_0_ with *α* decreasing (Figs. 5 B and S11 D), which is consistent with the result of the DnaA titration experiment [85]. It also explains why DnaA expression from extra copies of the *dnaA* gene has little effect on the timing of replication initiation and the distributions of genome content and cell mass [86]. Note that the quantitative relationship between the initiation mass and *α* shown in Figs. 5 B and S11 D is impossible to predict from a phenomenological model that does not involve the synthesis and regulation of DnaA. Furthermore, the average cell size obtained by the simulation increases linearly with the initiation mass, exactly following the initiation mass growth law shown in Eq.20 (Fig. 5 B and S11 D), irrespective of the choice of parameter values. Note that as *S*_0_ increases, both the experimental genome content and the experimental origin number tend to decrease, and the cell size from the experiment increases more slowly than the cell size obtained by the simulation and the growth law (Figs. 5 B, S10 A-B, and S11 B-D). This is because the increase in *α* actually also brings a small change in growth rate. As Si *et al.* [11] did, we assumed that the growth rate varies linearly with *S*_0_ (i.e., *μ*(*S*_0_) = *a*(*S*_0_) + *b*), and then theoretical formulas for origin number, genome content, and cell size as a function of *S*_0_ can be corrected as: 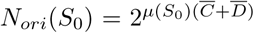 and 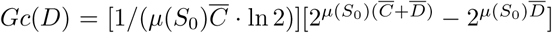, and 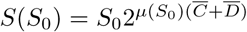. To update the simulation, we did several iterative runs: (1) performing a set of runs to generate *S*_0_ as a function of *α* regardless of the change in growth rate; (2) performing another set of runs by deriving growth rate from the newly generated *S*_0_ with *μ*(*S*_0_) = *aS*_0_ + *b*; (3) redoing step 2 for another 2 times. After the three iterative updates, we captured the decrease in origin number and genome content and the discounted increase in cell size, which were observed in the experiment (Figs. S13 B-D and S14 B-D) As *S*_0_ increases, the number (concentration) of DnaA in each form decreases (Figs. 5 C, S10 C, and S11 E-F), except that the number of free DnaA-ATP either increase (Fig. 5 C) or decrease (Fig. S11 E). This is understandable because the decrease in *α* results in both a decrease in the DnaA synthesis rate and an increase in cell size. The correction with the change in growth rate gives a faster decrease in the number (concentration) of DnaA in each form (or turning from increase to decrease). Figs. 5 D and S11 G show that, as *S*_0_ increases, *P*_*ini*_ at initiation is constant as expected, *x*_*f*_ */y*_*f*_ and *x/y* at initiation increase with similar slopes, while (*x* + *y*)/*V, x*_*f*_ */V, x/Gc*, and (*x* + *y*)*/n*_*box*_ at initiation all decrease. The correction that takes into account the decrease in growth rate just increases the difference between the three sets of initiation variables (Figs. S13 G and S14 G). These results indicate that when the initiation mass is changed, our initiation condition cannot be replaced by others and again our initiation condition is a tradeoff between full competition and non-competition initiation conditions.

In order to simulate the effect of *oriC* block [11], we varied the initiation threshold of *oriC* (Fig. S12 and S15). Oscillations of origin number, genome content, and DnaA number and the change in origin number, genome content, or cell size with *S*_0_ are similar to the above results for DnaA knockdown (Fig. S12 A-D and S15 A-D). The initiation mass *S*_0_ increases with the initiation threshold and at a bigger initiation threshold, the increase in *S*_0_ is larger (Fig. S12 D and S15 B). Such quantitative dependence of the initiation mass on the initiation threshold cannot be obtained from a phenomenological model without an initiator. With *S*_0_ increasing, numbers of total DnaA and bound DnaA-ATP are basically constant, the number of bound DnaA-ADP decreases, whereas numbers of free DnaA-ATP and free DnaA-ADP increase (Figs. S12 E and S15 E). The concentration of each form of DnaA more or less decreases with *S*_0_ (Figs. S12 F and S15 F). At the replication initiation, *P*_*ini*_ increases with *S*_0_ because of the increase in the initiation threshold, (*x* + *y*)/*V* and *x*_*f*_ */V* decrease, whereas *x*_*f*_ */y*_*f*_, *x/y, x/Gc*, (*x* + *y*)*/n*_*box*_ increase (Figs. S12 G and S15 G). Therefore, our model not only captures the initiation mass growth law but also predicts the similarity and difference of the changes in the average level of each form of DnaA between different perturbation methods.

**FIG 5.**
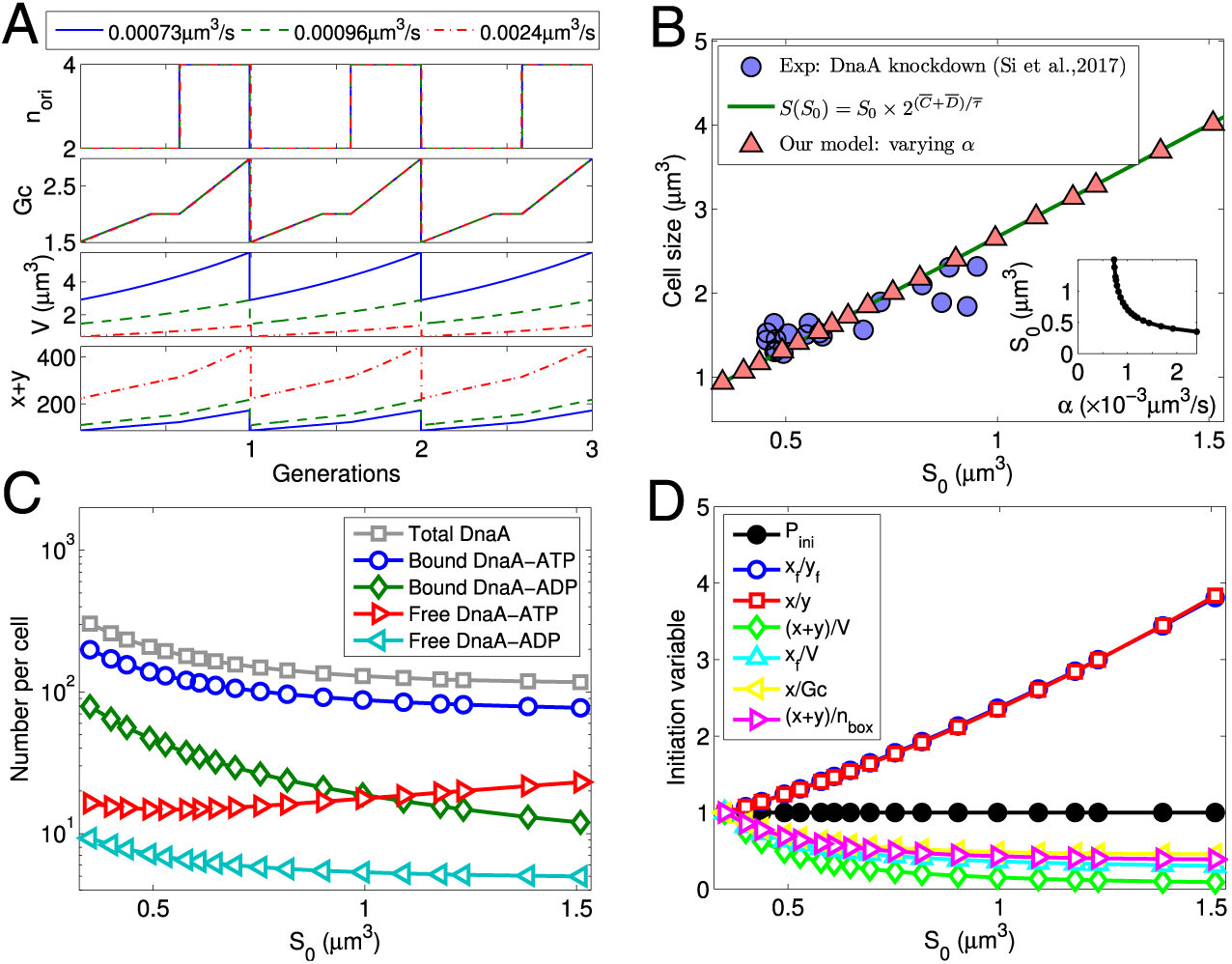
The model quantifies the changes in origin number, genome content, cell size, DnaA levels, and initiation variables with increasing initiation mass (varying *α*) based on experimental data of Si *et al.* [11] under DnaA knowdown (growth medium: MOPS glucose + 6 a.a.) In the simulation, doubling time and durations of C and D periods were assigned experimental averages (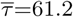 min, 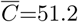min, 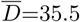 min). No fitting was done here. Other parameters were predetermined (see the Supporting Text). (A) At steady state with different values of *α*, regular oscillations of origin number, genome content, DnaA level and cell size are coupled with the cell cycle. (B) As a prediction of the model, cell size increases linearly with *S*0, precisely following the initiation mass growth law. The inset plot shows that initiation mass *S*_0_ increases with *α.* (C) As another prediction of the model, the number of each form of DnaA decreases with *S*_0_ except that the number of free DnaA-ATP increases. (D) The initiation probability (*P*_*ini*_) at replication initiation does not change with *S*_0_ since the same initiation threshold was used. Other initiation variables at initiation increase (*x/y* and *x*_*f*_ */y*_*f*_) or decrease ((*x* + *y*)/*V, x*_*f*_ */V, x/Gc* and (*x* + *y*)*/n*_*box*_) with *S*_0_. Each initiation variable was divided by the value at *α* = 2.4 *×* 10^−3^*μm*^3^*/s* (*S*_0_ = 0.39*μm*^3^).

## IV. DISCUSSION

Our model can predict changes in cell size, initiation mass, and DnaA number under more perturbations, such as altering *k*_*off*,1_*/k*_*on*,1_, *k*_*off*2_*/k*_*on*,2_, *β, k*_1_, *k*_2_, 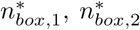, *x*_*datA*_, *K*_1_, and *K*_2_ (Figs. S16-S21). Cell size (initiation mass) is positively correlated with the frequency of replication initiation. Therefore, an increase in cell size (initiation mass) for maintaining a constant initiation frequency means a decrease in the initiation frequency for maintaining a fixed cell size (initiation mass). The simulation shows that cell size and initiation mass increase with 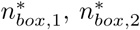, *k*_*off*,1_*/k*_*on*,1_, *k*_1_, *k*_2_, *x*_*datA*_, *K*_1_, and initiation threshold, and decrease with *α, k*_*off*,2_*/k*_*on*,2_, *β* and *K*_2_ (Figs. S16, S18, and S20). This is consistent with the dependence of initiation frequency on these parameters when cell size and initiation mass are fixed in advance [26]: the initiation frequency decreases with the former parameters and increases with the latter ones. The simulation also gives that DnaA number per cell increases with *α, k*_*off*,1_*/k*_*on*,1_, 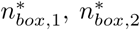, *K*_1_, and initiation threshold, decreases with *k*_1_, *k*_2_, *β, x*_*datA*_, and *K*_2_, and changes with *k*_*off*,2_*/k*_*on*,2_ little (Figs. S17, S19, S21). In a wet lab, the above parameters could be changed by mutation or other techniques (such as CRISPR interference [11]) to see if the results meet our expectations. Notice that the perturbation of some parameters may affect the growth rate as well, which results in the emergence of a mixed effect. In this case, the corresponding correction should be performed as we did above for DnaA knockdown and *oriC* block. From Figs. S16-S21, we also found that some parameters (such as *α, k*_1_, *k*_*off*,2_*/k*_*on*,2_, *x*_*datA*_, and *K*_2_) significantly affect the pattern of the change in cell size (initiation mass) with the growth rate. If the value of one of the above parameters in a particular bacterial strain or under a special growth condition differs from that in the wide type strain or under the normal growth condition, the initiation mass may be variant with the growth rate, and cell size may not increase with growth rate as predicted by the nutrient growth law (e.g., [20, 21, 85]).

The simulation also shows that if a parameter is outside its normal range, a regular oscillation cannot be obtained. To clarify which processes in the model are essential for a good coupling between replication initiation and the cell cycle, we eliminated the effect of each process separately by assigning zero or an extremely large value to the corresponding parameter (variable) at the (normal) growth rates between 0.5 and 3 doublings/hour. We found that *α* = 0, *k*_*on*,1_ = 0, *K*_1_ = 100, *K*_2_ = 100, initiation threshold = 0.9, and *ϵ*(*t*) ≡ 0, always infeasible to form a regular oscillation; *β* = 0 and *k*_*on*,2_ = 0, infeasible at slow growth; 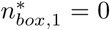, infeasible at fast growth; *k*_1_ = 0, *k*_2_ = 0, and 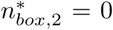, always feasible. It indicates that the crucial processes for a regular cell cycle include DnaA expression, titration to DnaA-ATP, titration to DnaA-ADP (at slow growth), titration at *datA* (at fast growth), hydrolysis of DnaA-ATP, reactivation of DnaA-ADP (at slow growth), binding of DnaA-ATP and DnaA-ADP to *oriC* and initiation mediated by DnaA-ATP. The remaining processes do not seem to matter. These results are basically consistent with the increased asynchrony index observed for mutations of DnaA, *datA*, Hda, and DARS [75, 87–90]. It requires further quantitative experiments to check the dependence of the importance of all the processes on growth rate and even other perturbation parameters.

The distribution of DnaA boxes, especially the location of *datA*, on the chromosome can also affect the initiation of DNA replication [24, 88, 89]. By simulating with *datA* at different positions, we found that the DnaA-ATP/DnaA-ADP ratio increases with the distance between *datA* and *oriC* (Fig. S22 A), in agreement with the speculation of Frimodt-Møller *et al.* [89]. Moreover, it is difficult to obtain a regular oscillation at a high growth rate when *datA* is far from *oriC* (Fig. S22 B). In the plot of the relative distance from *datA* to *oriC* versus growth rate, a phase diagram is formed with two distinct regimes: a feasible one where a regular oscillation emerges in the simulation and an infeasible one where no regular oscillation emerges (Fig. S22 B). It indicates that the position of *datA* is involved in building a regular DnaA oscillation. The closeness of *datA* to *oriC* should be required for timely hydrolysis and titration of DnaA at *datA* and a fast decrease in the probably of re-initiation after initiation. This is in line with the experimental result of Kitagawa *et al.* [88]: the movement of the position of *datA* does not affect the frequency and synchrony of initiation when *E.coli* cells grew in a minimal medium, whereas the abnormal initiation occurs when *datA* was moved to the locus closer to the terminus and the growth medium is rich. At faster cell growth, the dosage of *datA* is higher, then the titration role is stronger, and finally, the effect on DnaA dynamics is higher when *datA* is moved closer to the terminus. It was also reported that *datA* can hydrolyze DnaA-ATP independent of Hda-*β*-clamp-mediated RIDA, but *datA*-mediated hydrolysis is weaker than Hda-*β*-clamp-mediated RIDA [91, 92]. Considering this fact in the model may make the predictions of *datA* perturbations more comparable to experimental data. It has also been found that the chromosomal positions of DARS1 and DARS2 associated with DnaA reactivation are nontrivial for cell cycle control and bacterial fitness [89, 90]. If one knows more details about the role of DARS in DnaA reactivation, our model could even be improved to check the effect of the DARS location.

When the eclipse time is changed, the results change little, which implies that the eclipse time affects the robustness of replication initiation, rather than the average initiation frequency [24, 26]. The eclipse time is regulated by SeqA protein [43–45]. The previous experiment has shown that the knockdown of SeqA leads to an increase in cell size, C period, and D period [11], in which case the increase in cell size probably results from the increase in C and D periods, according to the general growth law. Therefore, SeqA may be not only in charge of the robustness of replication initiation but also involved in maintaining the initiation frequency by regulating replication elongation and cell division.

In our model, we considered several growth rate-dependent factors that affect DnaA dynamics and the coordination of replication initiation and cell growth at different growth rates. They include cell size, the dosage of *dnaA* gene and *datA*, genome content, RNAP, and ppGpp. Cell size is directly controlled by cell division mechanisms [93, 94]. In the model, cell size is a self-consistent variable from the coupling of DnaA dynamics to cell cycle. The gene dosage and genome content are strictly determined by the preset C, D, and growth rate in theory. The concentration of free RNAP is hardly measured directly in the experiment. Fortunately, it was predicted from a RNAP partition model and its dependence on growth rate was supported by the expression data from constitutive promoters [51–53]. If we remove the growth rate-dependence of free RNAP concentration in the model, with growth rate increasing, the increase in cell size will be faster while the increase in DnaA number will be slower over small growth rates (*<*∼1 dbls/h) (Fig. S23). This is in line with the faster increase in free RNAP concentration at slower cell growth, while closer to saturation at faster cell growth [52]. ppGpp is a major global regulator of controlling growth rate [56–58]. ppGpp is probably involved in the growth rate-dependent matching between replication initiation and cell division frequencies by directly inhibiting DnaA synthesis [49, 50, 59–61]. The inhibition of DnaA synthesis may be achieved through ppGpp inhibiting the transcription initiation from the *dnaA* promoter which contains a GC-rich discriminator region or through ppGpp affecting Fis level, DNA supercoiling, or the divergent ribosomal promoter *rpmH* [35, 50]. It may also be through ppGpp limiting translation capacity at slow growth [58]. If we remove the dependence of ppGpp on growth rate in the model, cell size will increase much more while the DnaA number will increase much less over the whole normal range of growth rates (Fig. S23). It indicates an potentially important role of ppGpp in the coordination between DNA replication and growth rate.

ppGpp may also inhibit the replication initiation not through directly affecting DnaA synthesis [76, 81, 95, 96]. A recent study showed that ppGpp can indirectly regulate the supercoiling at *oriC* to inhibit the initiation of DNA replication [81]. It implies another possible mechanism of coupling DNA replication to growth rate: ppGpp inhibits replication initiation by raising the initiation threshold. In addition, Fis protein has been found to activate the reactivation role of DARS [76]. Fis concentration increases with growth rate because of the inhibition of ppGpp [77– 79]. It provides a third possible mechanism of coupling DNA replication to growth rate: ppGpp inhibits replication initiation by lowering the reactivation rate. In order to test the effects of these two mechanisms separately, we removed the inhibition of ppGpp on DnaA expression (i.e. replacing *G*([*ppGpp*]) with a constant). Then we assumed that the initiation threshold increases with ppGpp concentration for the second mechanism, and the reactivation rate factor *β* decreases with ppGpp concentration for the third mechanism, respectively. By redoing the simulations, we obtained similar changes in origin number, genome content, and cell size, while a smaller increase in DnaA number and a more constant DnaA concentration (Figs. S24 and S25), compared to those under the mechanism of ppGpp inhibiting DnaA expression. Surprisingly, the relationship between DnaA number and growth rate under the second and third mechanisms is consistent with the experimental data of Hansen *et al.* [84] (Figs. S24 F and S25 F). Moreover, the ratio of DnaA-ATP to DnaA-ADP increases slowly with growth rate in the third mechanism, but decreases rapidly in the other two mechanisms (Figs. S24 I and S25 I). Thus, here we provide a feasible method (i.e., measuring the dependence of the DnaA number (concentration) and DnaA-ATP/DnaA-ADP on growth rate) to determine which mechanism is dominant in the experiment. In different strains or under different growth conditions, the dominant mechanism may be different, corresponding to the difference in the growth rate-dependence of the DnaA level and DnaA-ATP/DnaA-ADP ratio.

Elevated ppGpp does not affect the completion of replication elongation [59–61, 97], but it may reduce the elongation rate by inhibiting primase activity in *E.coli* [98] just like in *B. subtilis* [96]. Consistent with this finding, the measured C period tends to be longer at a smaller growth rate [11, 15]. By integrating the decrease in C period with growth rate (fitting with the experimental data of Si *et al.* [11]) into the simulation for the growth rate variation, we did not find any significant changes in the results.

For the prolonged C period, we considered a constantly limited replication elongation speed. DNA replication fork can be stalled for a number of reasons, such as DNA break, lack of nucleotides, presence of alkylation on the DNA template, and barriers in the normal process [99]. These perturbations can also increase the C period duration, but they affect the elongation rate unevenly. A simple test by adding a stall at two symmetric replication sites shows a reduction in cell size, suggesting that the growth law fails in this case. It could be validated by inducing replication fork to stall in the experiment.

Multiple molecular mechanisms (for example, Min system oscillation, nucleotide occlusion, and FtsZ assembly) affect cell division timing and cell size [93, 94]. In our model, the time of cell division is phenologically determined by the time of replication initiation, as in the Cooper-Helmstetter and Donachie models. One underlying idea is that the cell can divide only after DNA replication is finished because a molecular signal (SlmA protein in *E. coli*) binding to the chromosome interferes with the formation of FtsZ ring, the so-called “nucleotide occlusion” [93, 94].

Current theoretical models of cell division mainly focused on the mechanism of Min protein oscillation, FtsZ assembly, or division protein allocation (e.g. [28, 100, 101]). A model that integrates Min system oscillation, nucleotide occlusion, and FtsZ assembly may be helpful for a more comprehensive understanding of the coordination between DNA replication and cell division as well as the general growth law. The noise in DnaA expression, DNA replication, cell division, cell elongation rate, etc. can cause the variability in the single-cell size [102], although the homeostasis remains for cell size [103]. An integrative and stochastic model based on this study might be able to explain the change in the distribution of cell size under different perturbations and provide another view of cell size homeostasis.

## V. CONCLUSION

Here we developed a kinetic model of DnaA-mediated replication initiation to reproduce the general bacterial growth law, which phenomenologically relates cell size to cell cycle timings (doubling time, C and D periods) in bacteria. Compared with the previous initiation models [22, 24–26, 34, 35], our model involves more up-to-date DnaA regulation details that affect replication initiation. However, we did not consider any molecular mechanism that regulates C and D periods. Instead, we assume that cell division occurs after a time of *C*+*D* from replication initiation, as done in the Cooper-Helmstetter model [7]. This allows us to focus on how the general growth law emerges from the coordination of DnaA dynamics and the entire cell cycle. In our model, the parameters should be self-consistent to form a regular oscillation of DnaA and make the inter-initiation time equal to the inter-division time. It explains why the converged cell size is obtained in the iterative update. When a parameter deviates too far from the normal value, the regular oscillation cannot be obtained regardless of how cell size changes. It reflects physical constraints on biological parameters. Our model basically reproduces the general growth law, when doubling time, C period, D period or initiation mass is perturbed specifically. It suggests the invariance of initiation mass first proposed by Donachie [9] can be generally achieved by fine-tuning DnaA oscillation. The simulation of DnaA dynamics also predicts the change in the level of each form of DnaA with the specifically perturbed parameter. These results can be tested with more experiments. Especially, the predictions with the three different regulatory mechanisms of ppGpp inhibiting replication initiation could be used to determine the major mechanism in response to growth rate changing in a specific strain or under a specific cultivation condition. Because of its predictability, our model may play a guiding role in synthetic biology than a more coarse-grained initiation model.

## ACKNOWLEDGMENTS

This work was supported by the Strategic Priority Program of Chinese Academy of Sciences (No. XDA17010504), the National Natural Science Foundation of China (Grant Nos. 11774359 and 11947302), and the Interdisciplinary Innovation Team of Chinese Academy of Sciences (No. 2060299). We thank Bianca Sclavi and Marco Cosentino Lagomarsino for helpful discussion.

## References

[1] D. P. Haeusser and P. A. Levin, Current opinion in microbiology 11, 94 (2008).

[2] S. Cooper, Journal of bacteriology 170, 5001 (1988).

[3] S. Cooper, Theoretical Biology and Medical Modelling 3, 10 (2006).

[4] P. Wang, L. Robert, J. Pelletier, W. L. Dang, F. Taddei, A. Wright, and S. Jun, Current biology 20, 1099 (2010).

[5] M. Mir, Z. Wang, Z. Shen, M. Bednarz, R. Bashir, I. Golding, S. G. Prasanth, and G. Popescu, Proceedings of the National Academy of Sciences 108, 13124 (2011).

[6] M. Osella, E. Nugent, and M. C. Lagomarsino, Proceedings of the National Academy of Sciences 111, 3431 (2014).

[7] S. Cooper and C. E. Helmstetter, Journal of molecular biology 31, 519 (1968).

[8] M. Schaechter, O. Maaløe, and N. O. Kjeldgaard, Microbiology 19, 592 (1958).

[9] W. D. Donachie, Nature 219, 1077 (1968).

[10] H. Zheng, P. Y. Ho, M. Jiang, B. Tang, W. Liu, D. Li, X. Yu, N. E. Kleckner, A. Amir, and C. Liu, Proc Natl Acad Sci U S A 113, 15000 (2016).

[11] F. Si, D. Li, S. E. Cox, J. T. Sauls, O. Azizi, C. Sou, A. B. Schwartz, M. J. Erickstad, Y. Jun, X. Li, et al., Current Biology 27, 1278 (2017).

[12] M. Zhu, X. Dai, W. Guo, Z. Ge, M. Yang, H. Wang, and Y. P. Wang, Mbio 8 (2017).

[13] C. E. Helmstetter and S. Cooper, Journal of molecular biology 31, 507 (1968).

[14] W. Donachie, K. Begg, and M. Vicente, Nature 264, 328 (1976).

[15] H. Bremer and P. P. Dennis, “Escherichia coli and salmonella: Cellular and molecular biology, 2nd ed.” (American Society for Microbiology, Washington, DC, USA, 1996) Chap. Modulation of chemical composition and other parameters of the cell by growth rate, pp. 1553–1569.

[16] M. E. Sharpe, P. M. Hauser, R. G. Sharpe, and J. Errington, Journal of bacteriology 180, 547 (1998).

[17] M. Basan, M. Zhu, X. Dai, M. Warren, D. Sévin, Y.-P. Wang, and T. Hwa, Molecular systems biology 11, 836 (2015).

[18] M. Wallden, D. Fange, E. G. Lundius, Ö. Baltekin, and J. Elf, Cell 166, 729 (2016).

[19] X. Dai, Z. Shen, Y. Wang, and M. Zhu, mSphere 3, e00567 (2018).

[20] G. Churchward, E. Estiva, and H. Bremer, Journal of bacteriology 145, 1232 (1981).

[21] S. Wold, K. Skarstad, H. B. Steen, T. Stokke, and E. Boye, The EMBO journal 13, 2097 (1994).

[22] J. M. Mahaffy and J. W. Zyskind, Journal of theoretical biology 140, 453 (1989).

[23] J. Keasling, H. Kuo, and G. Vahanian, Journal of theoretical biology 176, 411 (1995).

[24] F. G. Hansen, B. B. Christensen, and T. Atlung, Research in Microbiology 142, 161 (1991).

[25] S. T. Browning, M. Castellanos, and M. L. Shuler, Biotechnology and bioengineering 88, 575 (2004).

[26] Q. Zhang and H. Shi, Journal of theoretical biology 314, 164 (2012).

[27] A. Amir, Physical Review Letters 112, 208102 (2014).

[28] F. Bertaux, J. Von Kügelgen, S. Marguerat, and V. Shahrezaei, bioRxiv, doi: 10.1101/078998 (preprint posted February 22, 2019) (2019).

[29] P. Thomas, G. Terradot, V. Danos, and A. Y. Weiße, Nature communications 9, 4528 (2018).

[30] C. Speck and W. Messer, The EMBO journal 20, 1469 (2001).

[31] M. L. Mott and J. M. Berger, Nature Reviews Microbiology 5, 343 (2007).

[32] J. M. Kaguni, Annu. Rev. Microbiol. 60, 351 (2006).

[33] K. Skarstad and T. Katayama, Cold Spring Harbor perspectives in biology 5, a012922 (2013).

[34] W. D. Donachie and G. W. Blakely, Current Opinion in Microbiology 6, 146 (2003).

[35] M. A. Grant, C. Saggioro, U. Ferrari, B. Bassetti, B. Sclavi, and M. C. Lagomarsino, BMC systems biology 5, 201 (2011).

[36] F. Jacob, S. Brenner, and F. Cuzin, in Cold Spring Harbor symposia on quantitative biology, Vol. 28 (Cold Spring Harbor Laboratory Press, 1963) pp. 329–348.

[37] L. Sompayrac and O. Maaløe, Nature New Biology 241, 133 (1973).

[38] W. Messer, Fems Microbiology Reviews 26, 355 (2002).

[39] A. C. Leonard and J. E. Grimwade, Molecular Microbiology 55, 978 (2005).

[40] D. T. Miller, J. E. Grimwade, T. Betteridge, T. Rozgaja, J. C. Torgue, and A. C. Leonard, Proceedings of the National Academy of Sciences of the United States of America 106, 18479 (2009).

[41] A. C. Leonard and J. E. Grimwade, Annual Review of Microbiology 65, 19 (2011).

[42] C. Helmstetter, “Escherichia coli and salmonella typhimurium: Cellular and molecular biology.” (Washington, DC: American Society for Microbiology Press, 1996) Chap. Timing of synthetic activities in the cell cycle, p. 1627C1649.

[43] N. K. Torheim and K. Skarstad, The EMBO journal 18, 4882 (1999).

[44] U. von Freiesleben, M. A. Krekling, F. G. Hansen, and A. Løbner-Olesen, The EMBO journal 19, 6240 (2000).

[45] A. Løbner-Olesen, O. Skovgaard, and M. G. Marinus, Current opinion in microbiology 8, 154 (2005).

[46] J. L. Campbell and N. Kleckner, Cell 62, 967 (1990).

[47] J. Olsson, S. Dasgupta, O. G. Berg, and K. Nordström, Molecular microbiology 44, 1429 (2002).

[48] C. Speck, C. Weigel, and W. Messer, The EMBO journal 18, 6169 (1999).

[49] A. E. Chiaramello and J. W. Zyskind, Journal of Bacteriology 172, 2013 (1990).

[50] C. Saggioro, A. Olliver, and B. Sclavi, Biochemical Journal 449, 333 (2013).

[51] S.-T. Liang, M. Bipatnath, Y.-C. Xu, S.-L. Chen, P. Dennis, M. Ehrenberg, and H. Bremer, Journal of molecular biology 292, 19 (1999).

[52] S. Klumpp and T. Hwa, Proceedings of the National Academy of Sciences 105, 20245 (2008).

[53] Q. Zhang, E. Brambilla, R. Li, H. Shi, M. C. Lagomarsino, and B. Sclavi, bioRxiv, doi: 10.1101/599183 (preprint posted October 09, 2019) (2019).

[54] F. G. Hansen, E. B. Hansen, and T. Atlung, The EMBO journal 1, 1043 (1982).

[55] S. Klumpp, Z. Zhang, and T. Hwa, Cell 139, 1366 (2009).

[56] A. Srivatsan and J. D. Wang, Current Opinion in Microbiology 11, 100 (2008).

[57] K. Potrykus, H. Murphy, N. Philippe, and M. Cashel, Environmental microbiology 13, 563 (2011).

[58] X. Dai, M. Zhu, M. Warren, R. Balakrishnan, V. Patsalo, H. Okano, J. R. Williamson, K. Fredrick, Y.-P. Wang, and T. Hwa, Nature microbiology 2, 1 (2016).

[59] A. Levine, F. Vannier, M. Dehbi, G. Henckes, and S. J. Séror, Journal of molecular biology 219, 605 (1991).

[60] J. W. Zyskind and D. W. Smith, Cell 69, 5 (1992).

[61] G. Schreiber, E. Z. Ron, and G. Glaser, Current microbiology 30, 27 (1995).

[62] K. Sekimizu, D. Bramhill, and A. Kornberg, Cell 50, 259 (1987).

[63] J. Ryals, R. Little, and H. Bremer, Journal of bacteriology 151, 1261 (1982).

[64] R. Kitagawa, H. Mitsuki, T. Okazaki, and T. Ogawa, Molecular microbiology 19, 1137 (1996).

[65] A. Roth and W. Messer, Molecular microbiology 28, 395 (1998).

[66] T. Ogawa, Y. Yamada, T. Kuroda, T. Kishi, and S. Moriya, Molecular microbiology 44, 1367 (2002).

[67] J. Herrick and B. Sclavi, Molecular microbiology 63, 22 (2007).

[68] S. Nozaki, H. Niki, and T. Ogawa, Journal of bacteriology 191, 4807 (2009).

[69] T. Katayama, T. Kubota, K. Kurokawa, E. Crooke, and K. Sekimizu, Cell 94, 61 (1998).

[70] J.-i. Kato and T. Katayama, The EMBO journal 20, 4253 (2001).

[71] M. Su’etsugu, T.-r. Shimuta, T. Ishida, H. Kawakami, and T. Katayama, Journal of Biological Chemistry 280, 6528 (2005).

[72] J. S. Kim, M. T. Nanfara, S. Chodavarapu, K. S. Jin, V. M. Babu, M. A. Ghazy, S. Chung, J. M. Kaguni, M. D. Sutton, and Y. Cho, Nucleic acids research 45, 3888 (2017).

[73] K. Boeneman and E. Crooke, Current opinion in microbiology 8, 143 (2005).

[74] A. Aranovich, G. Y. Gdalevsky, R. Cohen-Luria, I. Fishov, and A. H. Parola, Journal of Biological Chemistry 281, 12526 (2006).

[75] K. Fujimitsu, T. Senriuchi, and T. Katayama, Genes & development 23, 1221 (2009).

[76] K. Kasho, K. Fujimitsu, T. Matoba, T. Oshima, and T. Katayama, Nucleic acids research 42, 13134 (2014).

[77] O. Ninnemann, C. Koch, and R. Kahmann, The EMBO journal 11, 1075 (1992).

[78] P. Mallik, B. J. Paul, S. T. Rutherford, R. L. Gourse, and R. Osuna, Journal of bacteriology 188, 5775 (2006).

[79] A. Schmidt, K. Kochanowski, S. Vedelaar, E. Ahrné, B. Volkmer, L. Callipo, K. Knoops, M. Bauer, R. Aebersold, and M. Heinemann, Nature biotechnology 34, 104 (2016).

[80] S. Schaper and W. Messer, Journal of Biological Chemistry 270, 17622 (1995).

[81] J. A. Kraemer, A. G. Sanderlin, and M. T. Laub, mBio 10, e01330 (2019).

[82] H. Bremer and G. Churchward, Journal of Theoretical Biology 69, 645 (1977).

[83] A. E. Chiaramello and J. W. Zyskind, Journal of bacteriology 171, 4272 (1989).

[84] F. G. Hansen, T. Atlung, R. Braun, A. Wright, P. Hughes, and M. Kohiyama, Journal of Bacteriology 173, 5194 (1991).

[85] A. Løbner-Olesen, K. Skarstad, F. G. Hansen, K. von Meyenburg, and E. Boye, Cell 57, 881 (1989).

[86] I. Flåtten, S. Fossum-Raunehaug, R. Taipale, S. Martinsen, and K. Skarstad, PLoS genetics 11, e1005276 (2015).

[87] E. Boye, T. Stokke, N. Kleckner, and K. Skarstad, Proceedings of the National Academy of Sciences 93, 12206 (1996).

[88] R. Kitagawa, T. Ozaki, S. Moriya, and T. Ogawa, Genes & development 12, 3032 (1998).

[89] J. Frimodt-Møller, G. Charbon, K. A. Krogfelt, and A. Løbner-Olesen, PLoS genetics 12, e1006286 (2016).

[90] Y. Inoue, H. Tanaka, K. Kasho, K. Fujimitsu, T. Oshima, and T. Katayama, Genes to Cells 21, 1015 (2016).

[91] K. Kasho and T. Katayama, Proceedings of the National Academy of Sciences 110, 936 (2013).

[92] K. Kasho, H. Tanaka, R. Sakai, and T. Katayama, Journal of Biological Chemistry 292, 1251 (2017).

[93] L. Rothfield, A. Taghbalout, and Y.-L. Shih, Nature Reviews Microbiology 3, 959 (2005).

[94] J. Lutkenhaus, Annu. Rev. Biochem. 76, 539 (2007).

[95] T. Ogawa and T. Okazaki, Molecular and General Genetics MGG 230, 193 (1991).

[96] J. D. Wang, G. M. Sanders, and A. D. Grossman, Cell 128, 865 (2007).

[97] D. J. Ferullo and S. T. Lovett, PLoS genetics 4, e1000300 (2008).

[98] J. DeNapoli, A. K. Tehranchi, and J. D. Wang, Molecular microbiology 88, 93 (2013).

[99] K. Labib and B. Hodgson, EMBO reports 8, 346 (2007).

[100] H. Meinhardt and P. A. de Boer, Proceedings of the National Academy of Sciences 98, 14202 (2001).

[101] S. Arumugam, Z. Petrášek, and P. Schwille, Proceedings of the National Academy of Sciences 111, E1192 (2014).

[102] M. Osella, S. J. Tans, and M. C. Lagomarsino, Trends in microbiology 25, 250 (2017).

[103] F. Si, G. Le Treut, J. T. Sauls, S. Vadia, P. A. Levin, and S. Jun, Current Biology 29, 1760 (2019).

