## Supplemental Text and Supplemental Figures S1-S25 for "Coordination between *E. coli* Cell Size and Cell Cycle Mediated by DnaA"

### SUPPORTING TEXT

#### 1 Some details of the model

##### 1.1 The derivation of the initiation probability (1)

A series of DnaA binding sites (DnaA boxes) exists at the replication origin (*oriC*) of *E.coli* chromosome (2–5). At *oriC*, there are three widely spaced strong boxes (R1, R2, and R4) and more than five weak boxes (e.g., R3, R5M, I1, I2, I3) which are located between the strong ones (2–5). DnaA-ATP and DnaA-ADP can bind to strong boxes with a similarly high affinity (6). The weak ones independently bind DnaA-ATP or DnaA-ADP with a low affinity, but they can bind DnaA-ATP with a high affinity via the cooperation of the adjacent strong boxes bound with DnaA-ATP (1, 4, 7). In the initiation process, the strong boxes bind DnaA-ATP firstly as the anchors, and then the weak boxes bind DnaA-ATP gradually with the cooperation of adjacent DnaA-ATP (1, 4, 7). Finally, more than ten DnaA-ATPs form an ordered polymeric pre-replication complex (pre-RC), which opens the duplex unwinding element of *oriC* (3–5), and triggers the consecutive loading of replicative DNA helicase (DnaB), primase, and sliding clamp of DNA polymerase III holoenzyme on the nascent exposed single DNA strands (2, 8). To model the initiation process, we coarse-grained the wild-type *oriC* of *E.coli* (5) as a simpler version which includes three identical strong DnaA boxes and four identical weak DnaA boxes as done in Ref. (1). One weak DnaA box binds DnaA-ATP only if its adjacent strong box has bound DnaA-ATP. We denote the dissociation constant of DnaA-ATP from DnaA boxes is $K_1$ and of DnaA-ADP from strong DnaA boxes is $K_2$. The probability of all the strong DnaA boxes occupied by DnaA-ATP is $\left(1+\frac{K_1}{c_1}+\frac{K_1/K_2}{c_1/c_2}\right)^{-3}$. Given this condition, the probability of four weak DnaA boxes all occupied by DnaA-ATP is $\left(1+\frac{K_1}{c_1}\right)^{-4}$. Once all seven DnaA boxes are occupied by DnaA-ATP, more DnaA-ATP molecules will easily join to assemble the ordered multimeric pre-RC, which will then open the duplex unwinding region. Therefore, this simplified *oriC* shows that the binding of DnaA-ATP with these seven DnaA boxes determines the initiation of DNA replication. In this study, we simulate cell cycles with a mean-field method and ignore the cell-to-cell variability in the initiation process. We set the initiation condition as that *oriC* out of eclipse period is initiated when the probability of seven DnaA boxes binding DnaA-ATP (initiation probability) is equal to or higher than a threshold. The initiation probability can be expressed as (1)

$$P_{ini}=\left(1+\frac{K_1}{c_1}+\frac{K_1/K_2}{c_1/c_2}\right)^{-3}\left(1+\frac{K_1}{c_1}\right)^{-4} \quad (1)$$

##### 1.2 The derivation of DnaA synthesis rate

We derived the DnaA synthesis rate $\rho(t)$, adapting from ref. (1). Considering the weakness of DnaA promoter (9, 10), we approximately have that DnaA expression rate is proportional to the concentration of free RNAP, i.e.

$$\rho(t)\propto[RNAP]_f. \quad (2)$$

Here we assume that $[RNAP]_f$ is time (cell cycle)-independent in order to avoid more complex consideration. To consider the cell cycle dependence of $[RNAP]_f$, one may extend the model of Klumpp and Hwa (11) for RNAP partitioning. The *dnaA* gene is close to $oriC$, so the copy number of *dnaA* can be represented by the number of origins (i.e. $n_{ori}$). DnaA transcription is auto-repressed by DnaA-ATP and DnaA-ADP (12–15). Based on repression percents of DnaA-ATP and DnaA-ADP on *dnaA* promoters (*dnaAp1* and *dnaAp2*) measured by Speck et al. (15), the effect of DnaA auto-repression can be phenomenologically described by

$$\rho(t) \propto 1/\left(1 + [k_1 x_f(t)/V(t)]^2 + k_2 y_f(t)/V(t)\right), \tag{3}$$

where $k_1(= 0.024\mu m^3)$ and $k_2(= 0.0135\mu m^3)$ are binding constants of DnaA-ATP and DnaA-ADP, respectively, and the power 2 originates from DnaA-ATP self-cooperation (1). As a global regulator controlling growth rate (16–18), ppGpp also represses the expression of *dnaA* (10, 19, 20). ppGpp probably inhibits the transcription of *dnaA* by destabilizing the RNAP-promoter open complex in collaboration with DksA, or by decreasing availability of RNAP for DnaA promoter by globally altering the utilization of sigma factors (1, 16, 19, 21). So far, no regulator at the post-transcriptional level has been well characterized for DnaA synthesis. But ppGpp may globally inhibit the translation of a protein by reducing the active ribosome fraction at small growth rates (18). As commonly done for a complex effect with unclear details, we represent the effect of ppGpp inhibition on DnaA expression simply by a Hill function

$$\rho(t) \propto K_p^n/(K_p^n + [ppGpp]^n) \triangleq G([ppGpp]), \tag{4}$$

where $K_p$ and $n$ denote equilibrium constant and Hill coefficient respectively, which are determined by fitting experimental data. Just like dealing with free RNAP concentration, we ignore the time (cell cycle) dependence of $[ppGpp]$ and $G([ppGpp])$. Finally, we can express the DnaA synthesis rate ($\rho(t)$) as

$$\rho(t) = \frac{\alpha n_{ori}(t)[RNAP]_f G([ppGpp])}{1 + [k_{f,1} x_f(t)/V(t)]^2 + k_{f,2} y_f(t)/V(t)}, \tag{5}$$

where $\alpha$ is the synthesis rate constant.

#### 1.3 The derivation of DnaA inactivation rate

Let $n_{box}$, $n_{box,1}$, and $n_{box,2}$ denote the number of nonfunctional DnaA boxes, of those at $datA$ and of others elsewhere, respectively. As done previously (1), we assume nonfunctional DnaA boxes not at $datA$ homogeneously distribute on the chromosome, which is duplicated along with the replication. At the initiation of DNA replication ($t_{ini}$), replication origin number $n_{ori}$ is doubled and fork number $n_{fork}$ increases by $2n_{ori}(t_{ini}^-)$, while, at the termination ($n_{ori} + C$), $n_{fork}$ decreases by $2n_{ori}(t_{ini}^-)$. $n_{box,1}$ is proportional to copy number of $datA$, while $n_{box,1}$ is proportional to the genome content. Therefore, the change in $n_{fork}$ and $n_{ori}$ with the time can be formulated as

$$dn_{ori}/dt = n_{ori}(t_{ini}^-)\delta(t - t_{ini}), \tag{6}$$

$$dn_{fork}/dt = 2n_{ori}(t_{ini}^-)\left[\delta(t - t_{ini}) - \delta(t - t_{ini} - C)\right], \tag{7}$$

where $t_{ini}$ denotes initiation time of DNA replication, $t_{ini}^-$ denotes the time approaching $t_{ini}$ from left, and $\delta(x)$ denotes the Dirac delta function (1). Accordingly, the dynamics of $n_{box}$, $n_{box,1}$, and $n_{box,2}$ are given by (1)

$$n_{box,1}(t) = n_{box,1}^* \cdot n_{ori}(t - x_{datA}C), \tag{8}$$

$$dn_{box,2}(t)/dt = n_{box,2}^* \cdot n_{fork}(t)/(2C), \tag{9}$$

$$n_{box}(t) = n_{box,1}(t) + n_{box,2}(t). \tag{10}$$

where $n_{box,1}^*$ and $n_{box,2}^*$ denote the numbers of DnaA boxes per copy of *datA* and per chromosome excluding the *datA* region, respectively, and $x_{datA} = 0.2$, indicating the distance from *datA* to *oriC* divided by the chromosomal half-length (notice that the distance was assumed to be zero, i.e. $x_{datA} = 0$ in our previous model).

Hda protein, by forming a complex with $\beta$-subunit clamp of DNA polymerase III holoenzyme sliding on DNA, promotes the hydrolysis of ATP bound to DnaA along with chromosome replication, which is called "regulatory inactivation of DnaA (RIDA)" (22–25). Accordingly, we can assume that DnaA-ATPs bound at one chromosomal site are all inactivated into DnaA-ADPs when this chromosomal site is replicated. We use $\epsilon(t)$ to denote the hydrolysis rate of DnaA-ATP. Then the number of the bound DnaA-ATP hydrolyzed during the time between $t$ and $t + dt$ is $\epsilon(t)dt$. According to the above assumption, the number of bound DnaA-ATPs which are hydrolyzed during the period

$[t, t+dt]$ is equal to the number of bound DnaA-ATPs which are passed by the replication forks during $[t, t+dt]$. The nonfunctional DnaA boxes are assumed to have the same binding affinity with DnaA. So the number of bound DnaA-ATPs which are passed by replication forks during $[t, t+dt]$ is proportional to the number of DnaA boxes which are passed by replication forks during $[t, t+dt]$, in which the scale factor is $x_b(t)/n_{box}(t)$. The number of DnaA boxes passed by replication forks is doubled, so the change in the number of DnaA boxes during $[t, t+dt]$ (i.e. $dn_{box}$) is equal to the number of DnaA boxes passed by replication forks during $[t, t+dt]$. Therefore, $\epsilon(t)dt = x_b(t)/n_{box}(t)dn_{box}$, i.e., $\epsilon(t) = x_b(t)/n_{box}(t)dn_{box}/dt$. From equations 6-10, the inactivation rate of DnaA-ATP ($\epsilon(t)$) can be further represented by

$$\begin{aligned}\epsilon(t) &= \frac{x_b}{n_{box}}\left(\frac{dn_{box,1}}{dt} + \frac{dn_{box,2}}{dt}\right)\\ &= \frac{x_b}{n_{box}}\left[n^*_{box,1}\cdot\frac{dn_{ori}(t - x_{datA}C)}{dt} + \frac{1}{2C}n^*_{box,2}n_{fork}(t)\right]\\ &= \frac{x_b}{n_{box}}\left[n^*_{box,1}n_{ori}(t^-_{ini})\delta(t - x_{datA}C - t_{ini}) + \frac{1}{2C}n^*_{box,2}n_{fork}(t)\right]. \qquad (11)\end{aligned}$$

#### 1.4 The possible effect of SeqA on DnaA transcription and titration (1)

The *dnaA* promoter also contains some GATC sites, so SeqA may also take a role in the regulation of DnaA transcription during the eclipse period (26). In addition, SeqA probably affects DnaA titration by sequestering the hemimethylated region containing DnaA boxes, since SeqA acts in organizing newly duplicated DNA behind replication forks (27). The experiments of *seqA* knockout and GATC sites elimination inside the *dnaA* promoter showed that SeqA does not affect replication initiation much (28, 29). For simplicity, therefore, we neglected the possible repression of SeqA on DnaA transcription and titration.

#### 1.5 Comparing the results of linear and exponential growth patterns

Our simulation with linear and exponential growth patterns produced different initiation mass, but nearly the same unit cell size, average cell size, origin number, genome content, and DnaA level. These are in line with the literature (1, 30–32). Eq. 1 in the main text, which connects cell mass at division to initiation mass, depends on the assumption of the exponential growth model (30). However, Eq. 2 in the main text, which describes cell size as the sum of unit cells, independent of the growth model (32). The derivation of theoretical formulas (see below) expressing origin number and genome content as a function of $C$, $D$, and $\tau$ is also independent of the growth pattern (31). Earlier simulation of DnaA-mediating DNA replication also showed that linear and exponential growth patterns do not bring a significant difference in the DnaA level (1).

#### 1.6 Simulation and data fitting

According to the model, we simulated the evolution of levels of different forms of DnaA along with the cell cycle. Below are some details of the simulation, similar as in (1). In the simulation, we applied the Euler forward algorithm with time steps $\Delta t = 1s$ for Eqs. 11-12 in the main text and $\Delta t_s = 0.01s$ for Eqs. 13-16 in the main text. The usage of smaller time steps or fourth-order Runge-Kutta algorithm did not change the results much. Arbitrary initial values were assigned to $x$, $y$, $x_f$, $y_f$, and cell birth volume $V_0$. The initial DNA content $Gc$ was chosen as one genome equivalent. Initial values of other variables were: $n_{ori}(0) = 1$, $n_{fork}(0) = 0$, $n_{box,1}(0) = n^*_{box,1}$, $n_{box,2}(0) = n^*_{box,2}$ and $n_{box}(0) = n^*_{box,1} + n^*_{box,2}$. Cell volume increased exponentially with the time by $V(t) = V_0 2^{t/\tau}$. DNA replication was initiated when the initiation probability $P_{ini}$ goes up to the threshold (typically 0.3) after the eclipse period. Cell division happened after a time of $C + D$ from the initiation of DNA replication. At the cell division, all the extensive variables, including $x_f$, $y_f$, $x$, $y$, $n_{box}$, $n_{box,1}$, $n_{box,2}$, $n_{ori}$, $n_{fork}$ and $Gc$, for the new-born daughter cell were reduced to one half of that for the divided mother cell. We updated the cell birth volume ($V_0$) for a new cell cycle based on the relative deviation of the simulated inter-division time and the predetermined doubling time ($\tau$) by $V_0^{(k+1)} = V_0^{(k)} e^{\gamma(T_k - \tau)/\tau}$, where $T_k$ is the inter-division time of generation $k$, $V_0^{(k)}$ and $V_0^{(k+1)}$ are the birth volume of generations $k$ and $k+1$, respectively and $\gamma = 0.1$ (typically). The parameter $\gamma$ in the above update formula affects the convergence speed and precision. We tested with different forms of the update formula and did not find any difference in the main results. The simulation stopped when the relative difference between the inter-division (inter-initiation) time and the preset doubling time was smaller than a threshold or the preset run time (no less than $5 \times 10^6$s) was over. In the former case, a regular DnaA oscillation coupled with cell cycle and a converged birth volume will be achieved. In the latter case, an irregular DnaA oscillation and an abnormal cell cycle will be obtained.

In the simulation, $C$, $D$, and doubling time $\tau$ were determined by averaging their experimental values. The values of other parameters were obtained directly from the literature or determined by fitting data in the literature with the least-squares method. Values of parameters directly from the literature include: $k_1 = 0.024\mu m^3$, $k_2 = 0.0135\mu m^3$, $K_1 = K_2 = 1\mu m^{-3}$, and initiation threshold =0.3 (1, 6, 33); $n^*_{box,1} = 350$ (1, 34), $n^*_{box,2} = 300$ (1, 35, 36); $k_{on,1} = 3 \times 10^{-5}\mu m^3/s$, $k_{on,2} = 2 \times 10^{-5}\mu m^3/s$, $k_{off,1} = k_{off,2} = 0.003s^{-1}$ (1, 33), $x_{datA} = 0.2$ (34). By fitting the dependence of the concentrations of ppGpp and free RNAP on growth rate from Ref. (11, 37), we obtained $[RNAP]_{f,0} = 883\mu m^{-3}$, $\mu_r = 0.68$ dbls/h, $[ppGpp]_0 = 92$ pmol/$OD_{460}$, and $\mu_p = 1.11$ dbls/h (see Fig. S1). The remaining parameters were fixed by fitting experimental data on the relationship between cell size and the perturbed parameter (growth rate, $C$ or $D$) in the literature (32, 38, 39). Note that the fits on cell size were done separately for each set of experimental data of perturbing growth rate, C period, or D period, because some relevant parameters depend on strains and growth conditions used in the experiment. First, we fitted experimental data of Si et al. (32) under Rep knockdown and obtained $\beta = 2 \times 10^{-3}s^{-1}$ (common parameter) and $\alpha' = 0.125\mu m^3/s$ (case-dependent parameter). With the same $\beta$, then, we obtained: $\alpha' = 0.5$, $0.12$, and $0.55\mu m^3/s$ by fitting experimental data of Si et al. (32) under hydroxyurea induction, SulA overexpression, and cephalexin induction respectively; $\alpha' = 0.4\mu m^3/s$ by fitting experimental data of Zhu et al. (39) under ribonucleotide reductase titration; $\alpha' = 0.105$ (RDM + glycerol ) or $0.28\mu m^3/s$ (RDM + glucose) by fitting experimental data of Zheng et al. (38) under mreB/ftsZ-titration; $\alpha = 2.4 \times 10^{-3}\mu m^3/s$, $K_p = 11\mu m^{-3}$, $n = 2.9$ by fitting experimental data of Si et al. (32) under various nutrient conditions (these parameters were also used for perturbations of initiation mass $S_0$).

With proper parameters, we obtained a regular cell cycle, accompanying with periodic oscillations of cell cycle variables, for example, the number of DnaA molecules, the number of replication origins, genome content, and cell volume, when the steady state emerges in the simulation. The population average of a cell cycle variable derived with the cell age distribution ($g(t) = (2\ln 2/\tau) \cdot 2^{-t/\tau}$) can be expressed as a function of the perturbed parameter. We use $S$ to denote the average cell size (volume) i.e. $S = \langle V(t) \rangle$, $S_0$ to denote the average cell size per average origin (i.e. unit cell size ), and $S_b$ to denote the cell size at birth. At the exponential steady state, cell size and other cell cycle parameters for bacteria have some quantitative relationships (31, 32, 40):

1. The average cell size is proportional to the birth cell size (32), i.e., $S = 2\ln 2 \cdot S_b$.

2. The unit cell size is equal to the initiation mass (size) times $\ln 2$ (32), i.e., $S_0 = \ln 2 \cdot S_i$.

3. Origin number is a function of DNA replication duration ($C$), cell division duration ($D$) and doubling time ($\tau$) (31), i.e., $n_{ori} = 2^{(C+D)/\tau}$.

4. The genome content is a function of $C$, $D$, and $\tau$ (40), i.e., $Gc = [\tau/(C \cdot \ln 2)][2^{(C+D)/\tau} - 2^{D/\tau}]$.

Above formulas can be also viewed as criterions for achieving normal cell cycles.

### SUPPORTING REFERENCES


1. Zhang, Q., and H. Shi, 2012. Coupling chromosomal replication to cell growth by the initiator protein DnaA in Escherichia coli. *Journal of theoretical biology* 314:164–172.

2. Messer, W., 2002. The bacterial replication initiator DnaA. DnaA and oriC, the bacterial mode to initiate DNA replication [Review]. *Fems Microbiology Reviews* 26:355–74.

3. Leonard, A. C., and J. E. Grimwade, 2005. Building a bacterial orisome: emergence of new regulatory features for replication origin unwinding. *Molecular Microbiology* 55:978–85.

4. Leonard, A. C., and J. E. Grimwade, 2011. Regulation of DnaA Assembly and Activity: Taking Directions from the Genome. *Annual Review of Microbiology* 65:19.

5. Miller, D. T., J. E. Grimwade, T. Betteridge, T. Rozgaja, J. C. Torgue, and A. C. Leonard, 2009. Bacterial Origin Recognition Complexes Direct Assembly of Higher-Order DnaA Oligomeric Structures. *Proceedings of the National Academy of Sciences of the United States of America* 106:18479.

6. Sekimizu, K., D. Bramhill, and A. Kornberg, 1987. ATP activates dnaA protein in initiating replication of plasmids bearing the origin of the E. coli chromosome. *Cell* 50:259–265.

7. Leonard, A. C., and J. E. Grimwade, 2010. Regulating DnaA complex assembly: it is time to fill the gaps. *Current Opinion in Microbiology* 13:766.

8. Fang, L., M. J. Davey, and M. O'Donnell, 1999. Replisome assembly at oriC, the replication origin of E. coli, reveals an explanation for initiation sites outside an origin. *Molecular Cell* 4:541.

9. Hansen, F. G., E. B. Hansen, and T. Atlung, 1982. The nucleotide sequence of the dnaA gene promoter and of the adjacent rpmH gene, coding for the ribosomal protein L34, of Escherichia coli. *The EMBO journal* 1:1043–1048.

10. Saggioro, C., A. Olliver, and B. Sclavi, 2013. Temperature-dependence of the DnaA–DNA interaction and its effect on the autoregulation of dnaA expression. *Biochemical Journal* 449:333–341.

11. Klumpp, S., and T. Hwa, 2008. Growth-rate-dependent partitioning of RNA polymerases in bacteria. *Proceedings of the National Academy of Sciences* 105:20245–20250.

12. Atlung, T., E. S. Clausen, and F. G. Hansen, 1985. Autoregulation of the dnaA gene of Escherichia coli K12. *Molecular and General Genetics MGG* 200:442–450.

13. Braun, R. E., K. O'Day, and A. Wright, 1985. Autoregulation of the DNA replication gene dnaA in E. coli K-12. *Cell* 40:159–169.

14. Kücherer, C., H. Lother, R. Kölling, M.-A. Schauzu, and W. Messer, 1986. Regulation of transcription of the chromosomal dnaA gene of Escherichia coli. *Molecular and General Genetics MGG* 205:115–121.

15. Speck, C., C. Weigel, and W. Messer, 1999. ATP–and ADP–DnaA protein, a molecular switch in gene regulation. *The EMBO journal* 18:6169–6176.

16. Srivatsan, A., and J. D. Wang, 2008. Control of bacterial transcription, translation and replication by (p)ppGpp. *Current Opinion in Microbiology* 11:100.

17. Potrykus, K., H. Murphy, N. Philippe, and M. Cashel, 2011. ppGpp is the major source of growth rate control in E. coli. *Environmental microbiology* 13:563–575.

18. Dai, X., M. Zhu, M. Warren, R. Balakrishnan, V. Patsalo, H. Okano, J. R. Williamson, K. Fredrick, Y.-P. Wang, and T. Hwa, 2016. Reduction of translating ribosomes enables Escherichia coli to maintain elongation rates during slow growth. *Nature microbiology* 2:1–9.

19. Chiaramello, A. E., and J. W. Zyskind, 1990. Coupling of DNA replication to growth rate in Escherichia coli: a possible role for guanosine tetraphosphate. *Journal of Bacteriology* 172:2013.

20. Zyskind, J. W., and D. W. Smith, 1992. DNA replication, the bacterial cell cycle, and cell growth. *Cell* 69:5–8.

21. Traxler, M. F., S. M. Summers, H.-T. Nguyen, V. M. Zacharia, G. A. Hightower, J. T. Smith, and T. Conway, 2008. The global, ppGpp-mediated stringent response to amino acid starvation in Escherichia coli. *Molecular microbiology* 68:1128–1148.

22. Katayama, T., T. Kubota, K. Kurokawa, E. Crooke, and K. Sekimizu, 1998. The initiator function of DnaA protein is negatively regulated by the sliding clamp of the E. coli chromosomal replicase. *Cell* 94:61–71.

23. Kato, J.-i., and T. Katayama, 2001. Hda, a novel DnaA-related protein, regulates the replication cycle in Escherichia coli. *The EMBO journal* 20:4253–4262.

24. Su'etsugu, M., T.-r. Shimuta, T. Ishida, H. Kawakami, and T. Katayama, 2005. Protein associations in DnaA-ATP hydrolysis mediated by the Hda-replicase clamp complex. *Journal of Biological Chemistry* 280:6528–6536.

25. Kim, J. S., M. T. Nanfara, S. Chodavarapu, K. S. Jin, V. M. Babu, M. A. Ghazy, S. Chung, J. M. Kaguni, M. D. Sutton, and Y. Cho, 2017. Dynamic assembly of Hda and the sliding clamp in the regulation of replication licensing. *Nucleic acids research* 45:3888–3905.

26. Campbell, J. L., and N. Kleckner, 1990. E. coli oriC and the dnaA gene promoter are sequestered from dam methyltransferase following the passage of the chromosomal replication fork. *Cell* 62:967–979.

27. Løbner-Olesen, A., O. Skovgaard, and M. G. Marinus, 2005. Dam methylation: coordinating cellular processes. *Current opinion in microbiology* 8:154–160.

28. Camara, J. E., A. M. Breier, T. Brendler, S. Austin, N. R. Cozzarelli, and E. Crooke, 2005. Hda inactivation of DnaA is the predominant mechanism preventing hyperinitiation of Escherichia coli DNA replication. *EMBO reports* 6:736–741.

29. Wilkinson, T. G., G. Kedar, C. Lee, E. C. Guzmán, D. W. Smith, and J. W. Zyskind, 2006. The synchrony phenotype persists after elimination of multiple GATC sites from the dnaA promoter of Escherichia coli. *Journal of bacteriology* 188:4573–4576.

30. Donachie, W. D., 1968. Relationship between cell size and time of initiation of DNA replication. *Nature* 219:1077–1079.

31. Bremer, H., and G. Churchward, 1977. An examination of the Cooper-Helmstetter theory of DNA replication in bacteria and its underlying assumptions. *Journal of Theoretical Biology* 69:645–654.

32. Si, F., D. Li, S. E. Cox, J. T. Sauls, O. Azizi, C. Sou, A. B. Schwartz, M. J. Erickstad, Y. Jun, X. Li, et al., 2017. Invariance of initiation mass and predictability of cell size in Escherichia coli. *Current Biology* 27:1278–1287.

33. Schaper, S., and W. Messer, 1995. Interaction of the initiator protein DnaA of Escherichia coli with its DNA target. *Journal of Biological Chemistry* 270:17622–17626.

34. Kitagawa, R., H. Mitsuki, T. Okazaki, and T. Ogawa, 1996. A novel DnaA protein-binding site at 94.7 min on the Escherichia coli chromosome. *Molecular microbiology* 19:1137–1147.

35. Roth, A., and W. Messer, 1998. High-affinity binding sites for the initiator protein DnaA on the chromosome of Escherichia coli. *Molecular microbiology* 28:395–401.

36. Ogawa, T., Y. Yamada, T. Kuroda, T. Kishi, and S. Moriya, 2002. The datA locus predominantly contributes to the initiator titration mechanism in the control of replication initiation in Escherichia coli. *Molecular microbiology* 44:1367–1375.

37. Bremer, H., and P. P. Dennis, 1996. Escherichia coli and Salmonella: Cellular and Molecular Biology, 2nd ed., American Society for Microbiology, Washington, DC, USA, chapter Modulation of chemical composition and other parameters of the cell by growth rate, 1553–1569.

38. Zheng, H., P. Y. Ho, M. Jiang, B. Tang, W. Liu, D. Li, X. Yu, N. E. Kleckner, A. Amir, and C. Liu, 2016. Interrogating the Escherichia coli cell cycle by cell dimension perturbations. *Proc Natl Acad Sci U S A* 113:15000–15005.

39. Zhu, M., X. Dai, W. Guo, Z. Ge, M. Yang, H. Wang, and Y. P. Wang, 2017. Manipulating the Bacterial Cell Cycle and Cell Size by Titrating the Expression of Ribonucleotide Reductase. *Mbio* 8.

40. Cooper, S., and C. E. Helmstetter, 1968. Chromosome replication and the division cycle of Escherichia coli Br. *Journal of molecular biology* 31:519–540.

41. Zhang, Q., E. Brambilla, R. Li, H. Shi, M. C. Lagomarsino, and B. Sclavi, 2019. A decrease in transcription capacity limits growth rate upon translation inhibition. *bioRxiv* doi: 10.1101/599183 (preprint posted October 09, 2019).

42. Mahaffy, J. M., and J. W. Zyskind, 1989. A model for the initiation of replication in Escherichia coli. *Journal of theoretical biology* 140:453–477.

43. Browning, S. T., M. Castellanos, and M. L. Shuler, 2004. Robust control of initiation of prokaryotic chromosome replication: essential considerations for a minimal cell. *Biotechnology and bioengineering* 88:575–584.

44. Hansen, F. G., T. Atlung, R. Braun, A. Wright, P. Hughes, and M. Kohiyama, 1991. Initiator (DnaA) protein concentration as a function of growth rate in Escherichia coli and Salmonella typhimurium. *Journal of Bacteriology* 173:5194–5199.

45. Chiaramello, A. E., and J. W. Zyskind, 1989. Expression of Escherichia coli dnaA and mioC genes as a function of growth rate. *Journal of bacteriology* 171:4272–4280.

46. Grant, M. A., C. Saggioro, U. Ferrari, B. Bassetti, B. Sclavi, and M. C. Lagomarsino, 2011. DnaA and the timing of chromosome replication in Es-cherichia coli as a function of growth rate. *BMC systems biology* 5:201.

### SUPPORTING FIGURES

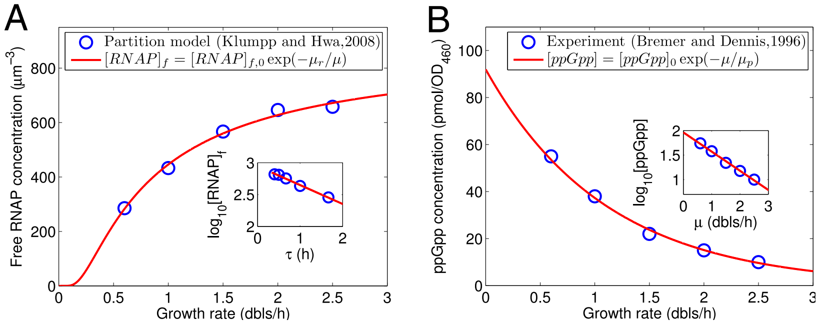


Figure S1: The dependence of concentrations of free RNAP and ppGpp on the growth rate (or doubling time) can be captured by two exponential functions. (A) Free RNAP concentration ($[RNAP]_f$) can be expressed as a function of growth rate ($\mu$), i.e. $[RNAP]_f = [RNAP]_{f,0}\exp(-\mu_r/\mu)$, based on the data of Klumpp and Hwa (11). The formula indicates an exponential dependence of $[RNAP]_f$ on doubling time ($\tau = 1/\mu$). By linearly fitting the data of $\log_{10}[RNAP]_f$ as a function of $\tau$ (inset), we obtained $[RNAP]_{f,0} = 883\mu m^{-3}$ and $\mu_r = 0.68$ dbls/h. (A) ppGpp concentration ($[ppGpp]$) can be expressed as an exponential function of growth rate ($\mu$), i.e. $[ppGpp] = [ppGpp]_0\exp(-\mu/\mu_p)$, based on experimental data of Bremer and Dennis (37) (also see Ref. (41)). By linearly fitting experimental data of $\log_{10}[ppGpp]$ as a function of $\mu$ (inset), we obtained $[ppGpp]_0 = 92\ pmol/OD_{460}$ and $\mu_p = 1.11$ dbls/h.

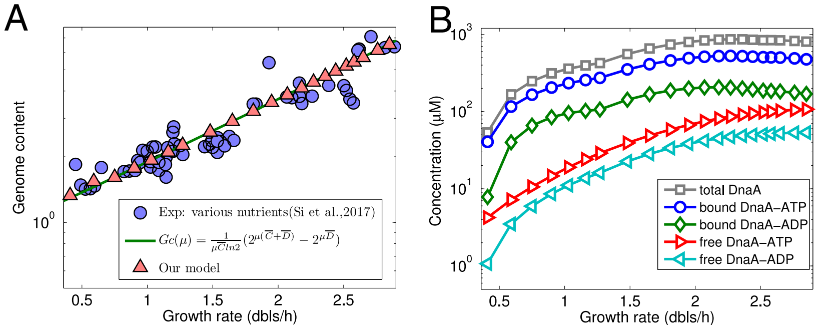


Figure S2: Our model quantifies the change in genome content and DnaA concentrations with the growth rate. The parameters used here are the same as those for Fig. 2 in the main text. (A) The simulated genome content increases with growth rate, exactly as the theoretical formula and in line with experimental data. (B) The predicted concentration of DnaA in each form basically increases with growth rate and the faster increase at the smaller growth rate.

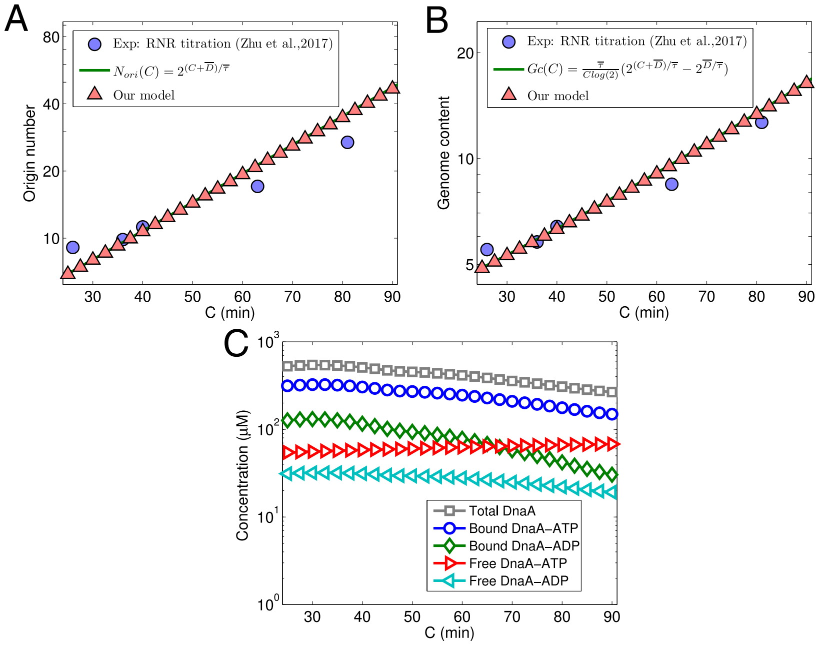


Figure S3: Our model quantifies the change in the origin number, genome content, and DnaA concentration with prolonged C period ($C$) based on the data of Zhu et al. (39) for ribonucleotide reductase (RNR) titration. The parameters used here are the same as those for Fig. 3 in the main text. (A,B) Origin number and genome content (per cell) from the simulation increase with $C$ exactly following theoretical formulas and in line with experimental data. (C) The predicted concentration of DnaA in each form decreases with $C$.

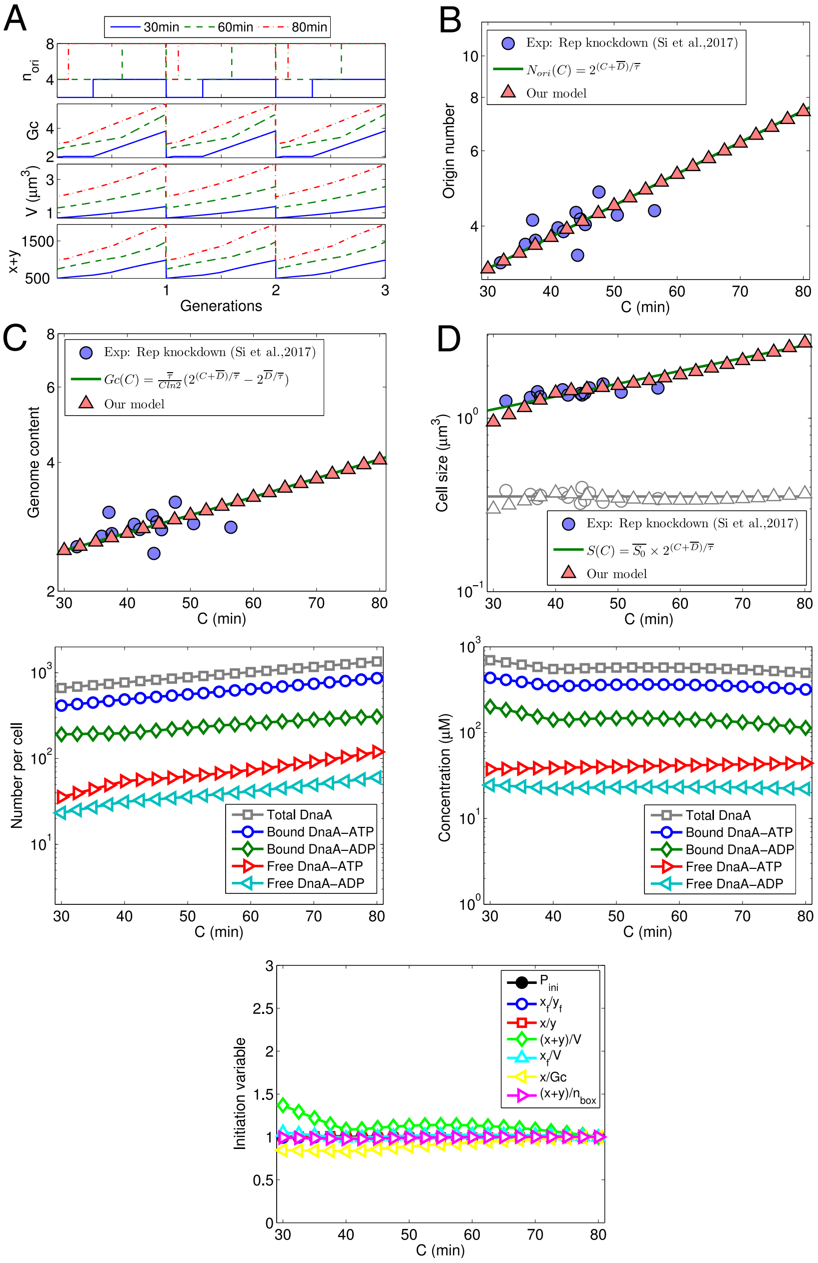


Figure S4: Our model quantifies the change in the origin number, genome content, and the concentration of each form of DnaA with the changing duration of C period ($C$) based on experimental data of Si et al. (32) for DNA helicase Rep repression. The experimental averages ($\overline{\tau}$=40.7 min, $\overline{D}$=38 min) were assigned to the doubling time and $D$ for the simulation. By fitting experimental data on cell size as a function of $C$, we have $\beta = 2 \times 10^{-3} s^{-1}$ (also used for all other perturbations) and $\alpha' = 0.125 \mu m^3/s$. Other parameters are pointed out in the text of Supporting Material. (A) Regular oscillations of origin number, genome content, DnaA level, and cell size are coupled with the cell cycle in the simulated steady states of different $C$. (B-C) The simulated origin number and genome content per cell increase with C period, exactly following theoretical formulas and in line with experimental data. (D) Obtained by fitting experimental data, cell size increases with $C$ roughly as the replication growth law. (E) The predicted number of DnaA in each form increases with $C$. (F) From $C = 30$ min to $C = 80$ min, the predicted concentration of total DnaA, bound DnaA-ATP, bound DnaA-ADP, or free DnaA-ADP is decreased slightly, while that of free DnaA-ATP is increased a little. (G) With prolonged C period, the variables $P_{ini}$, $x_f/y_f$, $x/y$, $x_f/V$, and $(x+y)/n_{box}$ at initiation of DNA replication change little, $x/Gc$ at initiation increases slowly, $(x+y)/V$ at initiation decreases slowly. Each initiation variable was divided by the value at $C = 80$ min.

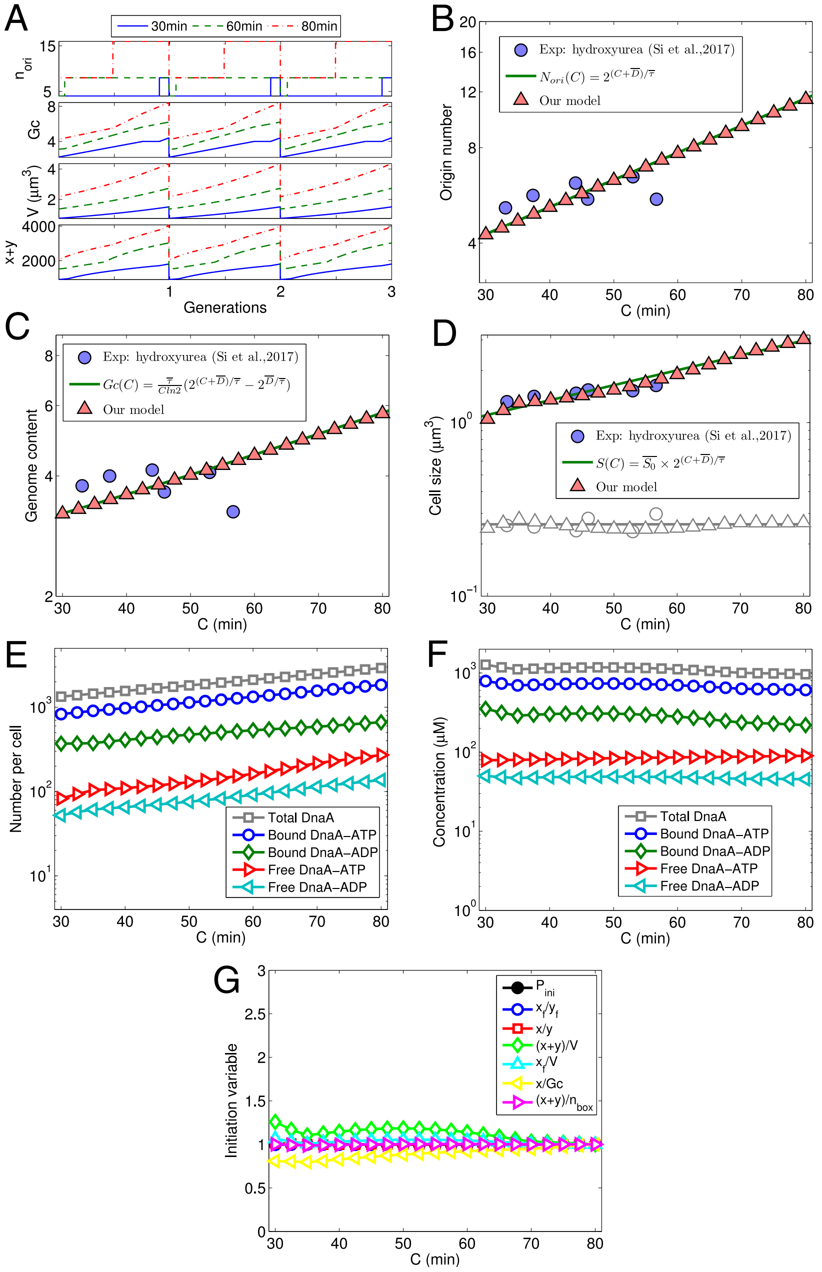


Figure S5: Our model quantifies the change in the origin number, genome content, cell size, DnaA levels and initiation variables with prolonged C period ($C$) based on experimental data of Si et al. (32) for hydroxyurea induction. The experimental averages ($\overline{\tau}$ =35.1 min, $\overline{D}$ =43.6 min) were assigned to doubling time and $D$ for the simulation. By fitting experimental data on cell size as a function of $C$, we have $\alpha' = 0.5\mu m^3/s$. Other parameters are pointed out in the Supporting Text. (A) Regular oscillations of origin number, genome content, DnaA level, and cell size are coupled with the cell cycle in the simulated steady states of different $C$. (B,C) The simulated origin number and genome content per cell increase with C period, exactly following theoretical formulas and in line with experimental data. (D) Obtained by fitting experimental data, cell size increases with $C$ basically as the replication growth law. (E) The predicted number of DnaA in each form increases with $C$. (F) From $C = 30$ min to $C = 80$ min, the predicted concentration of total DnaA, bound DnaA-ATP, bound DnaA-ADP, or free DnaA-ADP is decreased slightly, while that of free DnaA-ATP is increased a little. (G) With prolonged C period, the variables $P_{ini}$, $x_f/y_f$, $x/y$, $x_f/V$, and $(x+y)/n_{box}$ at initiation of DNA replication change little, $x/Gc$ at initiation increases slowly, $(x+y)/V$ at initiation decreases slowly. Each initiation variable was divided by the value at $C = 80min$. The variables at initiation of DNA replication change little with $C$.

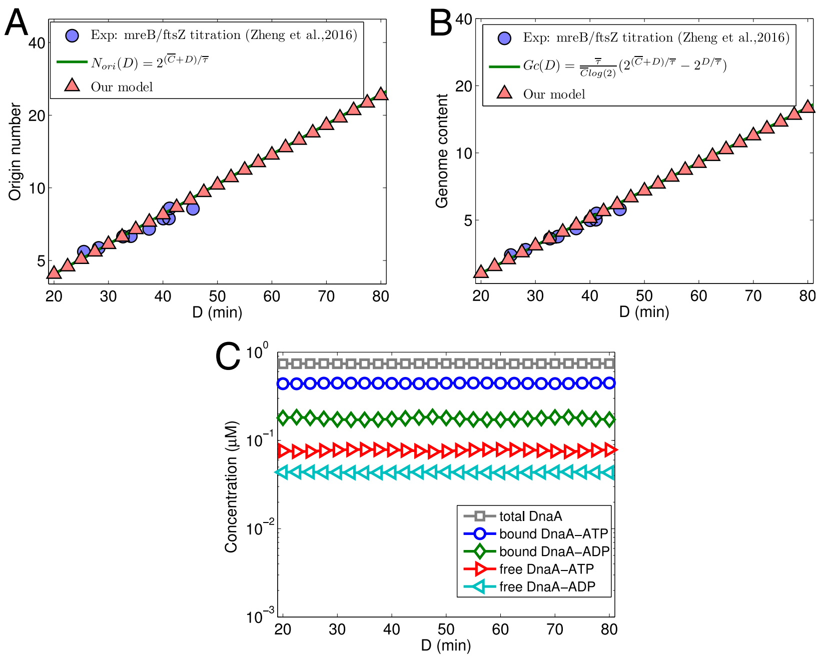


Figure S6: Our model quantifies the change in the origin number, genome content, cell size, DnaA levels and initiation variables with the prolonged D period ($D$) based on experimental data of Zheng et al. (38) for mreB/ftsZ titration (growth medium: RDM + glucose). The parameters used here are the same as those for Fig. 4 in the main text. (A-B) From the simulation, origin number and genome content per cell increase with $D$, exactly following the theoretical formula and in line with experimental data. (C) The predicted concentrations of total DnaA, bound DnaA-ATP and bound DnaA-ADP, free DnaA-ATP and free DnaA-ADP are invariant with $D$.

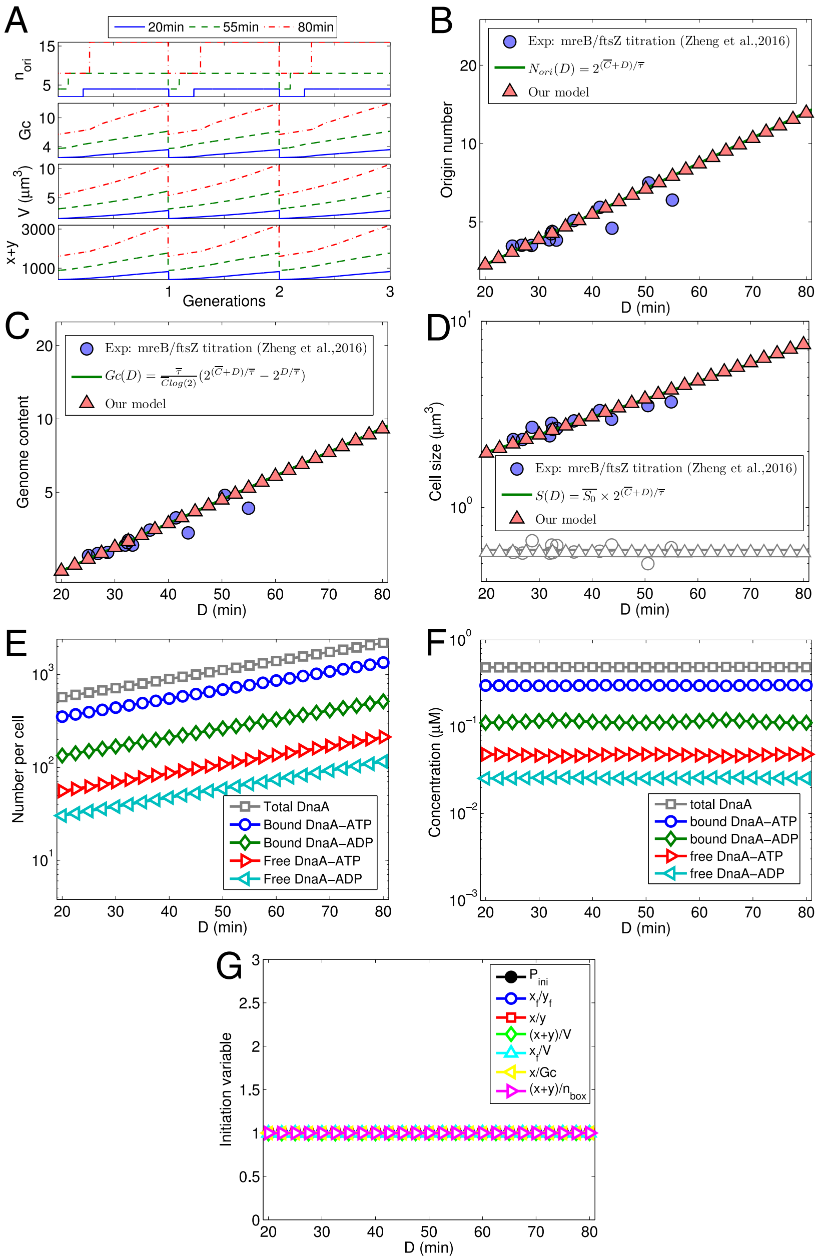


Figure S7: Our model quantifies the change in the origin number, genome content, cell size, DnaA levels and initiation variables with the prolonged D period based on experimental data of Zheng et al. (38) for mreB/ftsZ titration with growth medium RDM + glycerol. In the simulation, doubling time and C period were assigned the average experimental values ($\overline{\tau}$=30.8 min, $\overline{C}$=34.7 min). By fitting experimental data on cell size as a function of D period, we have $\alpha' = 0.105 \mu m^3/s$. Other parameters are present in the text of Supporting Material. (A) Regular oscillations of origin number, genome content, total DnaA number, and cell size coupled with cell cycles were obtained in the simulated steady states with different D periods. (B-C) From the simulation, origin number and genome content per cell increase with C period, exactly following the theoretical formula and in line with experimental data. (D) The fitted cell size exponentially increases D period, just as the division growth law. (E) The predicted number of DnaA in each form exponentially increases with D period. (F) The predicted concentration of DnaA in each form is constant. (G) The variables at initiation of DNA replication do not change with D period. Each initiation variable was normalized by dividing the value at $D = 80$ min.

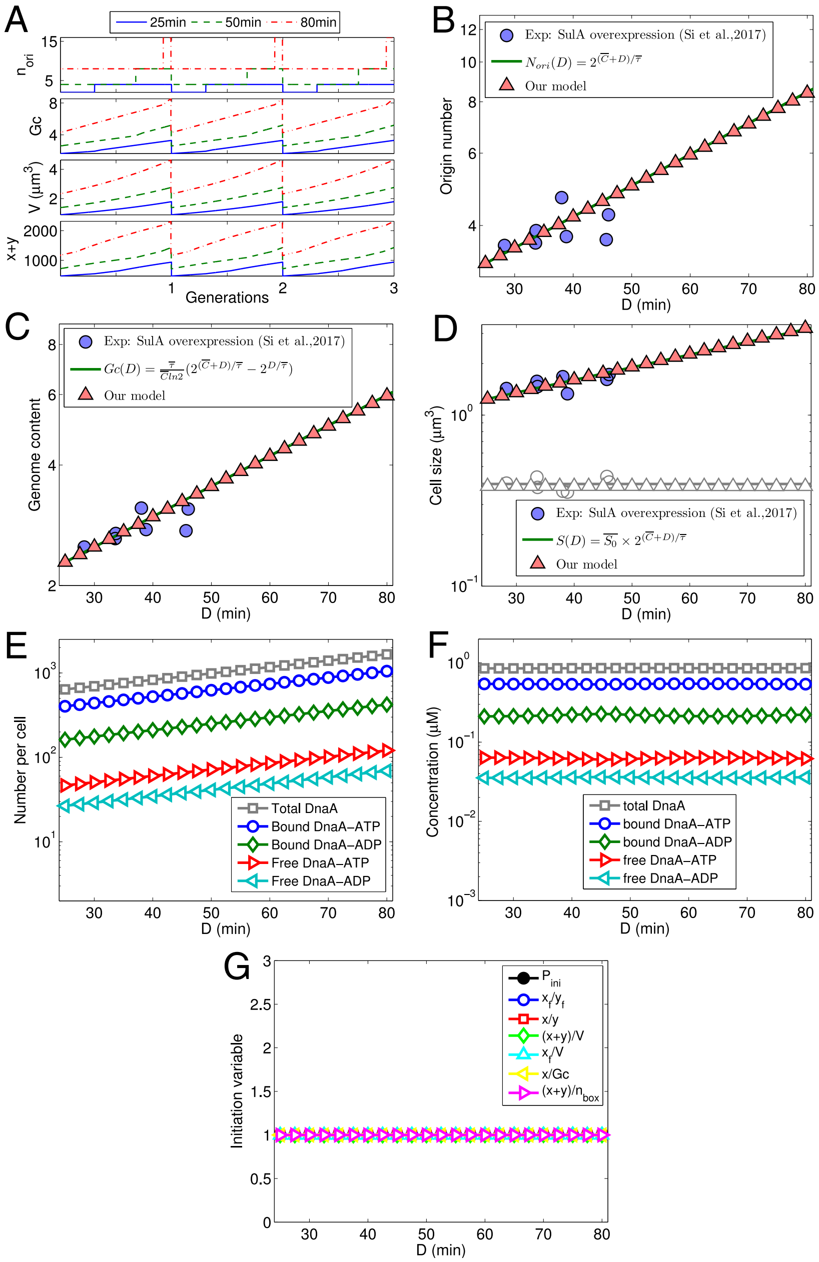

Figure S8: Our model quantifies the change in the origin number, genome content, DnaA levels and cell size with perturbed D period ($D$) based on experimental data for SulA overexpression (32). In the simulation, doubling time and $C$ were assigned the average experimental values ($\overline{\tau}$=40.7 min, $\overline{C}$=42.2 min). By fitting experimental data on cell size as a function of $D$, we have $\alpha' = 0.12\mu m^3/s$. Other parameters are present in the Supporting Text. (A) Regular oscillations of origin number, genome content, total DnaA number, and cell size coupled with cell cycles were obtained in the simulated steady states with different $D$. (B-C) From the simulation, origin number and genome content per cell increase with C period, exactly following the theoretical formula and in line with experimental data. (D) The fitted cell size exponentially increases with $D$, just as the division growth law. (E) The predicted number of DnaA in each form exponentially increases with $D$. (F) The predicted concentration of DnaA in each form is constant. (G) The variables at initiation of DNA replication do not change with $D$. Each initiation variable was normalized by dividing the value at $D = 80min$.

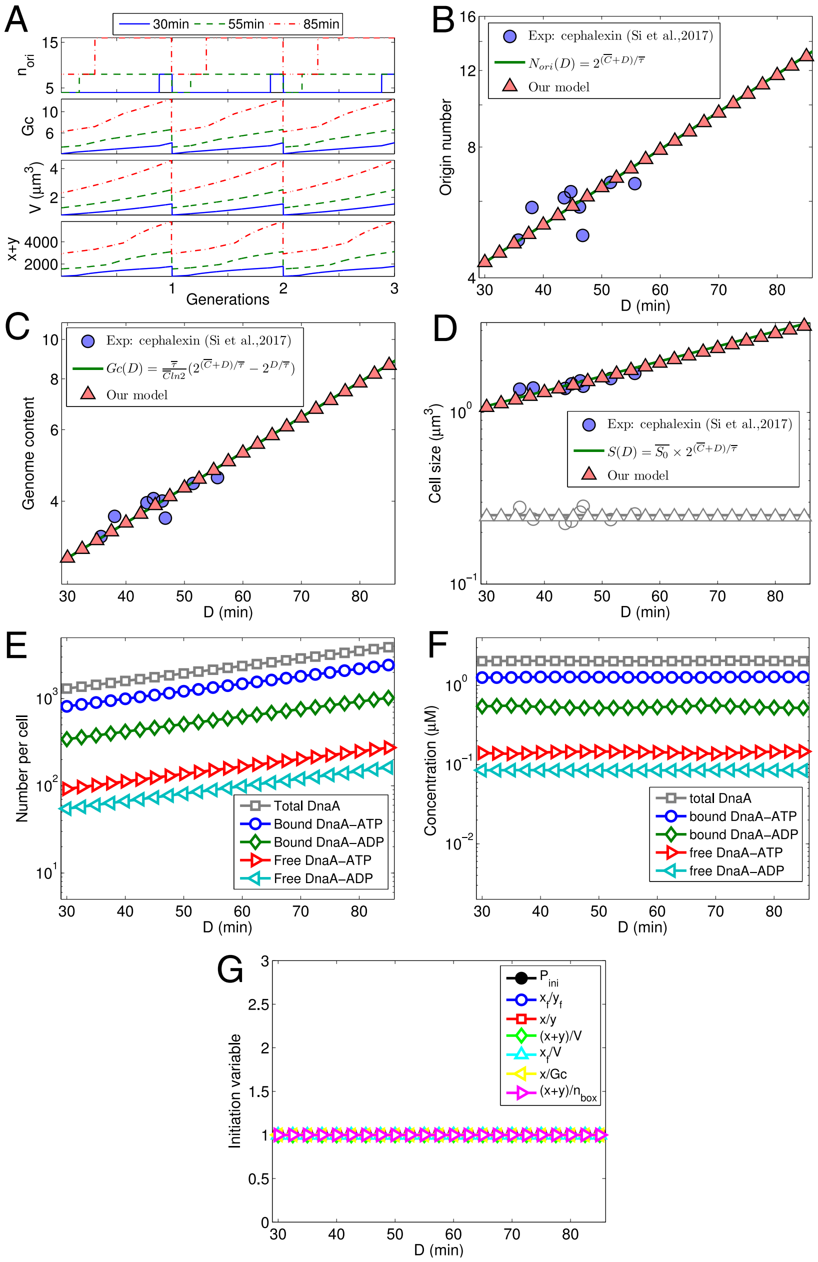


Figure S9: Our model quantifies the change in the origin number, genome content, cell size, DnaA levels and initiation variables with perturbed D period ($D$) based on experimental data of for SulA overexpression (32). In the simulation, doubling time ($\tau$) and the duration of C period ($C$) were assigned the average experimental values ($\overline{\tau}$=34.6 min, $\overline{C}$=43 min). By fitting experimental data on cell size as a function of $D$, we have $\alpha' = 0.55\mu m^3/s$. Other parameters are present in the Supporting Text. (A) Regular oscillations of origin number, genome content, total DnaA number, and cell size coupled with cell cycles were obtained in the simulated steady states with different $D$. (B-C) From the simulation, origin number and genome content per cell increase with C period, exactly following the theoretical formula and in line with experimental data. (D) The fitted cell size exponentially increases with $D$, just as the division growth law. (E) The predicted number of DnaA in each form exponentially increases with $D$. (F) The predicted concentration of DnaA in each form is constant. (G) The variables at initiation of DNA replication do not change with $D$. Each initiation variable was normalized by dividing the value at $D = 80$ min.

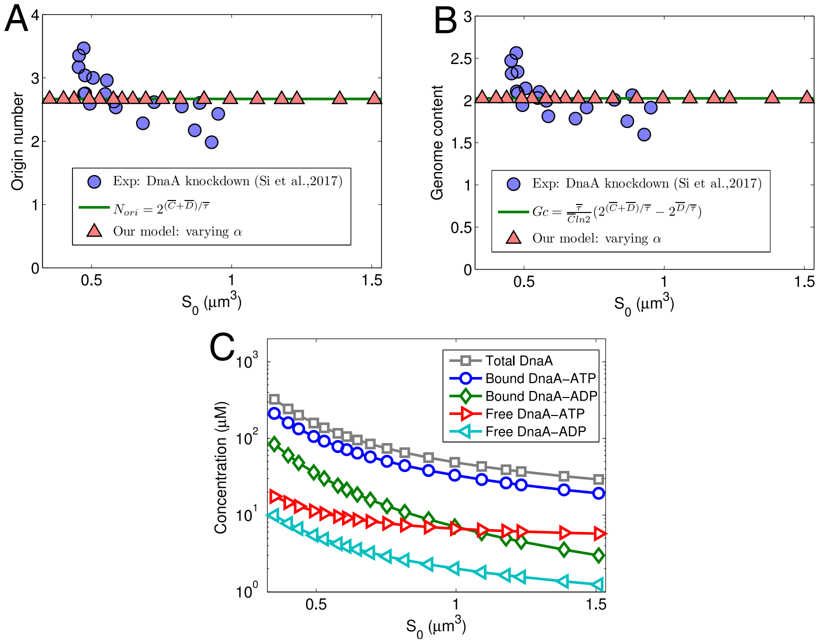


Figure S10: Our model quantifies the change in the origin number, genome content, and DnaA concentrations when initiation mass (i.e. $S_0$) is altered by changing the DnaA synthesis rate constant $\alpha$, based on experimental data of Si et al. (32) for DnaA knowndown (growth medium: MOPS glucose + 6 a.a.) The parameters used here are the same as those for Fig. 4 in the main text. (A, B) Origin number and genome content from the simulation are invariant with $S_0$, as theoretical formulas, but experimental ones tend to decrease with $S_0$. (C) The predicted concentration of DnaA in each form decreases with $S_0$.

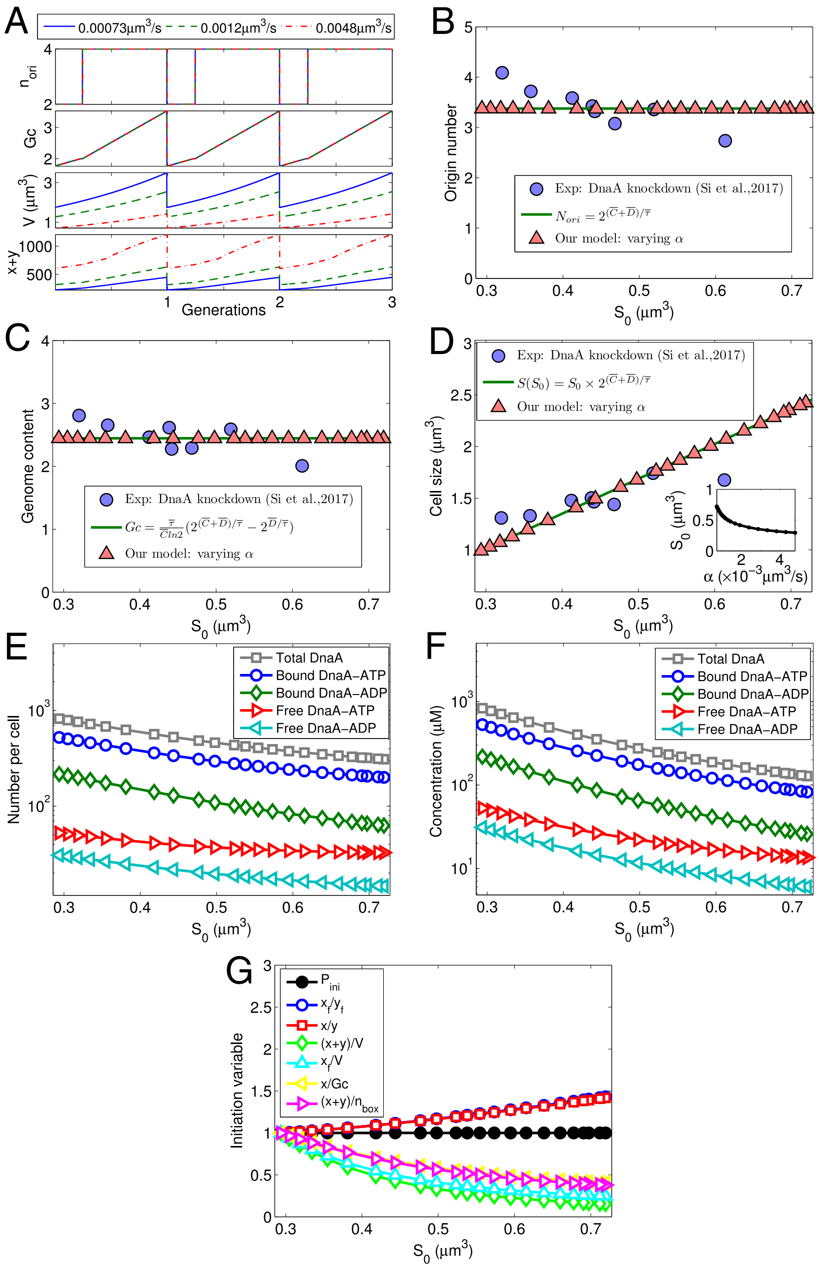


Figure S11: Our model quantifies changes in origin number, genome content, cell size, DnaA levels and initiation variables when initiation mass ($S_0$) is altered by changing DnaA synthesis rate constant $\alpha$ based on experimental data of Si et al. (32) for DnaA knowndown (growth medium: MOPS glucose + 6 a.a. + 0.2mM uracil). Experimental averages ($\overline{\tau}$= 43.2 min, $\overline{C}$= 42.5 min, $\overline{D}$=33.2 min) were assigned to doubling time and durations of C and D periods in the simulation. No fitting was done here. Values for other parameters are present in the Supporting Text. (A) Regular oscillations of origin number, genome content, total DnaA number, and cell size are coupled with cell cycles in simulated steady states with different values of $\alpha$. (B, C) The simulated origin number and genome content per cell are invariant with $S_0$, as theoretical formulas, but experimental ones tend to decrease with $S_0$. (D) The predicted cell size linearly increases with $S_0$, precisely following the growth law, while the predicted increase is a little faster than the experimental one. The inset plot shows $S_0$ decreases with $\alpha$. (E, F) The predicted number (concentration) of DnaA in each form decreases with $S_0$. (G) The initiation probability ($P_{ini}$) does not change with $S_0$ since the same threshold was used, $x_f/y_f$ and $x/y$ at initiation increase with $S_0$ similarly, while $(x+y)/V$, $x_f/V$, $x/Gc$ and $(x+y)/n_{box}$ at initiation all decrease with $S_0$.

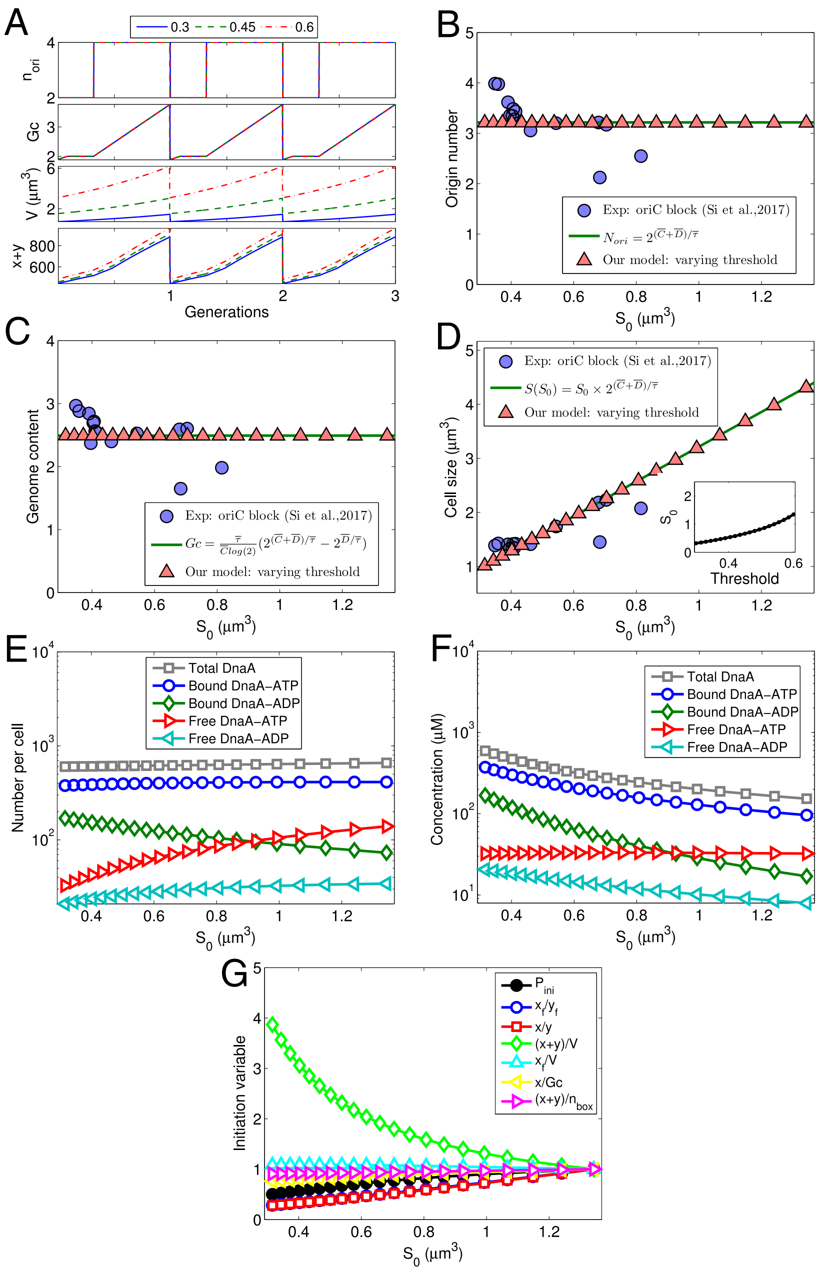


Figure S12: Our model quantifies changes in origin number, genome content, cell size, DnaA level and initiation variables when the initiation mass ($S_0$) is altered by changing the initiation threshold based on experimental data of (32) for *oriC* block. Experimental averages ($\overline{\tau}$= 43.6 min, $\overline{C}$=33.5 min, $\overline{D}$=39.9 min) were assigned to doubling time and durations of C and D periods in the simulation, respectively. No fitting was done here. Values for other parameters are present in the Supporting Text. (A) Regular oscillations of origin number, genome content, total DnaA number, and cell size are coupled with cell cycles in simulated steady states with different values of $\alpha$. (B, C) The simulated origin number and genome content per cell are invariant with $S_0$, as theoretical formulas, but experimental ones tend to decrease with $S_0$. (D) The predicted cell size linearly increases with $S_0$, precisely following the growth law, while the predicted increase is a little faster than the experimental one. The inset plot shows the increase in $S_0$ with the initiation threshold. (E) As $S_0$ increasing, numbers of total DnaA and bound DnaA-ATP increase slightly, numbers of free DnaA-ATP and free DnaA-ADP increase significantly, whereas the number of bound DnaA-ADP decreases. (F) The concentration of each form of DnaA decreases with $S_0$. (G) As $S_0$ increasing, at the initiation of DNA replication, $P_{ini}$ increases (resulting from the increase in initiation threshold), $x_f/y_f$, $x/y$, $x/Gc$, and $(x+y)/n_{box}$ increase differently from $P_{ini}$, whereas $(x+y)/V$ and $x_f/V$ decrease.

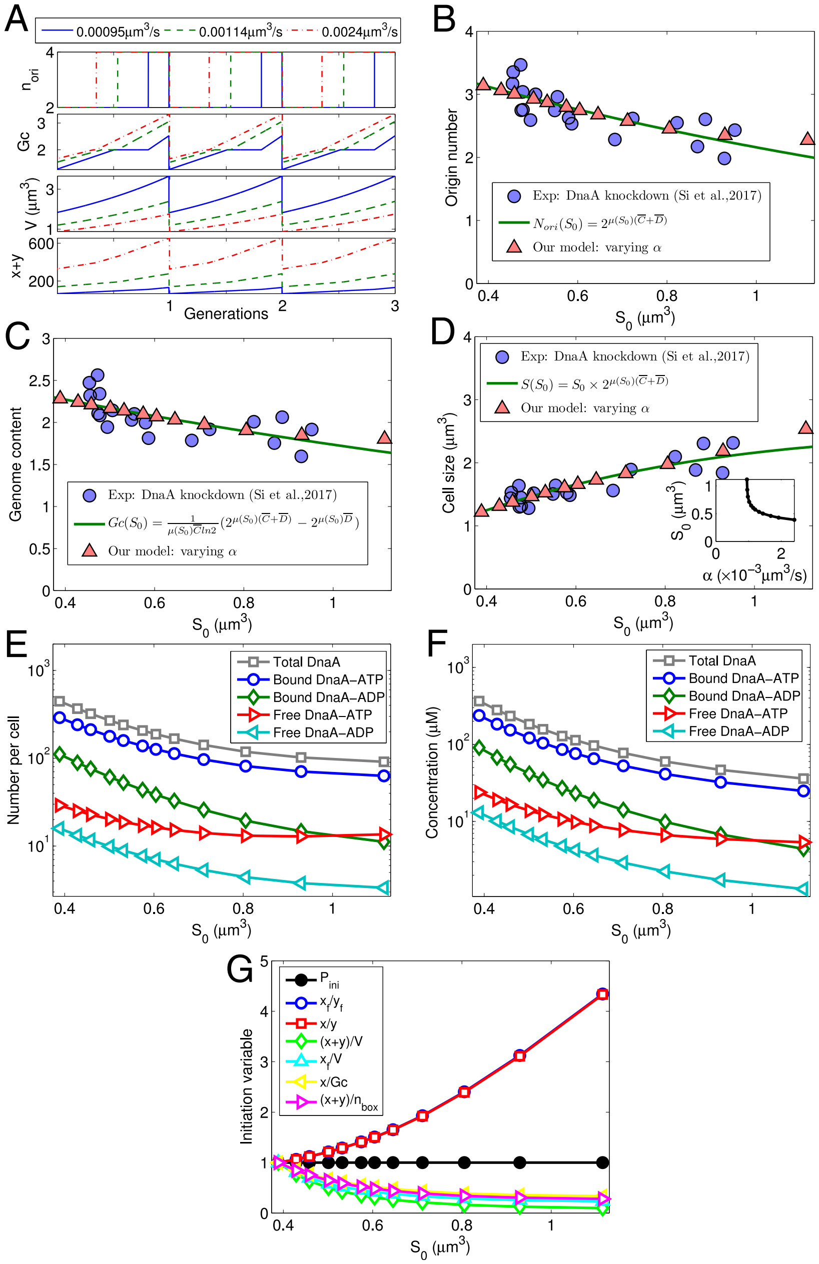

Figure S13: Our model quantifies the change in the origin number, genome content, and DnaA concentrations when initiation mass (i.e. $S_0$) and growth rate were altered by changing the DnaA synthesis rate constant $\alpha$ based on experimental data for DnaA knowndown (growth medium: MOPS glucose + 6 a.a) (32). From the experiment, the growth rate decreases slightly with initiation mass which can be fitted with a linear line $\mu = -0.61S_0 + 1.38$. With the same parameters for Fig. S10, we did one simulation run. With the simulated initiation mass, we produced the growth rate changing with $\alpha$ for a new simulation run. Three iterative updates were done for the results shown here. (A) At steady state, regular oscillations of origin number, genome content, DnaA number, and cell size are coupled with cell cycles with different values of $\alpha$. (B, C) Origin number and genome content from the simulation decreases with $S_0$, basically following the corrected formulas, and in agreement with experimental data. (D) The predicted cell size increases with $S_0$ following the corrected growth law and in agreement with experimental data. Inset plot presents unit cell size $S_0$ increases with $\alpha$. (E, F) The predicted number (concentration) of DnaA in each form decreases with $S_0$. (G) The initiation probability ($P_{ini}$) does not change with $S_0$ since the same initiation threshold was used, other initiation variables increase ($x/y$ and $x_f/y_f$) or decrease ($(x+y)/V$, $x_f/V$, $x/Gc$ and $(x+y)/n_{box}$) with $S_0$. Each initiation variable was divided by the value at $\alpha = 2.4 \times 10^{-3} \mu m^3/s$ ($S_0 = 0.35 \mu m^3$).

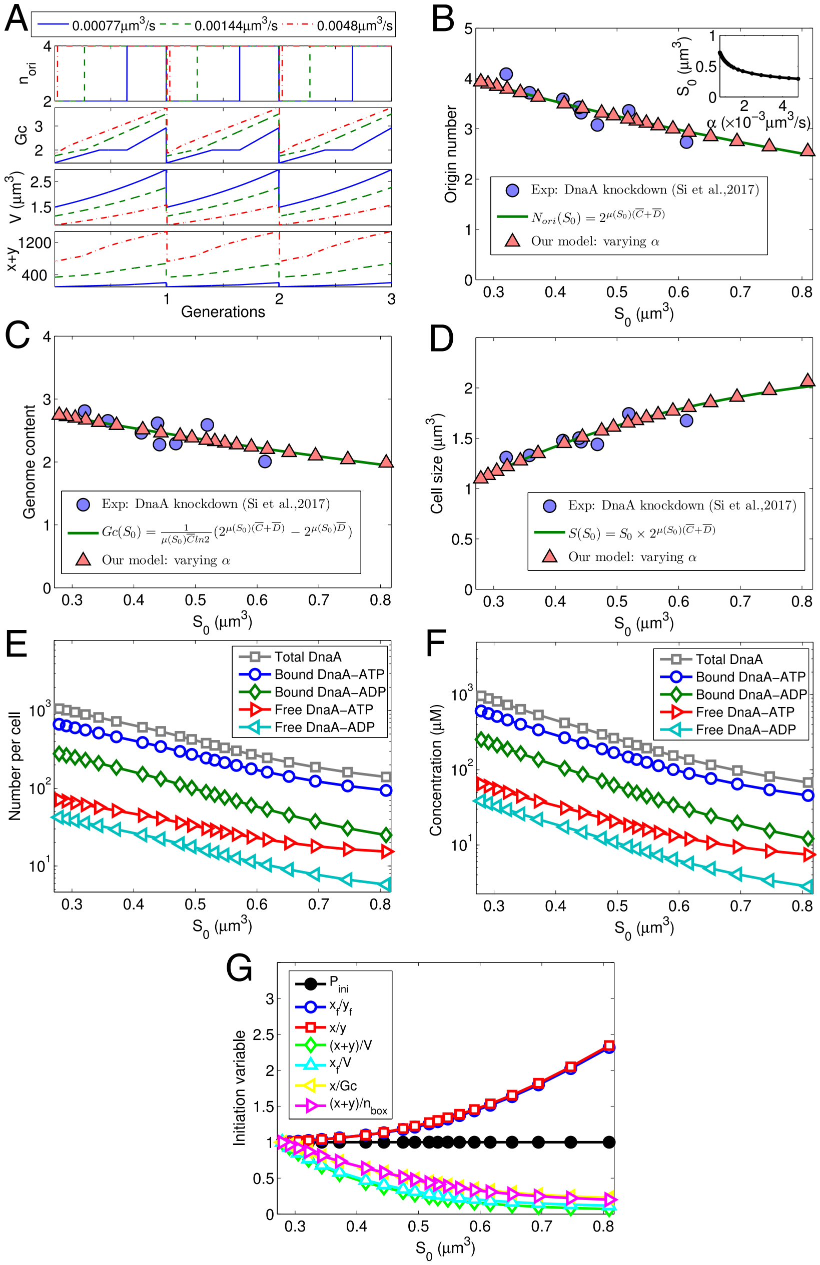

Figure S14: Our model quantifies changes in origin number, genome content, cell size, DnaA levels and initiation variables when initiation mass ($S_0$) and growth rate were altered by changing the DnaA synthesis rate constant $\alpha$ based on experimental data for DnaA knowndown (growth medium: MOPS glucose + 6 a.a. + 0.2mM uracil) (32). From the experiment, the growth rate decreases slightly with initiation mass which can be fitted with a linear line $\mu = -0.99S_0 + 1.84$. With the same parameters for Fig. S11, we did one simulation run. With the simulated initiation mass, we produced the growth rate changing with $\alpha$ for a new simulation run. Three iterative updates were done for the results shown here. (A) At steady state, regular oscillations of origin number, genome content, DnaA number, and cell size are coupled with cell cycles with different values of $\alpha$. (B, C) Origin number and genome content from the simulation decrease with $S_0$, as the corrected theoretical formulas and in agreement with experimental data. The inset plot shows $S_0$ decreases with $\alpha$. (D) The predicted cell size increases with $S_0$, following the corrected growth law and in agreement with experimental data. (E, F) The predicted number (concentration) of DnaA in each form decreases with $S_0$. (G) At the initiation of DNA replication, the initiation probability ($P_{ini}$) does not change with $S_0$ since the same initiation threshold was used, other initiation variables increase ($x/y$ and $x_f/y_f$) or decrease ($(x+y)/V$, $x_f/V$, $x/Gc$ and $(x+y)/n_{box}$) with $S_0$. Each initiation variable was divided by the value at $\alpha = 4.8 \times 10^{-3} \mu m^3/s$ ($S_0 = 0.28\ \mu m^3$)

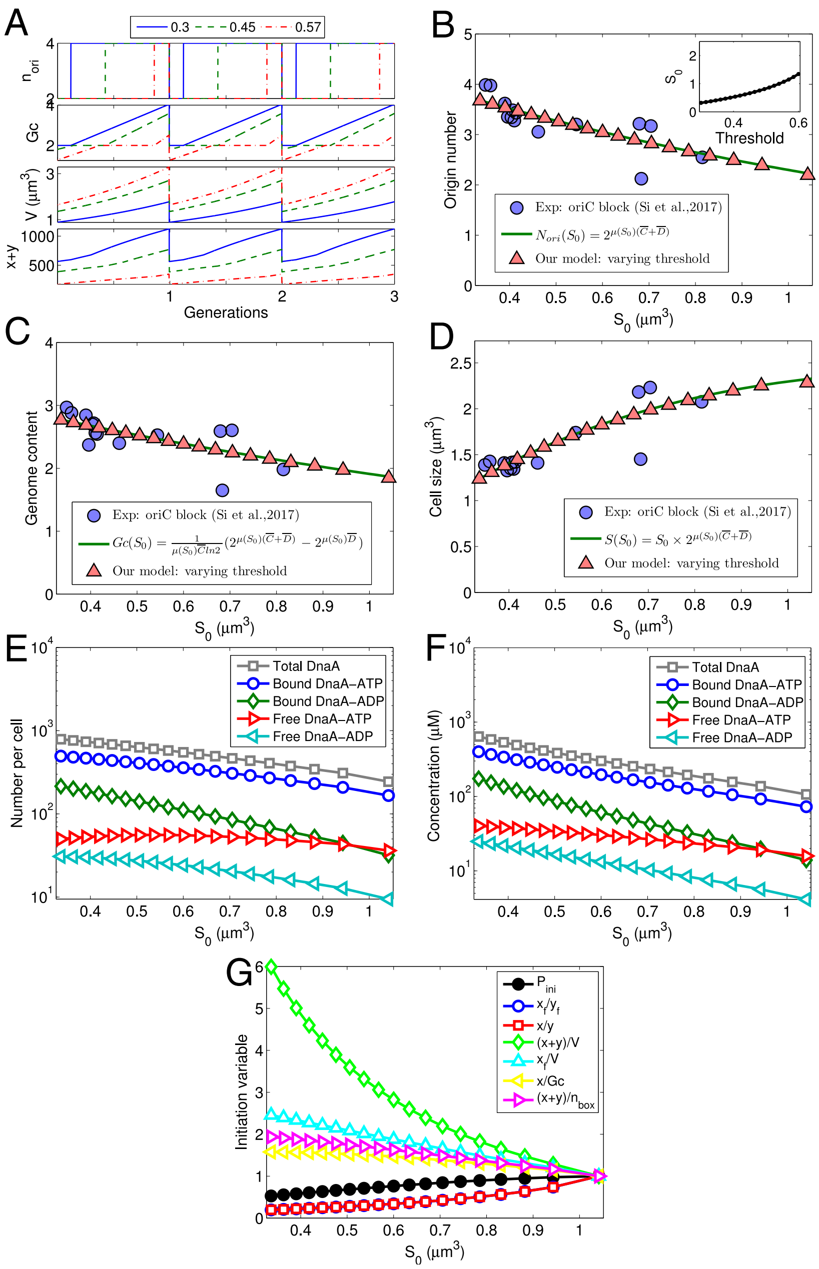


Figure S15: Our model quantifies changes in origin number, genome content, cell size, DnaA levels and initiation variables when initiation mass ($S_0$) and growth rate were altered by changing initiation threshold based on experimental data for *oriC* block ((32)). From the experiment, the growth rate decreases slightly with initiation mass which can be fitted with a linear line $\mu = -0.84S_0 + 1.82$. With the same parameters for Fig. S12, we did one simulation run. With the simulated initiation mass, we produced the growth rate changing with the initiation threshold for a new simulation run. Three iterative updates were done for the results shown here. Relevant experimental data for origin number, genome content, and cell size as a function of $S_0$ were used for comparison. (A) Regular oscillations of origin number, genome content, total DnaA number, and cell size are coupled with cell cycles in steady states with different initiation thresholds. (B, C) Origin number and genome content from the simulation decrease with $S_0$, as the corrected theoretical formulas and in agreement with experimental data. The inset plot shows the increase in $S_0$ with initiation threshold. (D) The predicted cell size increases with $S_0$, following the corrected growth law and in agreement with experimental data. (E, F) The predicted number (concentration) of DnaA in each form decreases with $S_0$ (Free DnaA-ATP increases a little when $S_0 <\sim 0.6$). (G) As $S_0$ increasing, at initiation of DNA replication, $P_{ini}$ increases (resulting from the increase in initiation threshold), $x_f/y_f$ and $x/y$ increase differently from $P_{ini}$, whereas $x/Gc$, $(x+y)/n_{box}$, $(x+y)/V$, and $x_f/V$ decrease. Each initiation variable was divided by the value at initiation threshold=0.57 ($S_0 = 1.04\mu m^3$)

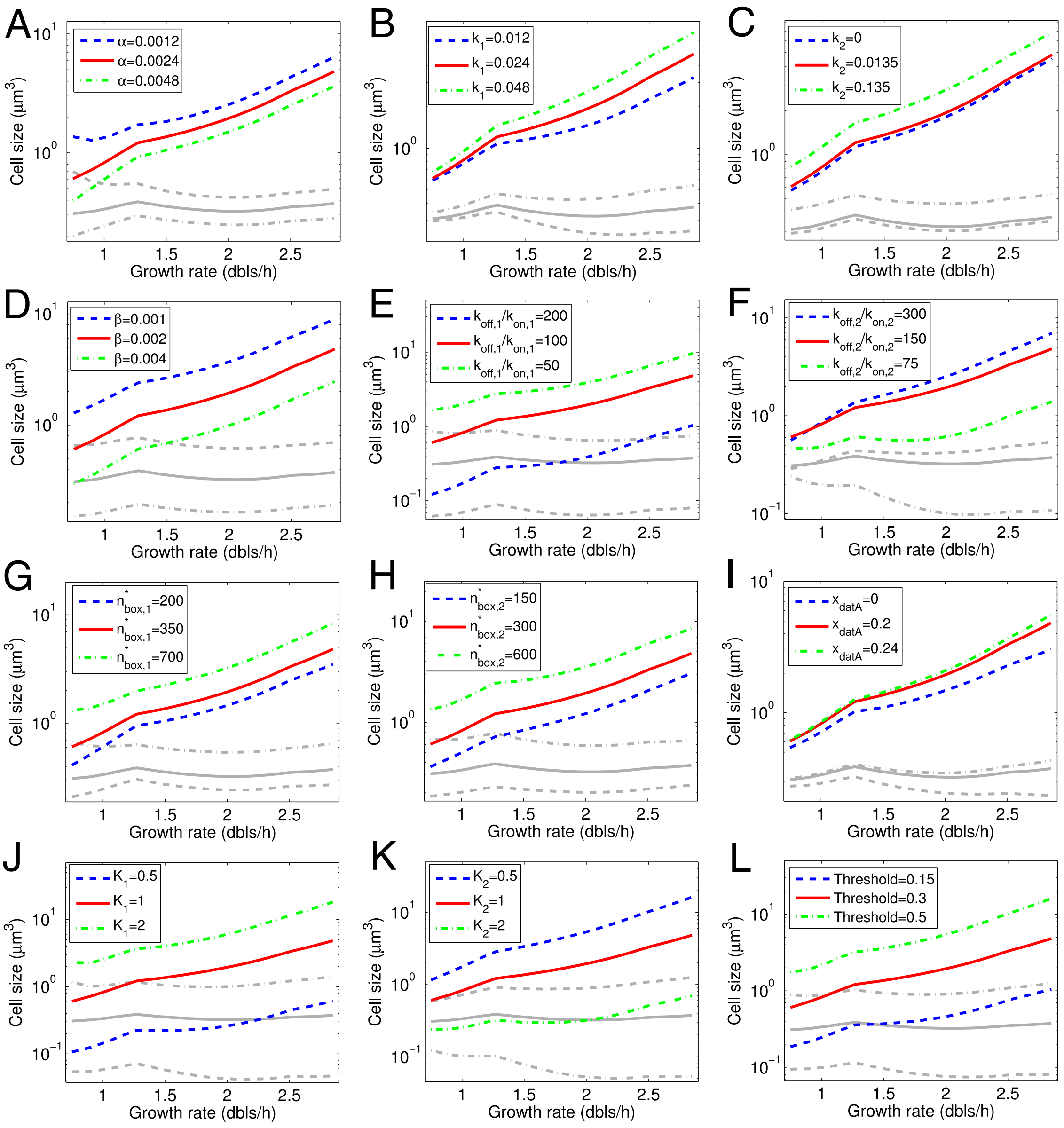


Figure S16: The dependence of cell size as a function of growth rate on each parameter in our model. The parameters shown by red lines are the same as those for Fig. 2 in the main text. The units of the perturbed parameters in the panels (A)-(L) are $\mu m^3/s$, $\mu m^3$, $\mu m^3$, $s^{-1}$, $\mu m^{-3}$, $\mu m^{-3}$, 1, 1, 1, $\mu m^{-3}$, $\mu m^{-3}$, and 1 respectively.

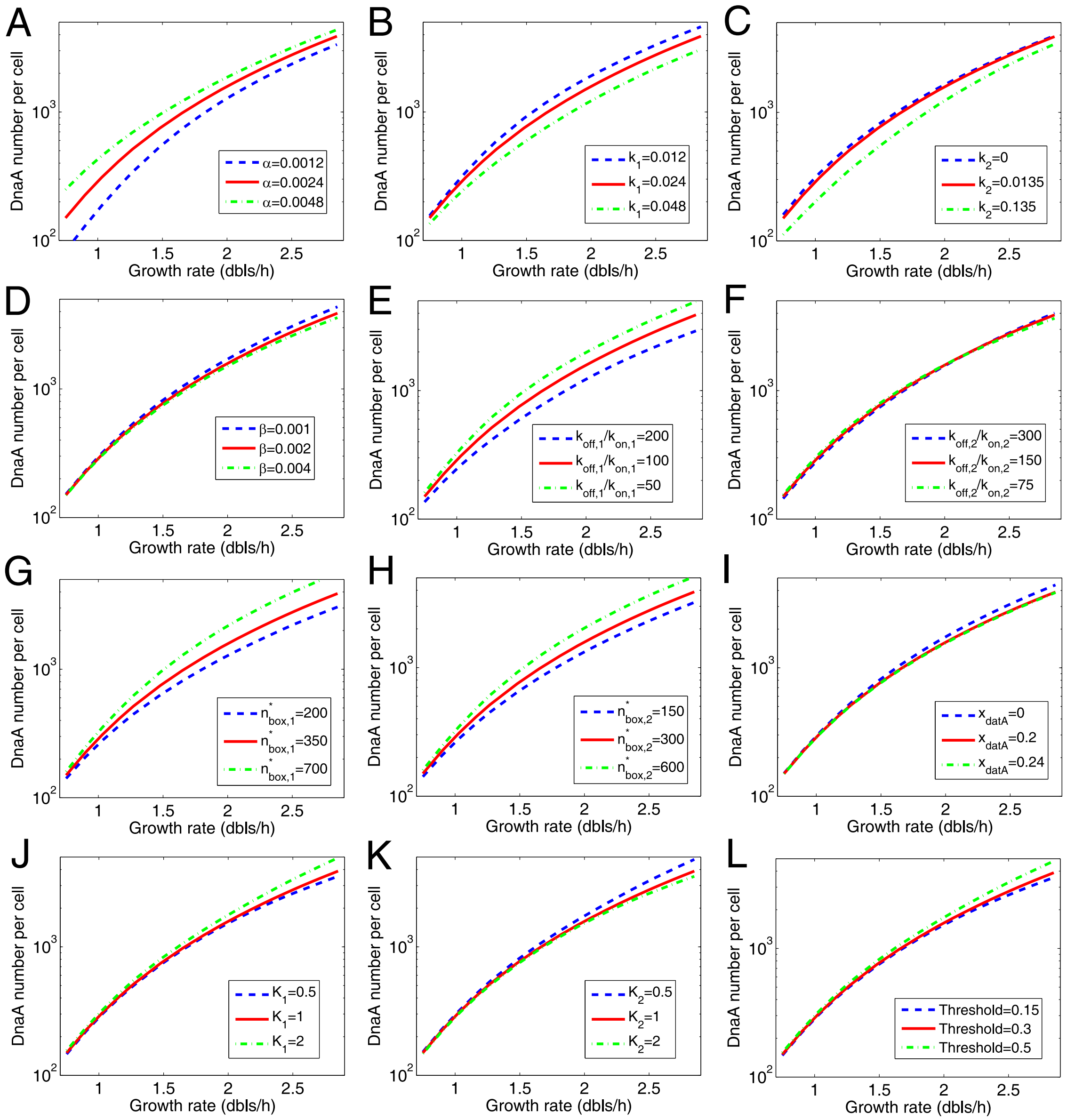


Figure S17: The dependence of DnaA number per cell as a function of growth rate on each parameter in our model. The parameters shown by red lines are the same as that for Fig. 2 in the main text. The units of the parameters shown in the panels (A)-(L) are $\mu m^3/s$, $\mu m^3$, $\mu m^3$, $s^{-1}$, $\mu m^{-3}$, $\mu m^{-3}$, 1, 1, 1, $\mu m^{-3}$, $\mu m^{-3}$, and 1 respectively.

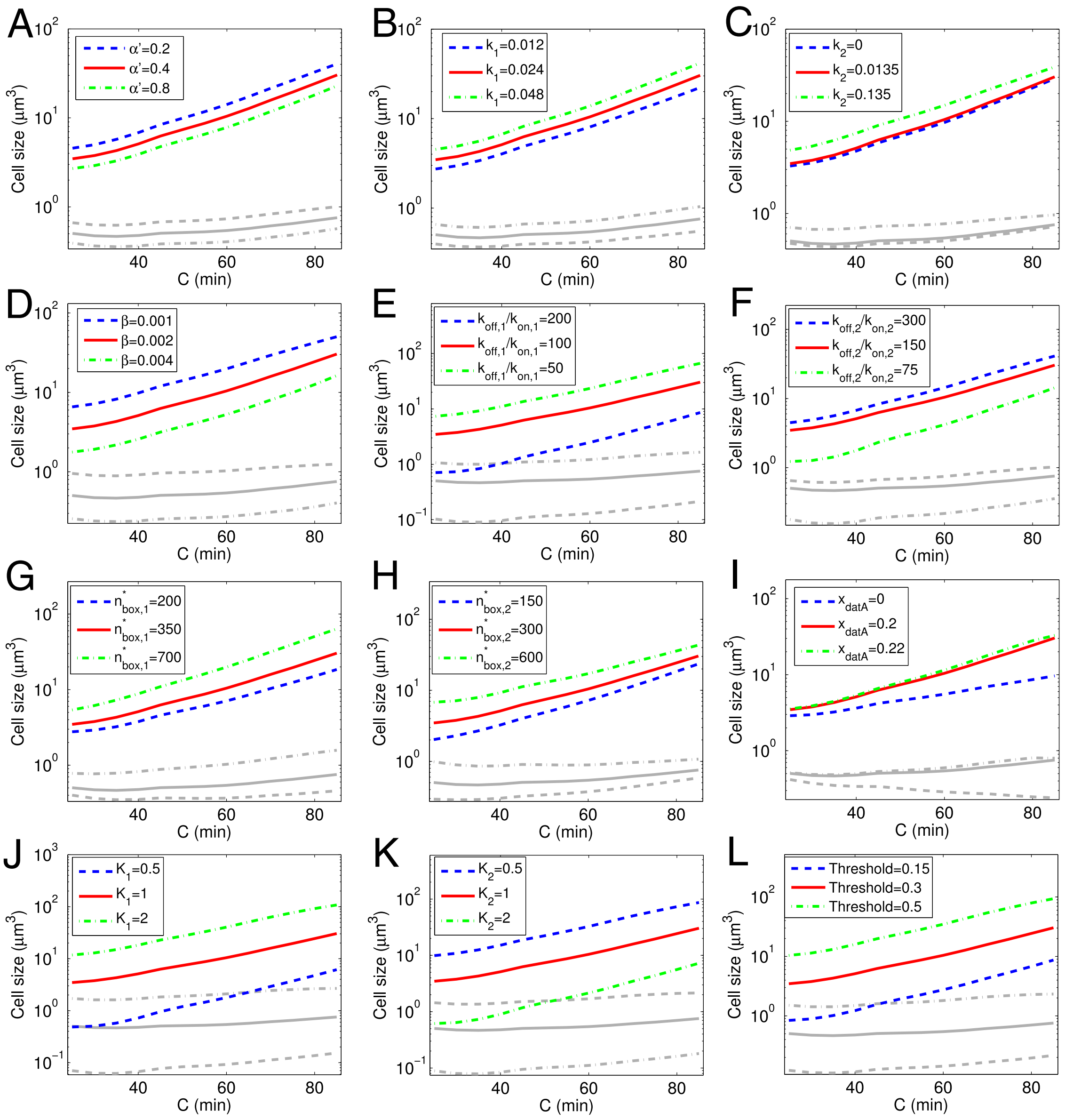


Figure S18: The dependence of cell size as a function of the duration of C period on each parameter in our model. The parameters shown by red lines are the same as those for Fig. 3 in the main text. The units of the parameters shown in the panels (A)-(L) are $\mu m^3/s$, $\mu m^3$, $\mu m^3$, $s^{-1}$, $\mu m^{-3}$, $\mu m^{-3}$, 1, 1, 1, $\mu m^{-3}$, $\mu m^{-3}$, and 1 respectively.

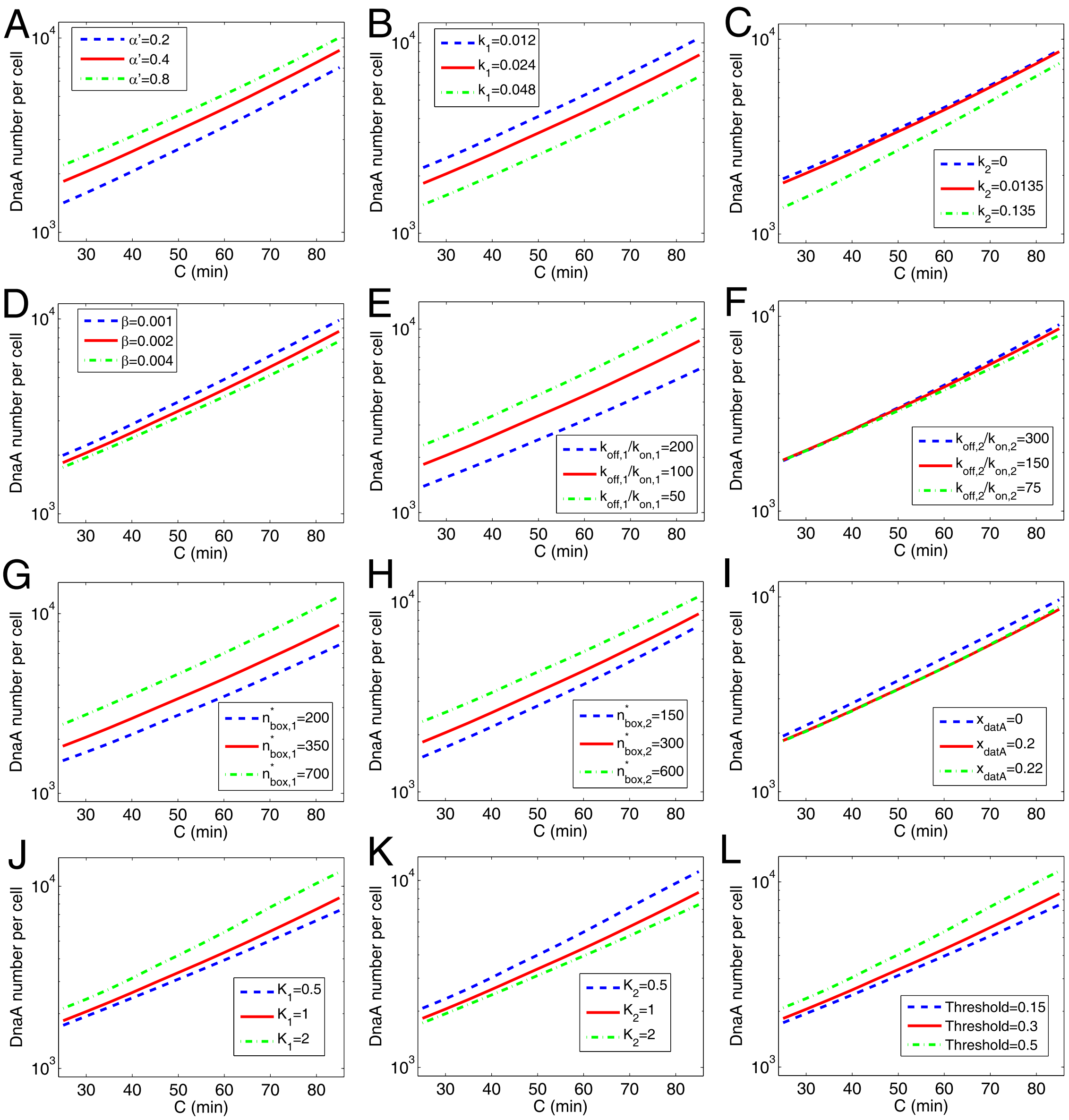


Figure S19: The dependence of DnaA number per cell as a function of the duration of C period on each parameter in our model. The parameters shown by red lines are the same as those for Fig. 3 in the main text. The units of the parameters shown in the panels (A)-(L) are $\mu m^3/s$, $\mu m^3$, $\mu m^3$, $s^{-1}$, $\mu m^{-3}$, $\mu m^{-3}$, 1, 1, 1, $\mu m^{-3}$, $\mu m^{-3}$, and 1 respectively.

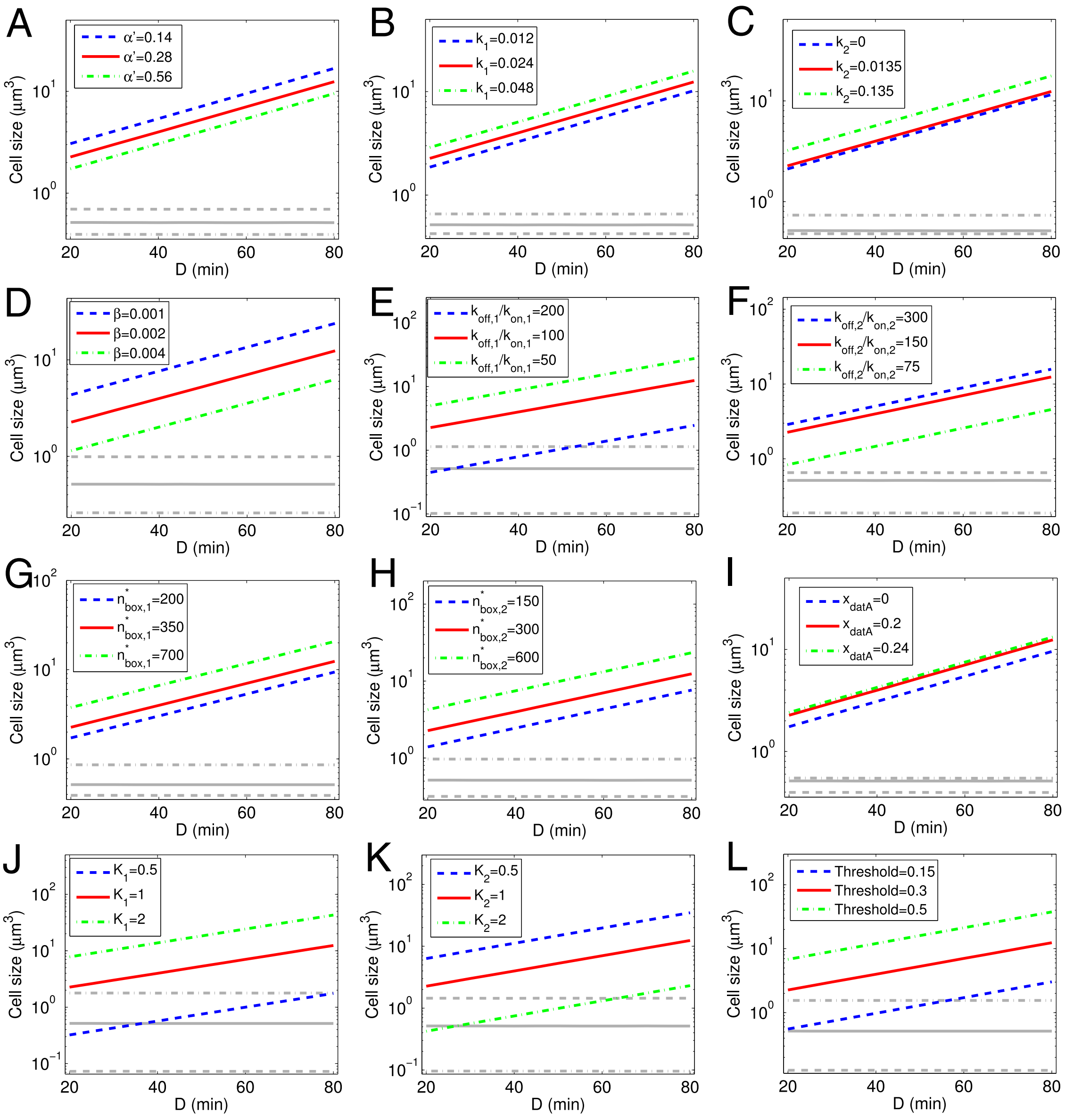


Figure S20: The dependence of cell size as a function of the duration of D period on each parameter in our model. The parameters shown by red lines are the same as those for Fig. 4 in the main text. The units of the parameters shown in the panels (A)-(L) are $\mu m^3/s$, $\mu m^3$, $\mu m^3$, $s^{-1}$, $\mu m^{-3}$, $\mu m^{-3}$, 1, 1, 1, $\mu m^{-3}$, $\mu m^{-3}$, and 1 respectively.

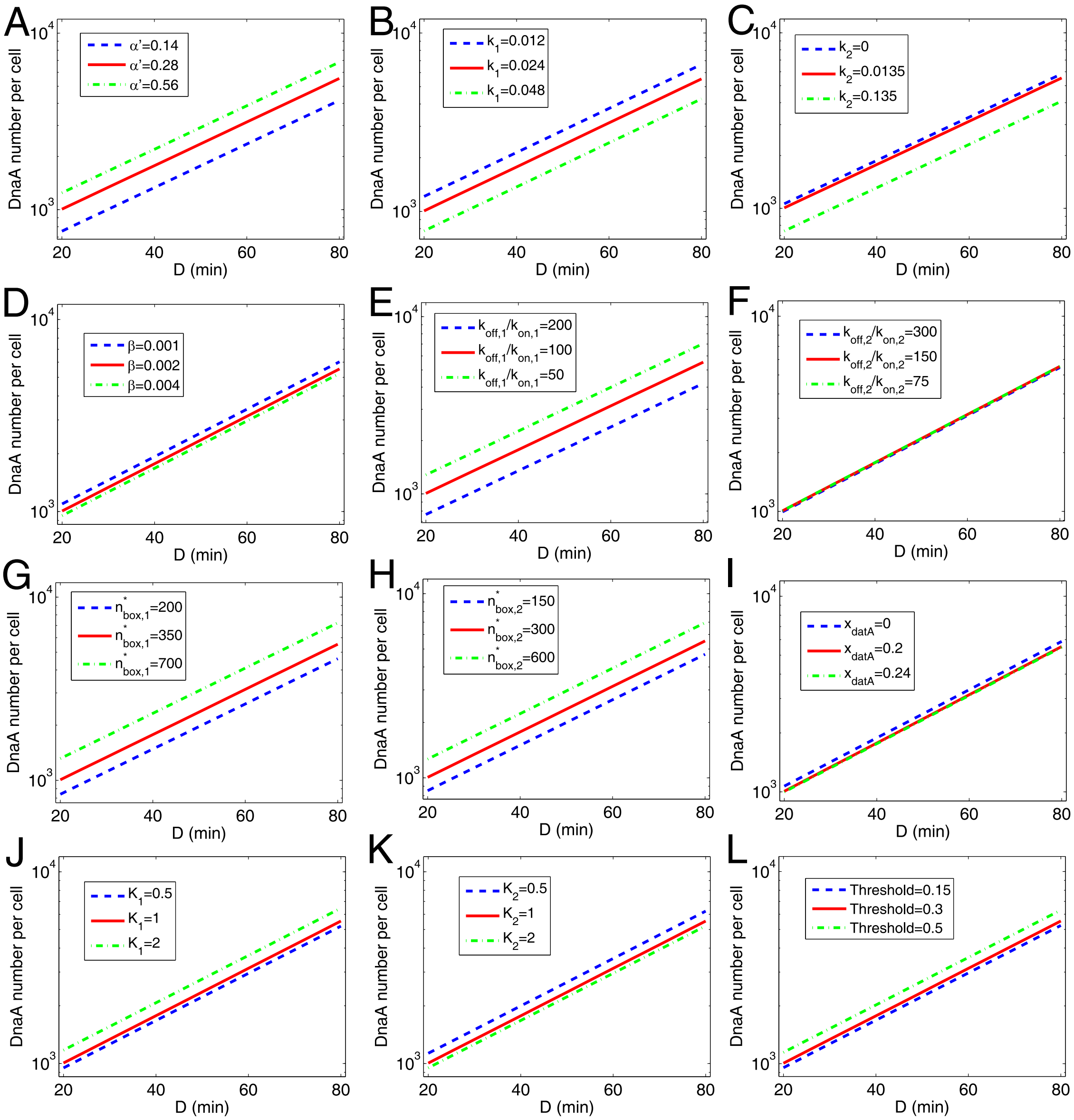


Figure S21: The dependence of DnaA number per cell as a function of the duration of D period on each parameter in our model. The parameters shown by red lines are the same as those for Fig. 4 in the main text. The units of the parameters shown in the panels (A)-(L) are $\mu m^3/s$, $\mu m^3$, $\mu m^3$, $s^{-1}$, $\mu m^{-3}$, $\mu m^{-3}$, 1, 1, 1, $\mu m^{-3}$, $\mu m^{-3}$, and 1 respectively.

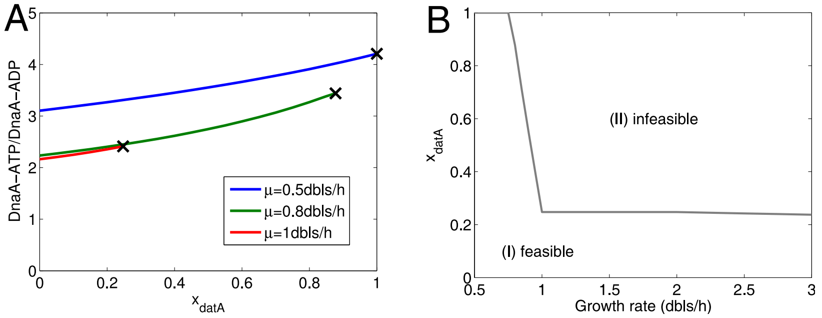


Figure S22: Our model predicts the effect of the chromosomal position of *datA*. (A) The ratio of DnaA-ATP to DnaA-ADP increases with the relative distance from *datA* to *oriC* ($x_{datA}$), however, no regular oscillation is obtained (with $10^7$ steps and each step = 1 s) when $x_{datA}$ is bigger than the point at the cross. The lines in different colors show the dependence of cell size on $x_{datA}$ at different growth rates ($\mu$). (B) At high growth rates, a long distance from *datA* to *oriC* is infeasible, i.e. no regular oscillation emerges in the simulation (with $10^7$ steps and each step = 1 s). This forms two regimes on the plot of $x_{datA}$ versus growth rate: a feasible one where a regular oscillation emerges in the simulation and an infeasible one where no regular oscillation emerges. The grey line shows the border (corresponding to the crosses in A) between the two regimes.

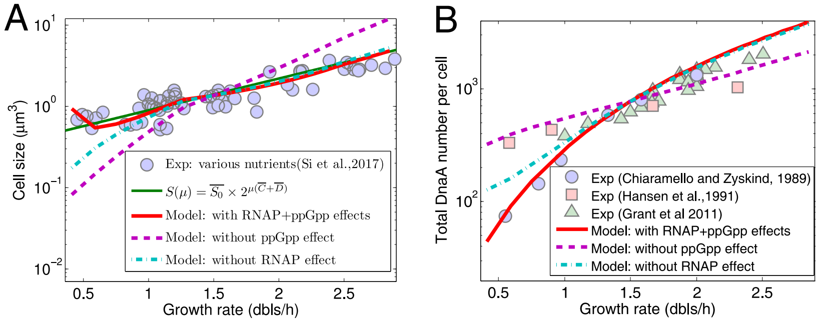


Figure S23: The result changes significantly if we do not consider the change in RNAP or ppGpp with the growth rate (by replacing Eq. 7 or Eq.8 in the main text with $[RNAP]_f = [RNAP]_{f,0} \exp(-\mu_r/1.5\ dbl/h)$ or $[ppGpp] = [ppGpp]_0 \exp(-1.5\ dbl/h/\mu_p)$). (A) Without RNAP or ppGpp effect (no change in RNAP or ppGpp), cell size from the simulation changes with the growth rate faster. (A) Without RNAP or ppGpp effect (no change in RNAP or ppGpp), DnaA number per cell changes with the growth rate more slowly.

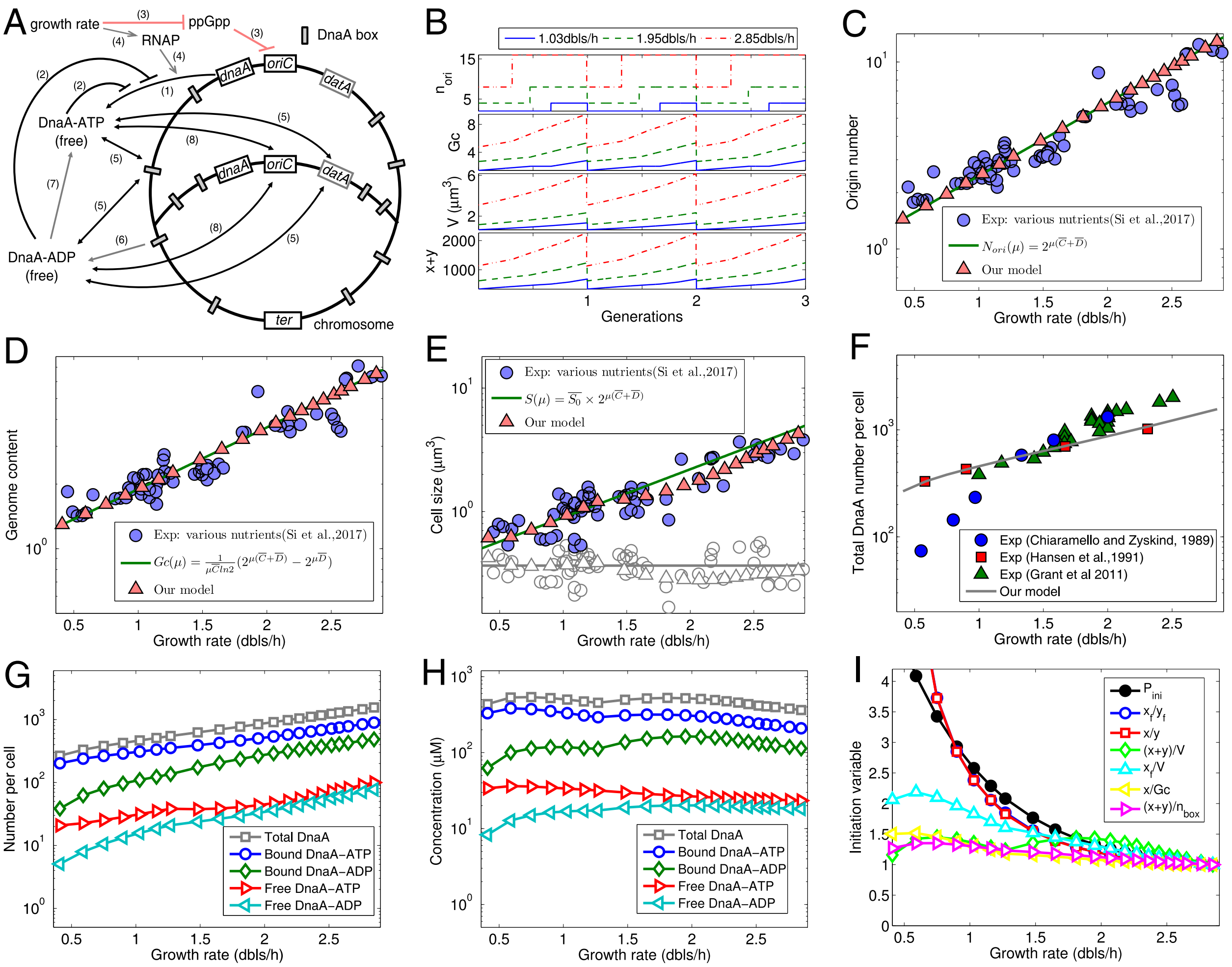

Figure S24: Possible effects of the global regulator ppGpp inhibiting *oriC* independently of DnaA synthesis. (A) Regulatory processes for DnaA (adapted from (1, 42, 43)). The processes (1)-(2) and (4)-(8) are the same as in Figure 1 B in the main text. The process (3) (red line) indicates that (p)ppGpp, whose concentration decreases with growth rate, reduces the initiation probability of *oriC* may by affecting supercoiling instead of inhibiting DnaA synthesis. In the simulation, the inhibition of ppGpp on DnaA synthesis was removed by replacing the inhibition function $G([ppGpp])$ by its value at growth rate of 1.33 doublings/hour (i.e. $G([ppGpp]) \equiv 0.0625$), the initiation threshold $c$ was expressed as a function of $[ppGpp]$, i.e., $c = c_0[(ppGpp/K_g)^m+1]$, where $K_g = 24pmol/OD460$, $m = 1.6$, $c_0 = 0.13$ obtained by fitting the data of cell size. Other parameters are the same as used in Figure 1 B. (B) At steady state with different growth rates, regular oscillations in total DnaA number, origin number, genome content, and cell size are coupled to the cell cycle. (C-D) The predicted number of origins or genome content increases exponentially with the growth rate, exactly following the theoretical formula. (E) By fitting experimental data, cell size increases with growth rate, roughly following the nutrient growth law. Unit cell sizes (initiation masses) from the model (grey open triangles) and experiment (grey open circles) are located around the line $\overline{S_0}$ (grey). (F) The predicted total DnaA number per cell increases with growth rate (increasing around 3 times from 0.5 dbls/h to 2.5 dbls/h), in agreement with experimental data of Hansen et al. (44). The predicted increase is significantly slower than experimental data of Chiaramello and Zyskind (45) and Grant et al. (46). (G) The predicted number of each form of DnaA per cell increases with growth rate. (H) The predicted concentration of each form of DnaA changes with growth rate slightly. (I) $P_{ini}$ at the initiation decreases with growth rate since ppGpp and the initiation threshold both decrease with growth rate. Other variables at the initiation all decrease with growth rate: $x_f/y_f$ and $x/y$ decrease more rapidly than $P_{ini}$, whereas others decrease more slowly. Each initiation variable was divided by the value at a growth rate of 2.85 dbls/h.

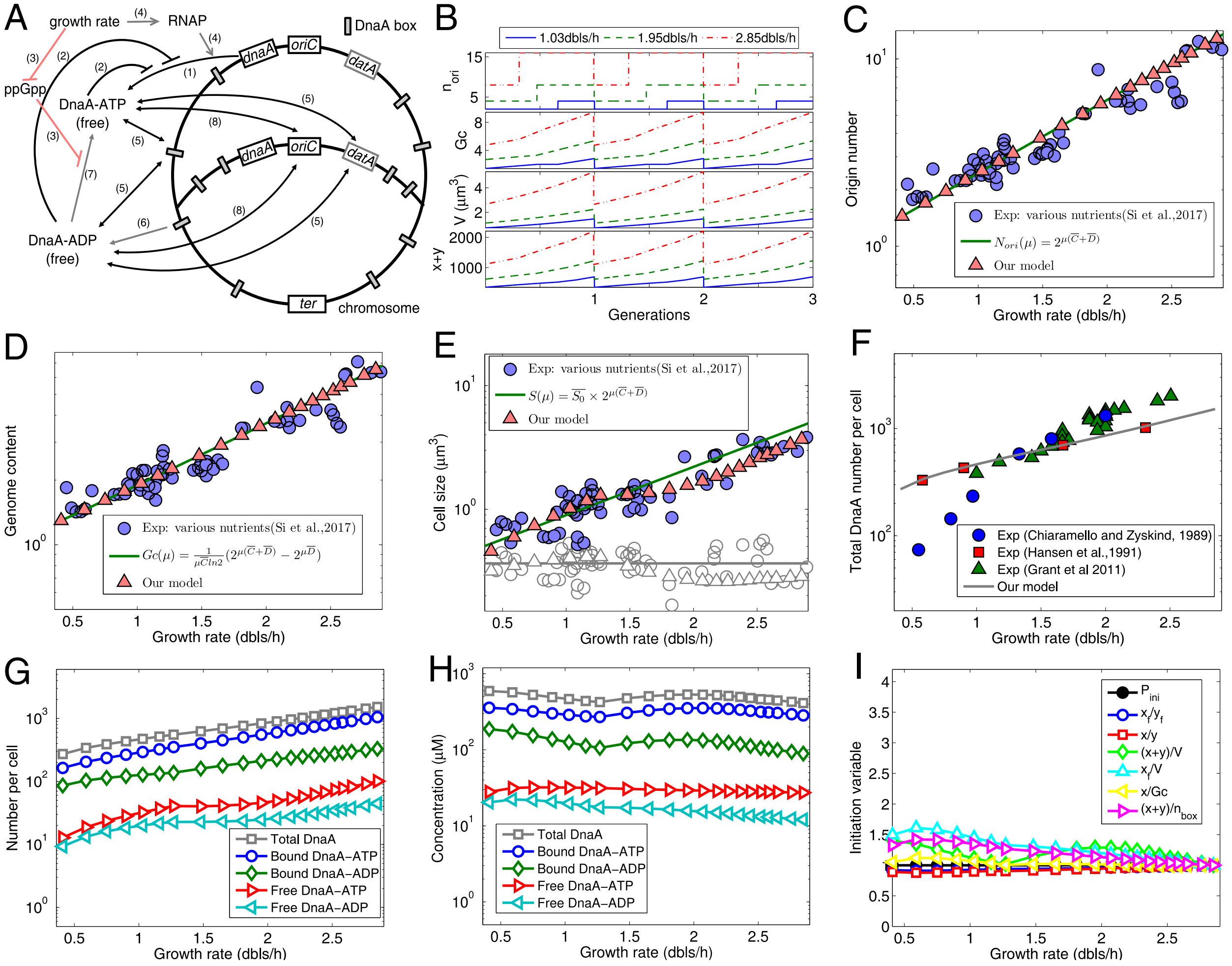


Figure S25: Possible effects of the global regulator ppGpp reducing the reactivation rate of DnaA-ADP. (A) Regulatory processes for DnaA (adapted from (1, 42, 43)). The processes (1)-(2) and (4)-(8) are the same as in Figure 1 B in the main text. The process (3) (red line) indicates that (p)ppGpp, whose concentration decreases with growth rate, reduces the reactivation rate of DnaA-ADP may by inhibiting Fis which activates DARS. In the simulation, the inhibition of ppGpp on DnaA synthesis was removed by replacing the inhibition function $G([ppGpp])$ by its value at growth rate of 1.33 doublings/hour (i.e. $G([ppGpp]) \equiv 0.0625$), and $\beta$ was expressed as a function of $[ppGpp]$, i.e., $\beta = \beta_0/[(ppGpp/K_g)^m + 1]$, where $K_g = 15pmol/OD460$, $m = 2.1$, and $\beta_0 = 9 \times 10^{-3}$, obtained by fitting the data of cell size. Other parameters are the same as used in Figure 1 B. (B) At steady state with different growth rates, regular oscillations in total DnaA number, origin number, genome content, and cell size are coupled to the cell cycle. (C-D) The predicted number of origins or genome content increases exponentially with the growth rate, exactly following the theoretical formula. (E) By fitting experimental data, cell size increases with growth rate, roughly following the nutrient growth law. Unit cell sizes (initiation masses) from the model (grey open triangles) and experiment (grey open circles) are located around the line $\overline{S_0}$ (grey). (F) The predicted total DnaA number per cell increases with growth rate (increasing around 3 times from 0.5 dbls/h to 2.5 dbls/h), in agreement with experimental data of Hansen et al. (44). The predicted increase is significantly slower than experimental data of Chiaramello and Zyskind (45) and Grant et al. (46). (G) The predicted number of each form of DnaA per cell increases with growth rate. (H) The predicted concentration of each form of DnaA changes with growth rate slightly. (I) $P_{ini}$ at the initiation of DNA replication is constant since the same initiation threshold was used at different growth rates. At the initiation, $x_f/y_f$ and $x/y$ increase slowly, $x_f/V$, $x/Gc$, and $(x+y)/n_{box}$ basically decrease more or less, whereas $(x+y)/V$ fluctuates with growth rate. Each initiation variable was divided by the value at a growth rate of 2.85 dbls/h.
